# Quantifying microbial associations of dissolved organic matter under global change

**DOI:** 10.1101/2021.08.12.456177

**Authors:** Ang Hu, Mira Choi, Andrew J. Tanentzap, Jinfu Liu, Kyoung-Soon Jang, Jay T. Lennon, Yongqin Liu, Janne Soininen, Xiancai Lu, Yunlin Zhang, Ji Shen, Jianjun Wang

## Abstract

Microbes play a critical role in regulating the size, composition, and turnover of dissolved organic matter (DOM), which is one of the largest pools of carbon in aquatic ecosystems. Global change may alter DOM-microbe associations with implications for biogeochemical cycles, although disentangling these complex interactions remains a major challenge. Here we develop a framework called Energy-Diversity-Trait integrative Analysis (EDTiA) to examine the associations between DOM and bacteria along temperature and nutrient gradients in a manipulative field experiment on mountainsides in contrasting subarctic and subtropical climates. In both study regions, the chemical composition of DOM correlated with bacterial communities, and was primarily controlled by nutrients and to a lesser degree by temperature. At a molecular-level, DOM-bacteria associations depended strongly on the molecular traits of DOM, with negative associations indicative of decomposition as molecules are more biolabile. Using bipartite networks, we further demonstrated that negative associations were more specialized than positive associations indicative of DOM production. Nutrient enrichment promoted specialization of positive associations, but decreased specialization of negative associations particularly at warmer temperatures in subtropical climate. These global change drivers influenced specialization of negative associations most strongly via molecular traits, while both molecular traits and bacterial diversity similarly affected positive associations. Together, our framework provides a quantitative approach to understand DOM-microbe associations and wider carbon cycling across scales under global change.

## Introduction

Dissolved organic matter (DOM), one of the largest pools of carbon in aquatic ecosystems ^1^, is intimately interlinked with the metabolic processes of complex microbial communities ^2^. Microbial consortia generate “chemodiversity” in the DOM pool by degrading larger molecules into smaller molecules and by synthesizing more refractory compounds from labile substrates ^3^. These basic processes together lead to the emergence of molecular traits of DOM including chemical structure, stoichiometry, oxidation state, and bioavailability ^4-6^ that directly determine its environmental persistence ^7, 8^. DOM, as a carbon source for microbial metabolism, also influences the diversity, structure, and functioning of microbial communities via decomposition and biosynthetic processes ^9-13^. The resulting resource-consumer relationships can now be characterised in both aquatic^14-16^ and terrestrial ^17^ ecosystems owing to recent advances in ultrahigh-resolution mass spectrometry and high-throughput sequencing. Despite the availability of these technologies, little is known about how DOM-microbe associations can be quantified in nature, and are interactively and independently affected by global change drivers, such as elevated temperatures and eutrophication.

The effects of global change on DOM-microbe associations can be viewed through three proximal controls (Fig. 1). First, energy supply, such as primary productivity, represents the major source of DOM that supports microbial metabolism ^18-20^. In particular, elevated temperature and nutrient inputs can stimulate primary productivity in ways that influences the composition and availability of organic matter ^21, 22^, but this process also indirectly influence DOM-microbe associations by controlling their diversity and traits ^22-25^. Second, diversity can generally beget diversity. For example, an increase in the diversity of DOM promotes microbial diversity, and vice versa ^15^, which should be reflected as signatures in resource-consumer relationships. Such patterns may arise because resource diversity promotes microbial specialization during biochemical transformations by creating more unique resource niches for consumers to partition ^26, 27^. Likewise, higher microbial diversity provides more metabolic pathways to decompose and produce molecules, which influences the vulnerability of DOM to degradation ^3^. Third, DOM-microbe associations depend on the molecular traits of DOM, such as its bioavailability, measured with H/C ratios of individual molecules ^28^, and microbial traits like life history (i.e., *r*-versus *K*-selection) ^29^ and resource acquisition (i.e., generalists versus specialists) ^27^.

**Figure 1.**
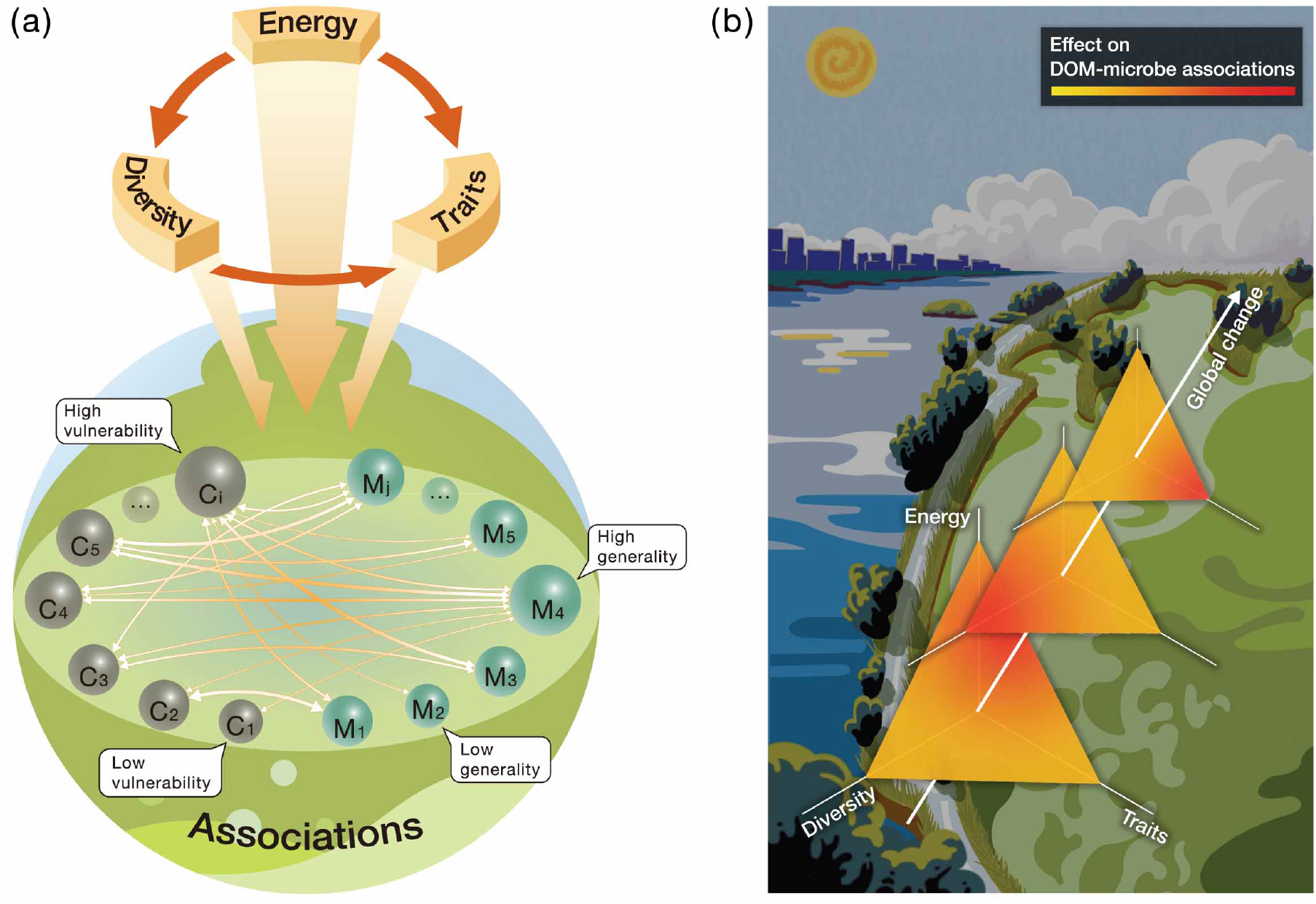
A framework for studying the effects of global change on DOM-microbe associations. (a) DOM-microbe associations affected by the three proximal controls, namely energy supply and both the diversity and traits of DOM and microbes. The relationships among the three controls and their influences on the associations are shown with single-sided arrows. The DOM-microbe associations, indicated by double-sided arrows, are measured by bipartite interactions between DOM molecules (circles C_1_-C_i_) and microbial species (circles M_1_-M_j_). The size of circles indicates the abundance of DOM molecules or microbial species, and the width of arrows is the magnitude of associations. Commonly used indices summarise the specialization of individual molecule i and microbial species j, such as *d*’ for DOM and microbes, which describes the levels of “vulnerability” of DOM molecules and “generality” of microbial species, respectively. (b) Conceptual framework for understanding DOM-microbe associations under distal drivers such as global change via the three proximal drivers. For better 3D visualization, the sizes of triangles decrease towards the top-right, and the color changes towards different corners of the triangles represent variations in the relative importance of different proximal drivers under a global change scenario. The background depicts the primary motivation of this study in examining distal drivers of climate change and eutrophication in Taihu Lake, China. The left and right waters indicate clean and cyanobacteria-dominated lake states, respectively, and are separated by a road having the shapes of western lakeshore and northern Zhushan and Meiliang Bays of Taihu Lake. We setup field microcosms on mountainsides by adding sediments collected from the lake centre, and designed nutrient levels and N/P ratio based on nutrient conditions of this lake^32^.

To integrate the three proximal controls to examine how DOM-microbe associations vary under global change, we developed a framework called Energy-Diversity-Trait integrative Analysis (EDTiA) (Fig. 1). EDTiA relies on the construction of bipartite networks ^30^ to quantify the specialization between organic molecules and microbial taxa. These networks are investigated using measures of entropy such as the *H*_2_’ index ^31^, which quantify resource-consumer relationships at an ecosystem-level. For example, elevated *H*_2_’ values convey that there is a high degree of specialization between DOM and microbes ^31^, where in the extreme example, one bacterial taxon consumes or produces a single DOM molecule. By contrast, lower *H*_2_’ reflects a more generalized bipartite network where different DOM molecules can be used by a large range of bacterial taxa. Furthermore, EDTiA allows for the integration of global change drivers to explore their relative importance of proximal controls on the specialization of DOM-microbe associations (Fig. 1).

We therefore used the EDTiA framework to test how associations between DOM and bacteria were jointly influenced by temperature and nutrient loading in a manipulative field experiment on subtropical and subarctic mountainsides in China and Norway ^32^. This macroecological approach involved creating microcosms with consistent initial DOM composition but different locally colonised microbial communities and newly produced endogenous DOM. Briefly, we selected five locations with different elevations on each mountainside that spanned a mean annual temperature gradient of 4.2-12.9°C in China and −2.9-0.7 °C in Norway. We established 300 sterile aquatic microcosms composed of natural lake sediments and artificial lake water, which included ten nutrient levels at each elevation. The sediments originated from Taihu Lake, a large eutrophic shallow lake in China, and were added to each microcosm after sterilisation to ensure identical initial DOM supply and composition. Microcosms were left in the field for one month allowing airborne bacteria to colonise, and sediment bacteria were examined using high-throughput sequencing of 16S rRNA genes ^32^. Additionally, we applied ultrahigh-resolution electrospray ionization Fourier transform ion cyclotron resonance mass spectrometry (FT-ICR MS) to examine sediment DOM features, such as chemodiversity and molecular traits.

Our study addresses three questions: (1) How do associations between chemodiversity and microbial biodiversity respond to temperature and nutrient enrichment? (2) How does the specialization of molecular-level associations between DOM and microbes vary along temperature and nutrient gradients? (3) How is the specialization interactively and independently influenced by temperature and nutrient enrichment via the three proximal controls? Results from our study will help advance biogeochemical modeling and improve predictions about carbon turnover along with feedbacks based on resource-based constraints on microbial diversity.

## Results and Discussion

### (1) DOM features and their microbial associations at a compositional-level

The diversity and molecular traits of DOM were strongly controlled by nutrient enrichment but less by temperature in both mountainsides (Figs. S2-4). Nutrient enrichment generally promoted molecular richness in both regions when all molecular components were considered (Figs. 2a, S5). Using piecewise regression ^33^ and gradient forest analysis ^34^, we identified breakpoints in molecular composition that mostly occurred between 1.80 and 4.05 mg N L^-1^ along the nutrient gradient for all molecules at each elevation (Figs. 2a, S6). The effects of nutrient enrichment on molecular traits, however, varied between the two ecoregions (Figs. 2a, S5, S7). For instance, the weighted mean of the H/C ratio in each microcosm decreased with nutrient addition to < 1.5, especially at high elevations in China, indicating less bioavailable DOM (Figs. 2a, S5c). The ratio remained consistently higher (≥ 1.5) across all nutrient levels in Norway (Figs. 2a, S5c). Given the identical DOM composition in our study initially, this finding suggests that the contrasting responses reflect differences in the temperature-sensitivity of decomposition and/or nutrient-limited production of DOM by colonising microbes. This inability to resolve the mechanisms underlying these patterns highlights the need for a more mechanistic approach offered by the EDTiA framework.

**Figure 2.**
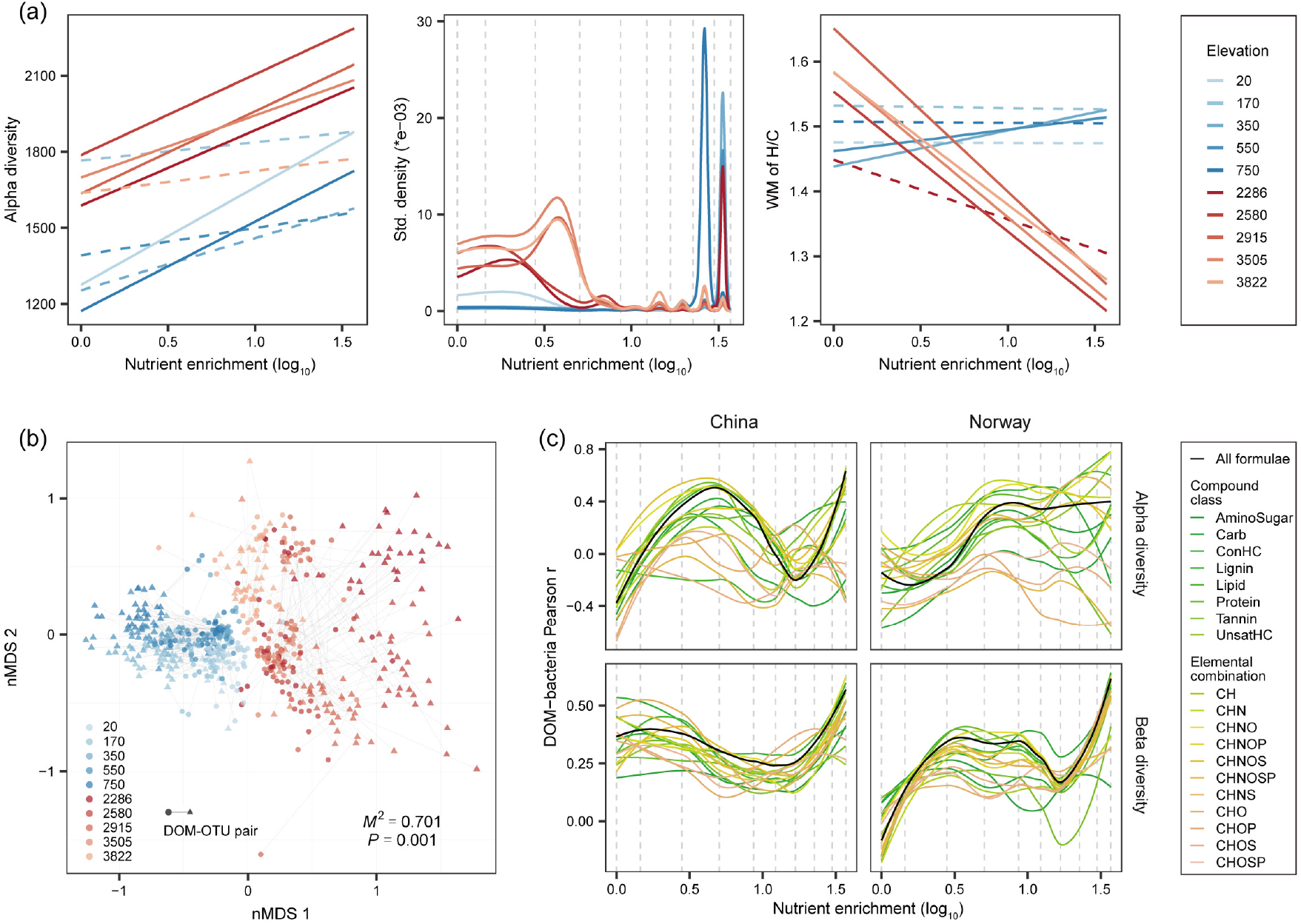
DOM features and their microbial associations at a compositional level. (a) The effects of nutrient enrichment on DOM alpha diversity (richness), composition and molecular traits (e.g., H/C ratio) for all formulae across different elevations in China (red lines) and Norway (blue lines). Molecular richness and weighted mean (WM) of H/C ratio were plotted against the nutrient gradient of nitrate, and their relationships are indicated by solid (*P* ≤ 0.05) or dotted (*P* > 0.05) lines estimated using linear models. For better visualization, we did not include the data points in Fig. 2a but showed detailed scatter plots in Fig. S5. To visualise the compositional turnover of DOM, we plotted the standardised density of splits showing where important changes in the abundance of multiple molecules occurred along the nutrient gradient. The standardised density of splits was determined by gradient forest analysis ^34^. (b) The congruence between DOM and bacterial compositions across different elevations in China and Norway was examined using Procrustes analysis ^35, 36^. Each line with circle and triangle ends connects to a single community of DOM and bacteria, respectively, and is colored by elevation in either China (red) or Norway (blue). The fit of overall Procrustes transformation is reported as the *M*^2^ value. (c) The effects of nutrient enrichment on DOM-microbe associations. The associations were quantified by the Pearson correlation coefficient r between alpha diversity of DOM and bacteria (upper panel), and by the Mantel r between the beta diversity of DOM and bacteria (lower panel). We then visualised these associations with loess regression models along the nutrient gradient. The colours of the lines indicate the DOM composition for all formulae and categories of compound classes or elemental combinations.

DOM composition was strongly associated with bacteria in both regions, and was mediated by temperature and nutrient enrichment. For instance, although environment (temperature and nutrients) and energy supply had dominant effects on DOM composition, their shared effects with biodiversity (1.0 to 26.7% of explained variation) indicated that these variables also indirectly influenced the associations between DOM and bacteria (Figs. S8-10). These DOM-microbe associations were also supported by a Procrustes analysis ^35, 36^, which revealed that more similar mixtures of molecules were related to more similar bacterial communities (*M*^2^ = 0.701, *P* ≤ 0.001; Fig. 2b), and their associations varied with temperature (that is, elevation) and nutrient enrichment. For example, compositional differences, indicated by the residuals of Procrustes analysis, significantly (*P* ≤ 0.05) decreased for all compound classes or elemental combinations at colder temperatures in China (Fig. S11). In Norway, the differences were always lower, on average, and did not vary with temperature (Fig. S11). Nutrient enrichment similarly influenced the correlations between DOM molecular and bacterial compositions estimated from both alpha and beta diversity (Fig. 2c), and these correlations also varied with nutrients for individual compound classes or elemental combinations (Figs. S12, S13). Interestingly, the coordinated compositional changes in DOM and bacteria, measured by the correlation between beta diversities, increased more strongly with nutrient enrichment in Norway than in China, especially at low nutrient levels beneath 1.80 mg N L^-1^ (Figs. 2c, S13).

### (2) Networks between DOM and bacteria at a molecular-level

To quantify the associations between DOM and bacteria further at a molecular-level, we first correlated the relative abundance of DOM molecules and bacterial taxa. According to resource-consumer relationships, negative associations likely indicate the degradation of larger molecules into smaller structures, while positive associations may relate to the production of new molecules via degradation or biosynthetic processes. We found that the distribution of negative and positive correlations between DOM molecules and bacterial operational taxonomic units (OTUs) depended strongly on molecular traits. For example, more labile molecules, such as those with H/C ≥ 1.5, were more likely to show negative Spearman’s correlation coefficients *ρ* with individual OTUs (*P* ≤ 0.05), whereas more recalcitrant molecules (H/C < 1.5) generally showed more positive correlations (*P* ≤ 0.05), especially in Norway (Fig. S14). These findings were even more clearly supported by the differences between the mean of the positive and negative *ρ* values for each molecule (Figs. 3a, S15). Correlations with individual OTUs were predominantly negative for molecules within a H/C of 1.5-2.0 and O/C of 0.4-1.0, suggesting they were the outcome of degradation processes, while *ρ* differences peaked with mainly positive values at a H/C of 1.0-1.5 and O/C of 0-0.5 indicative of *in situ* production (Fig. 3a).

**Figure 3.**
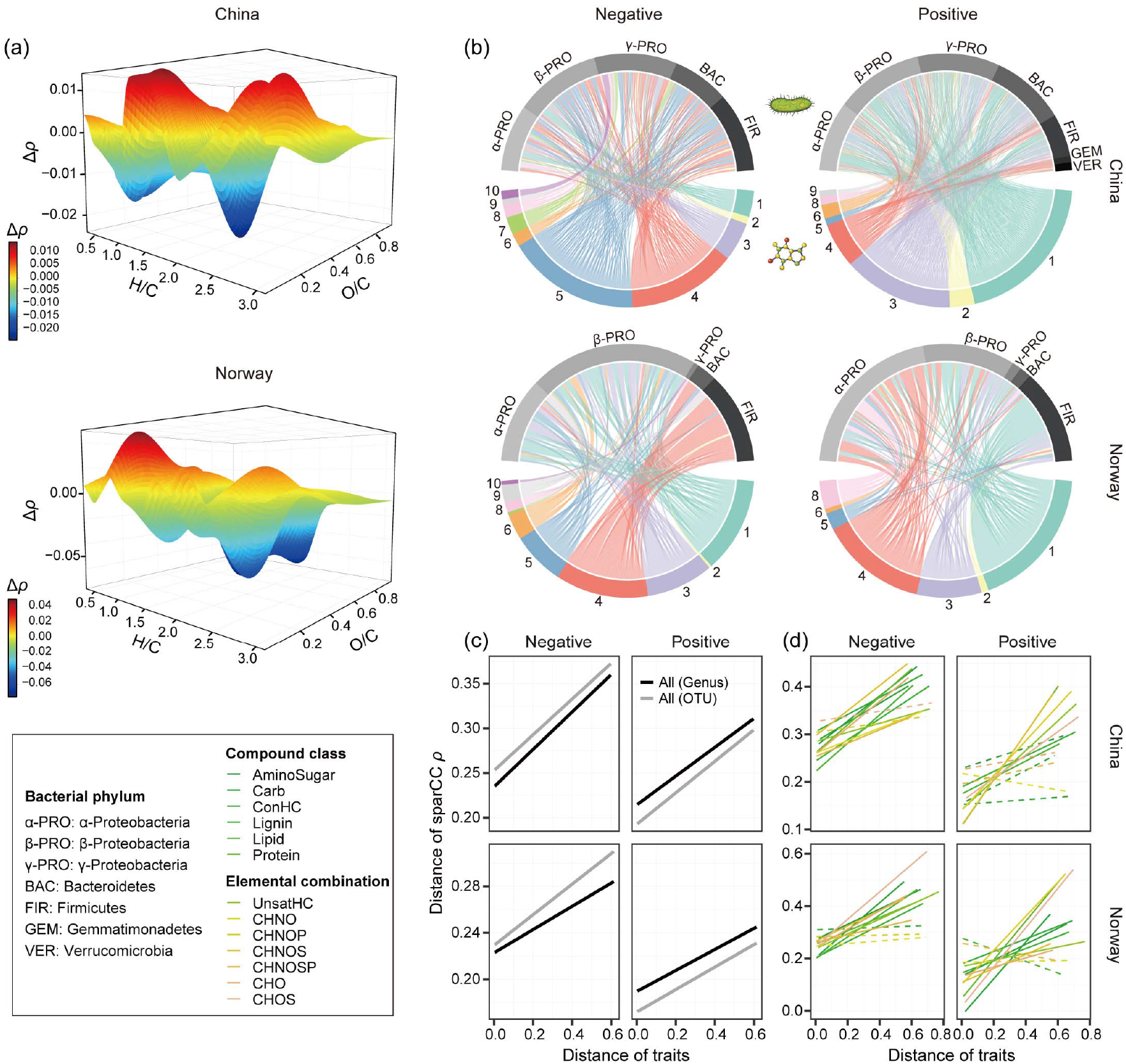
Networks between DOM and bacteria. (a) Strength of the correlations between DOM molecules and bacterial OTUs in China (upper panel) and Norway (lower panel). For each molecule, we subtracted the mean absolute Spearman’s rank correlation coefficient *ρ* of all the negative correlations with individual bacterial OTUs from the mean of the positive correlations to derive Δ*ρ*. Δ*ρ* was further visualised against the molecular traits H/C and O/C. (b) The negative and positive bipartite networks between DOM molecules and bacterial genera in China or Norway estimated using SparCC (Sparse Correlations for Compositional data) ^37^. Upper nodes represent bacterial genera coloured by their phylum, while lower nodes represent DOM molecules coloured by the ten clusters obtained with hierarchical cluster analysis based on 16 molecular traits described in Fig. S17 and Table S1. A line connecting two nodes indicates an interaction between a DOM molecule and bacterial genus. (c-d) We examined the relationships between molecular traits and the negative (left panel) or positive (right panel) DOM-microbe bipartite networks in China (upper panel) or Norway (lower panel). For all pairs of DOM molecules, we separately calculated pairwise Gower distances between the molecular traits and their SparCC *ρ* values with bacterial OTUs. Statistical significance between distance matrices was determined with a Mantel test with 999 permutations and indicated by solid (*P* ≤ 0.05) or dotted (*P* > 0.05) lines. We considered all formulae (c) and also subsets of formulae within the category of compound classes or elemental combinations (d). For all formulae (c), we calculated SparCC correlation coefficients based on both bacterial OTUs (grey lines) and genera (black lines).

Subsequently, we quantified DOM-microbe associations along temperature and nutrient gradients using the EDTiA framework. We built bipartite networks of negative and positive interactions between DOM and bacteria at the genus level using Sparse Correlations for Compositional data (SparCC) ^37^. SparCC relies on algorithms for sparse neighborhood and inverse covariance selection, and can infer correlations with a high degree of accuracy under these conditions ^37^. In total, there were 6,916 and 8,409 interactions for negative and positive networks (|SparCC *ρ*| ≥ 0.3), respectively, in China, and 1,313 and 2,888 negative and positive interactions, respectively, in Norway (Fig. 3b). The weighted mean of the percentage of SparCC *ρ* values that were strongly negative (*P* ≤ 0.05) increased towards high nutrient levels, with the reverse pattern for positive networks, almost exclusively in China (Fig. S16). Such patterns were consistent with the weighted mean SparCC *ρ* in China (Fig. S16).

The negative and positive interaction networks strongly depended on molecular traits, which was further supported by three observations. First, negative and positive networks were associated with different molecule groups categorised by hierarchical cluster analysis based on the 16 molecular traits (Figs. 3b, S17). In China, negative interactions were dominant between molecule clusters 4 and 5, which were largely comprised of recalcitrant molecules with a H/C of < 1.5 (Fig. S17), and bacteria in the phyla Proteobacteria, Bacteroidetes, or Firmicutes (Fig. 3b). The positive interactions were mostly linked to clusters 1 and 3 (Fig. 3b), which mostly represented labile molecules with a H/C of ≥ 1.5 (Fig. S17). In Norway, molecule cluster 4 was mainly negatively linked to Firmicutes and positively linked to α- and β-Proteobacteria (Fig. 3b). Second, molecules generally covaried more similarly with microbes as they shared more similar traits. For example, we detected statistically significant correlations between the pairwise Gower distances ^38^ of the traits and SparCC *ρ* values of DOM molecules in each region (Mantel test, *P* ≤ 0.001; Fig. 3c). Third, molecular traits were more strongly correlated with SparCC *ρ* in the negative than positive interaction networks for all molecules (Fig. 3c), which was also true for most of the networks when considering compound classes or elemental combinations (Fig. 3d). These correlations, consistent at the OTU level (Fig. 3c), indicate that molecular traits may be better at predicting the decomposition than production of DOM.

Finally, we calculated the degree of specialization between DOM and bacteria in the entire negative and positive interaction networks using the *H*_2_’ index ^31^. We also calculated specialization *d*’ indices for individual DOM molecules and bacterial genera ^31^. Elevated *H*_2_’ or *d*’ values indicate a high degree of specialization, while lower values suggest increased generalization. We found that networks that were more specialized in the negative associations between DOM and bacteria (i.e., higher *H*_2_’ values) corresponded with more specialized communities of DOM molecules (i.e., higher weighted mean *d*’; Figs. S18, S19) ^31^. For positive networks, *H*_2_’ values mirrored those of *d*’ for both DOM and bacteria (Figs. S18, S19). These results suggest that in addition to the specialization perspective of bacteria or DOM, *H*_2_’ can summarise resource-consumer relationships at an ecosystem-level. In both regions, *H*_2_’ was higher, on average, in negative than positive interaction networks (*t*-test, t = 2.11, *P* = 0.04 in China and t = 23.57, *P* ≤ 0.001 in Norway; Figs. 4a, S20), indicating a higher degree of specialization in the decomposition than production processes of microbes. Copiotrophs may have a high substrate specificity for labile resources as compared with oligotrophs ^39^, which have multiple metabolic pathways for resource acquisition of complex organic matter and hence lower specialization ^40^. The mean specialization *H*_2_’ of negative (*t*-test, t = −10.19, *P* ≤ 0.001) and positive (*t*-test, t = −6.56, *P* ≤ 0.001) networks were also significantly higher in Norway than in China (Fig. 4a), suggesting more specialized decomposition (i.e., negative networks) and thus potentially more degradable DOM in subarctic regions.

**Figure 4.**
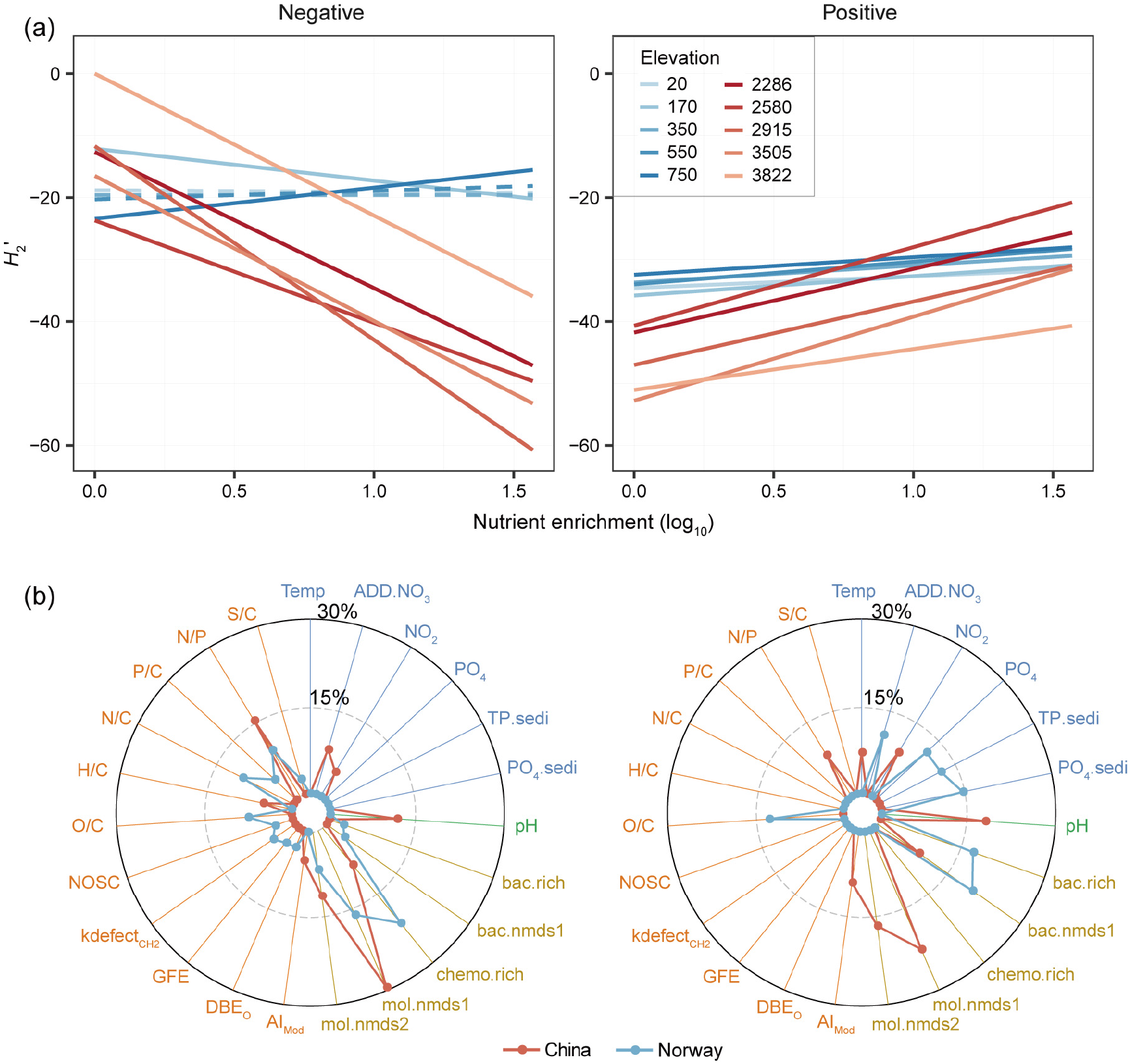
Relative importance of diversity and molecular traits in explaining specialization of DOM-microbe networks. (a) We plotted specialization *H*_2_’ against nutrient enrichment for negative (left panel) and positive (right panel) bipartite networks for each elevation in China (red lines) and Norway (blue lines). Statistical significance of linear model fits was indicated by solid (*P* ≤ 0.05) or dotted (*P* > 0.05) lines. For better visualization, we omitted the data points but these are shown in Fig. S20. (b) We examined the relative importance of all explanatory variables on the *H*_2_’ of negative (left panel) and positive (right panel) bipartite networks in China (red lines) and Norway (blue lines) using random forest. The relative contribution (%) of each variable towards *H*_2_’ is shown in radar plots. The explanatory variables were grouped by environment, energy, diversity and traits with consistent colors of ovals or rectangles as in Fig. S1. Abbreviations of explanatory variables are detailed in Table S1.

Nutrient enrichment showed divergent effects on the *H*_2_’ of negative or positive interaction networks between the two study regions. Specifically, nutrient enrichment substantially decreased the *H*_2_’ of negative networks for all molecules in China (Fig. 4a), which was particularly true when considering only recalcitrant components, such as lignin and CHNO (Fig. S21). Compared to Norway, nutrient enrichment increased the *H*_2_’ of positive interactions relatively more at lower elevations in China (Fig. 4a). Nutrient enrichment at the warmer temperatures in the subtropical region could thus contribute to the greater recalcitrance of DOM by reducing the specialization of decomposition (i.e., negative networks) and resulting in more specialized production of molecules (i.e., positive networks).

### (3) Drivers of DOM-microbe associations

We explored the following distal and proximal controls on negative and positive DOM-microbe networks under the EDTiA framework (Fig. 1). The distal drivers were temperature and nutrient enrichment as proxies of climate change and human impacts, respectively. The three proximal drivers were energy supply, such as primary productivity and sediment total organic carbon, the diversity of bacteria and DOM, that is the richness and composition of bacteria and DOM, and the DOM molecular traits (Table S1). In addition to bacterial diversity and chemodiversity, molecular traits strongly correlated with *H*_2_’ and influenced it through hypothesised casual relationships in structural equation models (SEM) ^41^ (Fig. S1).

The importance of molecular traits was supported by Pearson correlations (Fig. S22), multiple regression models (Fig. S23) and random forest analyses ^42^ (Fig. 4b). For the negative networks, *H*_2_’ showed the highest Pearson correlation coefficient of *r* = 0.77 with molecular composition (*P* ≤ 0.001), followed by molecular richness (*r* = −0.76, *P* ≤ 0.001) and N/P ratio (*r* = 0.76, *P* ≤ 0.001, Fig. S22). In contrast, *H*_2_’ was less correlated with molecular traits for the positive networks (Fig. S22). Multiple regression models revealed that, for negative and positive networks in China, there were statistically significant (*P* ≤ 0.01) improvements in the explained variances of models by between 6.2% and 9.1% from including either diversity or molecular traits (Fig. S23). These improvements were larger for the negative interaction networks in Norway (Fig. S23). These effects of diversity and molecular traits were further supported by random forest analyses. Diversity and molecular traits improved the predictive power of models of *H*_2_’ by 7.9-26.1% and 2.1-14.8%, respectively, and again, most strongly for the negative interactions in both regions (Fig. S23). Furthermore, *H*_2_’ was mainly affected by chemodiversity, such as molecular richness or DOM composition, followed by molecular traits, such as N/P or N/C ratios, in the negative networks, whereas chemodiversity, biodiversity, environmental variables and energy supply were all similarly important in the positive networks (Fig. 4b).

We also used SEM to test the hypothesised effects of two global change drivers, climate change and human impacts, on the specialization of DOM-bacteria networks. We compared these effects to other drivers like contemporary nutrients, energy supply, biodiversity, chemodiversity and molecular traits (Fig. S1). The SEM results strongly indicated that there were different constraints on DOM-microbe specialization between negative and positive interaction networks. For the negative networks, both global change drivers strongly influenced *H*_2_’ through indirect effects on energy supply and molecular traits, especially in China (Figs. 5, S24). In contrast to Norway, both climate change and human impacts had larger total mean effects of −0.23 and −0.49, respectively, on the *H*_2_’ of negative networks in China (Fig. 5a). However, molecular traits had the dominant direct effects on *H*_2_’ in both China and Norway, with similar mean standardised effect size of 0.57 (*P* ≤ 0.001; Figs. 5b, S24). For the positive networks, there were large total mean effects of climate change (0.51 and −0.40 for China and Norway, respectively) and human impacts (0.44 and 0.62, respectively), both of which indirectly influenced *H*_2_’ similarly through biodiversity, chemodiversity and molecular traits (Figs. 5, S24).

**Figure 5.**
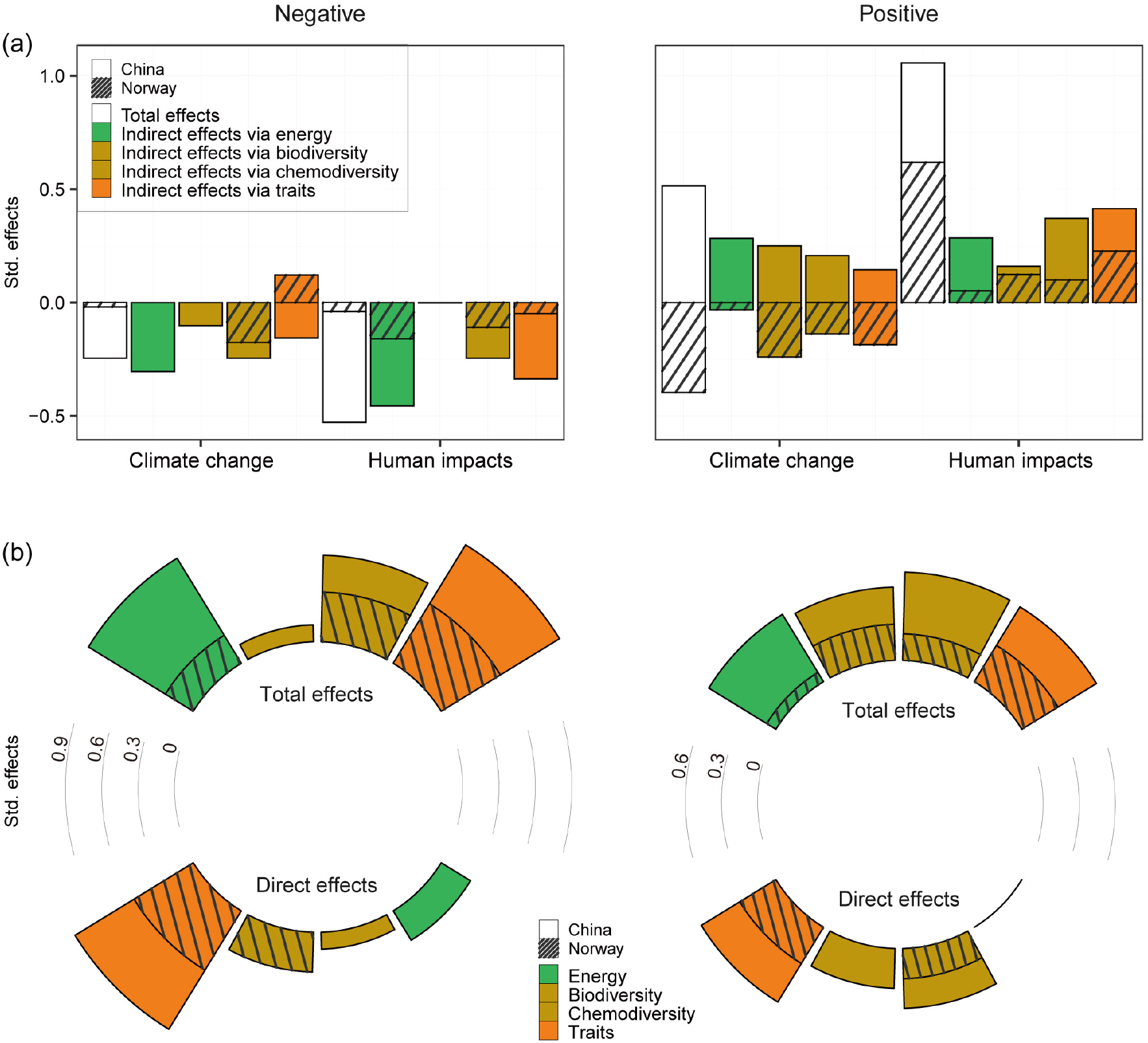
Structural equation models ^41^ to explain specialization of DOM-microbe networks. Stacked bar plots show the standardised effects (Std. effects) of predictor variables on the *H*_2_’ of negative (left panel) and positive (right panel) bipartite networks in China or Norway estimated from the best supported models. We considered (a) the total and indirect effects of global change and human impacts via proximal variables and (b) the total and direct effects of proximal variables. Proximal variables were energy supply, biodiversity, chemodiversity and molecular traits, and are described in detail in Table S2. Details of the full structural equation models are shown in Fig. S24.

### (4) Implications

The factors that control microbial processing of DOM composition, and consequently its degradation, remain challenging to discern ^43^, yet are critical for predicting carbon cycling under global change scenarios. We found that associations between DOM and microbial decomposers depended on universal drivers of ecosystem functioning, such as energy supply ^21, 22^, both DOM and microbial compositions ^8, 15^, and molecular traits ^7, 28^. The EDTiA framework we developed provides a unified approach to identify when each of these different proximal drivers is more important, and to separate contrasting biological processes associated with DOM degradation and production that may have obscured previous analyses of bulk DOM pools. In addition to energy supply and the diversity of DOM and bacteria, we found that molecular traits generally helped shape DOM-microbe networks across contrasting climatic zones, especially the negative networks indicative of degradation processes. Although molecular traits are well known to be associated with DOM persistence or vulnerability to degradation ^7, 44^, their influence on the underlying biological mechanisms has remained poorly understood. Our results advance this work by demonstrating when the specialization of DOM-microbe associations changes with molecular traits, and by providing predictions of how specialization might vary under global change.

We found that temperature and eutrophication can change DOM-microbe associations by shifting the three proximal drivers, namely energy, diversity, and traits. For positive bipartite networks, nutrient enrichment generally increased the specialization of DOM-microbe associations, and more so than temperature, by changing biodiversity, chemodiversity, and molecular traits. Positive interactions related to the production of new molecules depend on the specific interacting partners, which is partly supported by the positive relationships between *H*_2_’ and *d*’ (Fig. S19). By contrast, both temperature and nutrient enrichment reduced specialization in the negative networks, primarily via changing molecular traits and energy supply. Decomposition processes associated with negative networks may depend more on whether molecules contain structures that resist degradation ^7^, especially in the absence of temperature limitation ^45^. At higher temperatures, such as in subtropical China, energy to degrade these molecules may become more limiting ^45^. We also found that the importance of these distal drivers of climate change and human impacts varied between biomes. For instance, both elevated temperature and eutrophication reduced the specialization of negative DOM-microbe networks indicative of decomposition processes in subtropical China, but these two drivers were less important in subarctic Norway. As their indirect effects via microbial composition varied between biomes, these responses may reflect differences in the biological traits of communities. Future studies with metagenomics could offer a powerful complement to test how microbial traits vary with DOM traits.

As inland water worldwide continues to undergo changes in climate ^46^ and trophic state ^47^, our approaches could be applied to predict changes in how microbes degrade and produce DOM. For instance, since hyper-eutrophication occurred in Taihu Lake in May 2007, total nitrogen has been reduced by a mean (± SD) of 1.24 (± 1.41) mg L^-1^ with strong lake management (Figs. S25, S26). Based on the estimated direct and indirect effects of distal controls in the SEM fitted to the Chinese data (Fig. S1), this oligotrophication, combined with a mean decrease in water temperature of 0.20 (± 0.87) °C between 2007 and 2018, was predicted to change the specialization of DOM-microbe associations. Specifically, *H*_2_’ changed by +0.65 (± 0.58) and −0.65 (± 0.46) for negative and positive networks, respectively, over this period (Fig. 6a). The greatest changes happened in the most eutrophic part of the lake, including the northwestern lakeshore and the northern Zhushan and Meiliang Bays (Figs. 6b, S27). Although our predictions ignored detailed spatiotemporal environmental variations as used to parameterize the SEM models, they do illustrate the potential to upscale our predictions in real-world settings. This understanding, facilitated through our EDTiA framework, could provide the first steps for improving Earth system models of global biogeochemical cycles ^48^. More generally, our work shows how the molecular traits of DOM will control the responses of DOM-microbe networks and their associated biogeochemical cycles in a changing world.

**Figure 6.**
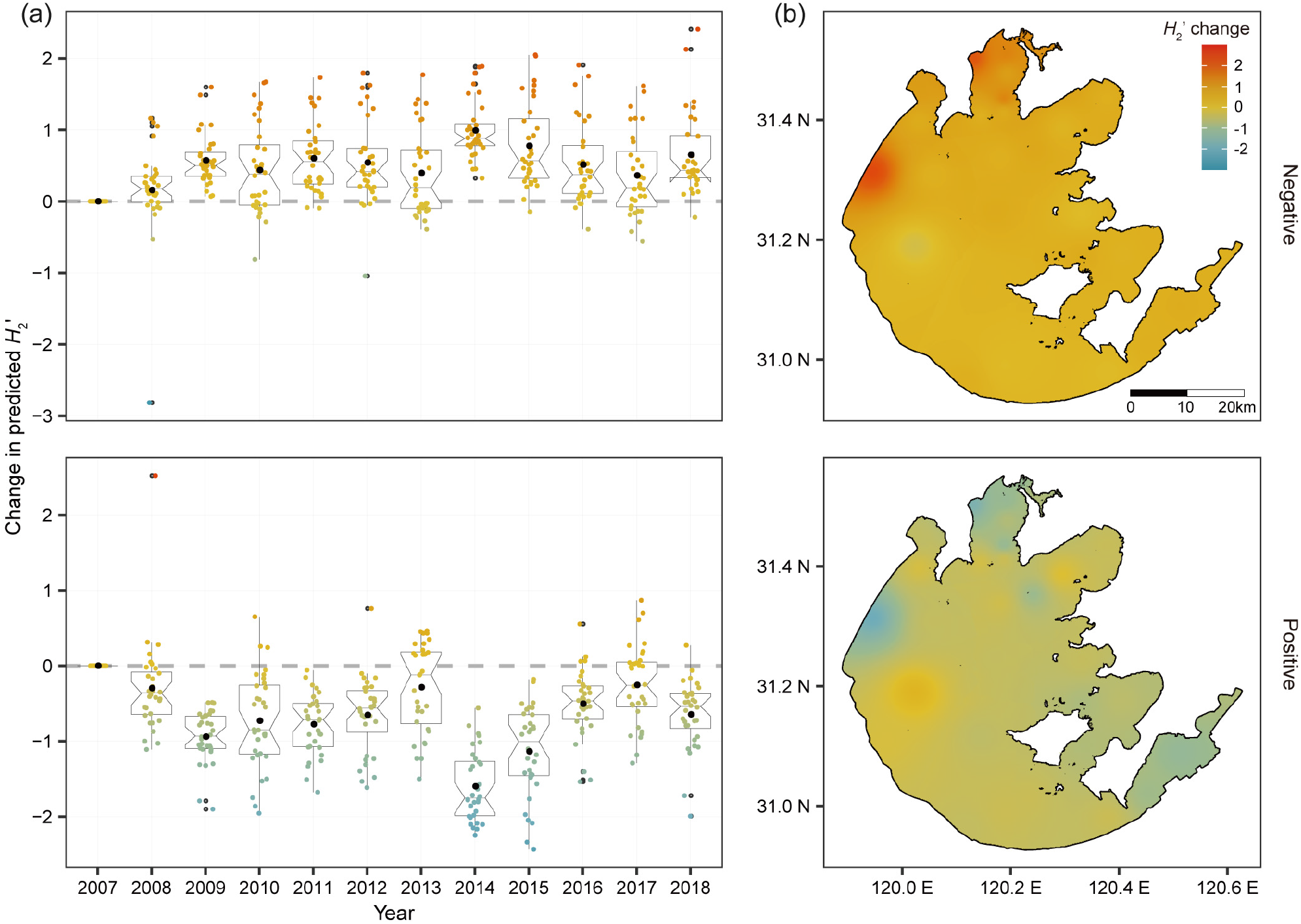
Decadal change in predicted specialization of DOM-microbe networks in Taihu Lake. (a) Changes in *H*_2_’ of negative (upper panel) and positive (lower panel) bipartite networks from 2007 to 2018. (b) The spatial distribution of changes in *H*_2_’ of negative (upper panel) and positive (lower panel) networks in 2018 across the Taihu Lake. Estimated changes in *H*_2_’ were calculated for the 32 sites across the whole of Taihu Lake (Fig. S27a) by comparing with the baseline of 2007, and represent the combined effects of climate change and eutrophication. The colored dots in (a) indicate *H*_2_’ changes for individual sites which are consistent with the figure legend of (b), and black dots are the mean values for each year. The box in (a) represents the interquartile (50% of data), the horizontal line in the box represents the median, the “notch” represents the 95% confidence interval of the median and the “whiskers” represent the maximum and minimum values.

## Acknowledgements

We appreciate C.Y. Zhang, L.Z. Dai and F.Y. Pan for field sampling and lab analyses, and T. Dittmar and C.L. Zhang for valuable suggestions. We thank Taihu Laboratory for Lake Ecosystem Research for providing long-term data of Taihu Lake. This study was supported by National Natural Science Foundation of China (91851117), CAS Key Research Program of Frontier Sciences (QYZDB-SSW-DQC043), and National Natural Science Foundation of China (42077052, 41871048). KJ was supported by Korea Basic Science Institute grants (C140222, C140444). AJT was supported by H2020 ERC Grant (sEEIngDOM 804673). JTL was supported by National Science Foundation (DEB-1442246, 1934554), US Army Research Office Grant (W911NF-14-1-0411), and the National Aeronautics and Space Administration (80NSSC20K0618).

## Conflict of Interest

The authors declare no conflict of interest.

## Author contributions

JW conceived the idea. JW carried out the field trips and provided the physiochemical and biological data. JL, MC and KJ analysed the DOM. AH and JW performed the statistical analyses. AH wrote the first draft of the manuscript. AH and JW finished the manuscript with the comments from AJT, JTL, JS, KJ, YL and XL. All authors contributed to the intellectual development of this study.

## Data availability

Bacterial sequences and environmental data are available in Wang *et al* (2016) ^32^. Other data are available from the corresponding author upon reasonable request.

## Material and methods

### Experimental design

The comparative field microcosm experiments were conducted on Laojun Mountain in China (26.6959 N; 99.7759 E) in September-October 2013, and on Balggesvarri Mountain in Norway (69.3809 N; 20.3483 E) in July 2013, designed to be broadly representative of subtropical and subarctic climatic zones, respectively, as first reported in Wang *et al*. (2016) ^32^. The annual temperatures ranged from 4.2-12.9 °C in China and −2.9-0.7 °C in Norway. The experiments were characterised by an aquatic ecosystem with consistent initial DOM composition but different locally colonised microbial communities and newly produced endogenous DOM. While allowing us to minimise the complexity of natural ecosystems, the experiment provided a means for investigating DOM-microbe associations at large spatial scales by controlling the initial DOM supply. Briefly, we selected locations with five different elevations on each mountainside. The elevations were 3,822, 3,505, 2,915, 2,580 and 2,286 m a.s.l. on Laojun Mountain in China, and 750, 550, 350, 170 and 20 m a.s.l. on Balggesvarri Mountain in Norway. At each elevation, we established 30 aquatic microcosms (1.5 L bottle) composed of 15 g of sterilised lake sediment and 1.2 L of sterilised artificial lake water, which included one of ten nutrient levels of 0, 0.45, 1.80, 4.05, 7.65, 11.25, 15.75, 21.60, 28.80 and 36.00 mg N L^-1^ of KNO_3_. Each nutrient level was replicated three times. The lake sediments were obtained from the centre of Taihu Lake, China, and were aseptically canned per bottle after autoclaving as previously described in Wang *et al*. (2016) ^32^. To compensate for nitrate additions shifting stoichiometric ratios, KH_2_PO_4_ was added to bottles so that the N/P ratio of the initial overlying water was 14.93, which was similar to the annual average ratio in Taihu Lake during 2007 (14.49). Nutrient levels for the experiments were selected based on conditions of the eutrophic Taihu Lake, and the highest nitrate concentration was based on the maximum total nitrogen in 2007 (20.79 mg L^-1^; Fig. S27). We chose the nutrient level of this year because a massive cyanobacteria bloom in Taihu Lake happened in May 2007 and initiated an odorous drinking water crisis in the nearby city of Wuxi. The microcosms were left in the field for one month allowing airborne bacteria to freely colonise the sediments and water, and the sediment bacteria were examined using high-throughput sequencing of 16S rRNA genes. The sequences were processed in QIIME (v1.9) ^49^ and OTUs were defined at 97% sequence similarity. The bacterial sequences were rarefied to 20,000 per sample. Further details on field experiments, sample collection, physicochemical and bacterial community analyses are available in Wang *et al*. (2016).

### ESI FT-ICR MS analysis of DOM samples

Highly accurate mass measurements of DOM within the sediment samples were conducted using a 15 Tesla solariX XR system, a ultrahigh-resolution Fourier transform ion cyclotron resonance mass spectrometer (FT-ICR MS, Bruker Daltonics, Billerica, MA) coupled with an electrospray ionization (ESI) interface, as demonstrated previously^50^ with some modifications. DOM was solid-phase extracted (SPE) with Agilent VacElut resins before FT-ICR MS measurement ^51^ with minor modifications. Briefly, an aliquot of 0.7 g freeze-dried sediment was sonicated with 30 ml ultrapure water for 2 h, and centrifuged at 5,000 g for 20 min. The extracted water was filtered through the 0.45 µm Millipore filter and further acidified to pH 2 using 1 M HCl. Cartridges were drained, rinsed with ultrapure water and methanol (ULC-MS grade), and conditioned with pH 2 ultrapure water. Calculated volumes of extracts were slowly passed through cartridges based on DOC concentration. Cartridges were rinsed with pH 2 ultrapure water and dried with N_2_ gas. Samples were finally eluted with methanol into precombusted amber glass vials, dried with N_2_ gas and stored at −20 °C until DOM analysis. The extracts were continuously injected into the standard ESI source with a flow rate of 2 μl min^-1^ and an ESI capillary voltage of 3.5 kV in negative ion mode. One hundred single scans with a transient size of 4 mega words, an ion accumulation time of 0.3 s, and within the mass range of m/z 150-1200, were co-added to a spectrum with absorption mode for phase correction, thereby resulting in a resolving power of 750,000 (FWHM at m/z 400). All FT-ICR mass spectra were internally calibrated using organic matter homologous series separated by 14 Da (-CH_2_ groups). The mass measurement accuracy was typically within 1 ppm for singly charged ions across a broad m/z range (150-1,200 m/z).

Data Analysis software (BrukerDaltonik version 4.2) was used to convert raw spectra to a list of m/z values using FT-MS peak picker with a signal-to-noise ratio (S/N) threshold set to 7 and absolute intensity threshold to the default value of 100. Putative chemical formulae were assigned using the software Formularity ^52^ following the Compound Identification Algorithm ^53^. In total, 19,538 molecular formulas were putatively assigned for all samples (n = 300) based on the following criteria: S/N > 7, and mass measurement error < 1 ppm, considering the presences of C, H, O, N, S and P and excluding other elements or an isotopic signature. All formula assignments were further screened to meet the criteria as follows ^54^: (1) formulae containing an odd number of nitrogen atoms had an even nominal m/z and those containing an even number of nitrogen atoms had an odd nominal m/z; (2) the number of hydrogen atoms was at least 1/3 of carbon and could not exceed 2C+N+2; (3) the number of nitrogen or oxygen atoms could not exceed the number of carbon atoms; (4) the ratio of O/C was set to 0-1, H/C ≥ 0.3, N/C ≤ 1, double bond equivalents (DBE) ≥ 0.

The assigned molecules were categorised into eight compound classes or 12 elemental combinations. The compound classes based on van Krevelen diagrams ^55^ were lipids (O/C = 0-0.3, H/C = 1.5-2.0), proteins and amino sugars (O/C = 0.3-0.67, H/C = 1.5-2.2), carbohydrates (Carb; O/C = 0.67-1.2, H/C = 1.5-2), unsaturated hydrocarbons (UnsatHC; O/C = 0-0.1, H/C = 0.7-1.5), lignin (O/C = 0.1-0.67, H/C = 0.7-1.5), tannin (O/C = 0.67-1.2, H/C = 0.5-1.5), and condensed aromatics (ConHC; O/C = 0-0.67, H/C = 0.2-0.7). The elemental combinations were CH, CHN, CHNO, CHNOP, CHNOS, CHNOSP, CHNS, CHO, CHOP, CHOS, CHOSP and CHS.

### Estimating DOM features

We considered DOM features from three aspects: alpha diversity, beta diversity and molecular traits. These features were considered for all molecules (19,538 different formulae), but also for subsets of molecules within each category of compound classes or elemental combinations. The dataset based on all molecular formulae was named “All molecules”, while the datasets of subsets of formulae were named by “category name + compounds”. The relative abundance of molecules was calculated by normalizing signal intensities of assigned peaks to the sum of all intensities within each sample. We considered two additional aspects of chemodiversity: chemical alpha diversity and chemical beta diversity. Chemical alpha diversity was calculated using a richness index that counts the total number of peaks in each sample. Chemical beta diversity was calculated with the Bray-Curtis dissimilarity metric, and further represented by the first two axes of a non-metric multidimensional scaling (NMDS) ordination of this dissimilarity. We also considered overall molecular composition, which was visualised across the elevations and nutrient enrichment treatments with detrended correspondence analysis (DCA) ^56^. The analyses of chemical diversity were performed using the R package vegan V2.4.6 ^57^.

We also calculated 16 molecular traits that could affect microbial associations and were related to molecular weight, stoichiometry, chemical structure, and oxidation state (Table S1). These traits were mass, the number of carbon (C) atoms, the modified aromaticity index (AI_Mod_) ^58^, DBE ^58^, DBE minus oxygen (DBE_O_) ^58^, DBE minus AI (DBE_AI_) ^58^, standard Gibb’s Free Energy of carbon oxidation (GFE) ^59^, Kendrick Defect (kdefect_CH2_) ^60^, nominal oxidation state of carbon (NOSC), O/C ratio, H/C ratio, N/C ratio, P/C ratio, S/C ratio, and carbon use efficiency (Y_met_) ^61^. All calculations were performed using the R package ftmsRanalysis V1.0.0 ^62^ and the scripts at https://github.com/danczakre/ICRTutorial. DBE represents the number of unsaturated bonds and rings in a molecule ^58^. Higher values of DBE, AI and NOSC all indicate a higher recalcitrance of DOM. A large Kendrick Defect can indicate a higher degree of oxidation. Lower values of Y_met_ indicate a higher thermodynamic efficiency of metabolic reactions involved in biomass production ^61^. Weighted means of formula-based molecular traits (for example the Mass_wm_ for Mass) were calculated as the sum of the product of the trait value for each individual molecule (Mass_i_) and relative intensity *I*_i_ divided by the sum of all intensities (Mass_wm_ = Σ(Mass_i_ × *I*_i_) / Σ(*I*_i_)) using the R package FD V1.0.12 ^63^. In addition, ten molecular sub-mixtures were grouped based on the 16 molecular traits by hierarchical cluster analysis using Ward’s minimum variance method with the R package stats V3.6.1.

### Estimating bacterial communities

The relative abundance of OTUs was calculated by the normalization of read counts of OTUs to the sum of all reads within each sample. Likewise, we considered two aspects of biodiversity: bacterial alpha diversity and beta diversity. Bacterial alpha diversity was calculated using species richness that counts the total number of OTUs in each sample. Bacterial beta diversity was calculated with the Bray-Curtis dissimilarity metric, and further represented by the first two axes of NMDS of this dissimilarity.

### Estimating associations between DOM and microbes

At the DOM composition level, we examined DOM-microbe associations from the following aspects: Pearson’s correlation between alpha diversity of DOM and bacteria, and a Mantel correlation between the beta diversity of DOM and bacteria. We also tested the congruence between DOM and bacterial composition using Procrustes analysis of NMDS coordinates estimated for each community across elevations and nutrient enrichment levels with the Bray-Curtis dissimilarity metric ^35, 36^. Procrustes analysis is a technique for comparing the relative positions of points in two multivariate datasets. It attempts to stretch and rotate the points in one matrix, such as points obtained from a NMDS, to be as close as possible to points in another matrix, thus preserving the relative distances between points within each matrix ^35, 36^. This procedure yields a measure of fit, *M*^2^, which is the sum of squared distances between corresponding data points after the transformation. Pointwise residuals indicate the difference between two different community ordinations for each sample. The statistical significance of the Procrustes analysis (i.e., *M*^2^) can then be assessed by randomly permutating the data 1,000 times ^64^. This analysis was performed using the R package vegan V2.4.6.

We further quantified DOM-microbe associations at a molecular level using two different co-occurrence analyses. First, Spearman’s rank correlation coefficient *ρ* was calculated between the relative abundance of each molecule m/z ion and bacterial OTU (or genus). For each molecule, we then calculated the Spearman *ρ* difference by subtracting the mean absolute *ρ* value of the negative correlations across all bacterial OTUs from the mean of the positive correlations. Larger positive and negative values indicate that molecules were more strongly positively and negatively correlated with bacterial communities, respectively. The relationships among the Spearman *ρ* difference, H/C and O/C were summarised using kriging interpolation with the R package automap V1.0.14 ^65^. Second, SparCC (Sparse Correlations for Compositional data) was applied to build DOM-microbe bipartite networks. SparCC is a correlation method that can infer the interrelationships between DOM and bacteria for compositional data with higher accuracy ^37^ than general correlation approaches, such as Spearman’s correlation, because it explicitly assumes that the underlying networks have many missing associations. We used bacterial genera rather than OTUs for bipartite network analysis because there were over 20,000 and 10,000 bacterial OTUs for Norway and China, respectively, and there are computational limits on handling such large bipartite networks for the analyses described in the next paragraph. However, using bacterial genera was reasonable as individual DOM-bacteria associations were similar for both bacterial OTUs and genera (R^2^ > 0.80, *P* ≤ 0.001; Fig. S14). Similar conclusions were also obtained with either OTUs or genera when relating the pairwise distances of molecular traits with SparCC *ρ* values among DOM molecules in Fig. 3c. To reduce type I errors in the correlation calculations created by low-occurrence genera or molecules, the majority rule was applied, retaining genera or molecules observed in more than half of the total samples (≥ 75 samples) in China or Norway. The filtered table, including 1,340 and 1,246 DOM molecules, and 75 and 49 bacterial genera in China and Norway, respectively, was then used for pairwise correlation calculation of DOM and bacteria using SparCC with default parameters ^37^.

Finally, bipartite network analysis at a molecule and network level was performed to quantify the specialization of DOM-microbe associations. The threshold correlation for inclusion in bipartite networks was |*ρ*| = 0.30 to exclude weak interactions and we retained the adjacent matrix with only the interactions between DOM and bacteria. We then constructed two types of networks based on negative and positive correlations (SparCC *ρ* ≤ −0.30 and *ρ* ≥ 0.30, respectively). The SparCC *ρ* values were multiplied by 10,000 and rounded to integers, and the absolute values were taken for negative networks to enable the calculations of specialization indices. A separate negative and positive network was obtained for each microcosm based on its species composition. For the network level analysis, we calculated *H*_2_’, a measure of specialization ^30^, for each network:

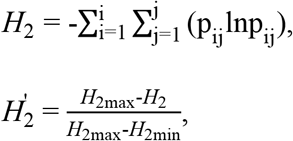

where p_ij_= a_ij_/m, represents the proportion of interactions in a i × j matrix. a_ij_ is number of interactions between DOM molecule i and bacterial genus j, which is also referred as “link weight”. m is total number of interactions between all DOM molecules and bacterial genera. *H*_2_’ is the standardised *H*_2_ against the minimum (*H*_2min_) and maximum (*H*_2max_) possible for the same distribution of interaction totals.

For the molecular level analysis, we calculated the specialization index Kullback-Leibler distance (*d*’) for DOM molecules 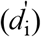 and bacterial genera 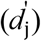, which describes the levels of “vulnerability” of DOM molecules and “generality” of bacterial genera, respectively:

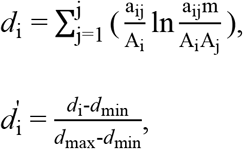

where 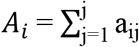 and 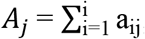, are the total number of interactions of DOM molecule i and bacterial genus j, respectively. *d*_i_’ is the standardised *d*_i_ against the minimum (*d*_min_) and maximum (*d*_max_) possible for the same distribution of interaction totals. The equations of 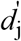 are analogous to 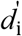, replacing j by i.

Both specialization indices consider interaction abundance and are standardised to account for heterogeneity in the interaction strength and species richness. Weighted means of *d*’ for DOM were calculated for each network as the sum of the product of *d*’ for each individual molecule i (*d*’_i_) and relative intensity *I*_i_ divided by the sum of all intensities *d*’ = Σ(*d*’_i_ × *I*_i_) / Σ(*I*_i_). Weighted means of *d*’ for bacteria were calculated as the sum of the *d*’ of each individual bacterial genus j (*d*’_j_) and relative abundance of bacterial genus *I*_j_ divided by the sum of all abundance. All calculations were performed using the R package FD V1.0.12. The observed *H*_2_’ and *d*’ values ranged from 0 (complete generalization) to 1 (complete specialization) ^31^ (Fig. S28). To directly compare the network indices across the elevations or nutrient enrichment levels, we used a null modelling approach. We standardised the three observed specialization indices (*S*_observed_; that is, *H*_2_’, *d*’ of DOM, and *d*’ of bacteria) by calculating their z-scores ^66^ using the equation 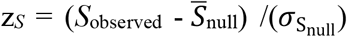 where 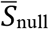 and 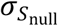 were, respectively, the mean and standard deviation of the null distribution of *S* (*S*_null_). One hundred randomised null networks were generated for each bipartite network to derive *S*_null_ using the *swap*.*web* algorithm, which keeps species richness and the number of interactions per species constant along with network connectance. The relationships among *H*_2_’, weighted means of *d*’ for DOM molecules and bacterial genera were compared using kriging interpolation with the R package automap V1.0.14. The obtained network was visualised using circlize V0.4.10 ^67^ and analysed using the R package bipartite V2.15 ^30^.

### Statistical analyses

We used the following explanatory variables related to distal and proximal controls on DOM-microbe associations. Distal environmental drivers included climate change (i.e., water temperature), human impacts (i.e., nutrient enrichment), and contemporary nutrients (i.e., sediment total nitrogen (TN), total phosphorus (TP), NO_x-_, NO_2-_, NH_4_^+^ and PO_4_^3-^, and water NO_3-_, NO2-, NH_4_^+^ and PO_4_^3-^). Proximal drivers included energy supply (i.e., sediment total organic carbon, dissolved organic carbon, water pH and sediment Chlorophyll *a* (Chl *a*)), biodiversity (i.e., the species richness and the first two axes of the NMDS of bacterial community composition), DOM chemodiversity (i.e., the species richness and the first two axes of the NMDS of molecular composition), and DOM molecular traits (i.e., mass, C, AIMod, DBE, DBEO, DBEAI, GFE, kdefectCH2, NOSC, O/C, H/C, N/C, P/C, S/C and Ymet). Detailed information about these explanatory variables is listed in Table S1. It should be noted that water pH could be considered to be relevant to primary productivity due to its strong positive correlation with sediment Chl *a*, but their relationships varied across elevations and nutrient levels ^32^. The response variables included DOM features (i.e., alpha diversity, beta diversity and molecular traits) and DOM-microbe network statistics (e.g., *H*2’), and were analysed for their patterns and underlying drivers along the two main environmental gradients: elevation and nutrient enrichment.

#### (1) Patterns of DOM features and DOM-microbe associations along the environmental gradients

For DOM features, the relationships between nutrient enrichment and DOM richness or molecular traits were visualised with linear models for all formulae and subsets of formulae within each category of compound classes or elemental combinations across different elevations. We further tested the breakpoints or abrupt changes in DOM composition (i.e., the first axis of DCA) along the gradient of nutrient enrichment using a piecewise linear regression with the R package segmented V1.3.0 ^33^. These breakpoint estimations were supported by gradient forest analysis ^34^, which was used to assess the DOM compositional changes and important breakpoints across multiple molecules along the gradient of nutrient enrichment. This analysis produces the standardised density of splits, that is the kernel density of splits divided by the observation density, which shows where important changes in the abundance of multiple molecules occur along the nutrient gradient and indicates the compositional rate of change. In addition, we estimated the standardised density of splits for subsets of molecules within each category of compound classes or elemental combinations across different elevations. This analysis was performed using the R packages gradientForest V0.1.17 ^34^ and extendedForest V1.6.1 ^68^.

For DOM-microbe associations, the relationships between nutrient enrichment and associations at both community and network levels were tested with linear models for all formulae and subsets of formulae within each category of compound classes or elemental combinations across different elevations.

#### (2) Drivers of DOM features and DOM-microbe associations

To evaluate the key drivers of DOM features and DOM-microbe associations, we used variation partitioning analysis (VPA) ^69^, multiple regression, random forest analysis^42^ and structural equation modelling (SEM) ^41^. In particular, the first analysis disentangled the important roles of microbes from other explanatory variables, while the latter three analyses tested the roles of molecular traits and diversity, and their interplay with environments and energy supply.

First, VPA was used to quantify the relative contributions of driver categories towards DOM features. We partitioned explanatory variables into the following driver categories: environments (that is, climate change, human impacts and contemporary nutrients), energy supply and biodiversity (Table S1). We selected explanatory variables for regression analyses by forward selection with Akaike information criterion (AIC) ^70^. We also quantified the relative contributions of driver categories for subsets of molecules within each category of compound classes or elemental combinations. VPA was performed with R package vegan V2.4.6 ^71^.

Second, stepwise multiple regression was performed to test the statistical significance and predictive power of the net effects of diversity (i.e., biodiversity and chemodiversity) or molecular traits on the bipartite network specialization *H*_2_’. The net effects of diversity or molecular traits were evaluated by the improvements in the explained variances relative to models without diversity and molecular traits (i.e., in models using only the variables associated with environments and energy supply). The analysis was conducted with forward selection of explanatory variables ^72^. We chose the final model that had the lowest AIC value ^73^. ANOVA was used to test the statistical significance of two models including or excluding diversity or molecular traits as predictors, and the increase in the model *R*^2^ was determined as the net effects of diversity or molecular traits on *H*_2_’.

Third, random forest analysis was conducted to identify the relative importance of environment variables, energy supply, bacterial diversity and DOM molecular drivers on specialization *H*_2_’. The importance of each predictor variable was determined by evaluating the decrease in prediction accuracy (that is, increase in the mean square error between observations and out-of-bag predictions) when the data for that predictor were randomly permuted. The accuracy importance measure was computed for each tree and averaged over the forest (2,000 trees). More details on this method were described in previous literature ^74^. In addition, random forest analysis was also used to test the net effects of diversity or molecular traits on *H*_2_’. This analysis was conducted using the R package randomForestSRC V2.8.0 ^75, 76^.

Finally, SEM was used to explore how specialization *H*_2_’ is interactively influenced by global changes (that is, temperature and nutrient enrichment), diversity and molecular traits. The approach begins by hypothesising the underlying structure of causal links as shown in Fig. S1. Then, the model is translated into regression equations, and these equations are evaluated against the data to test the hypothesised links. Through this process, SEM provides an understanding of direct and indirect links of climate change and human impacts on *H*_2_’. Before modelling, all variables in the SEMs were Z-score transformed to allow comparisons among multiple predictors and models. Similar to previous studies ^77^, we used composite variables to account for the collective effects of climate change, human impacts, contemporary nutrients, energy supply, biodiversity, chemodiversity and molecular traits, and the candidate observed indicators are given in Table S1. The indicators for each composite were selected based on the multiple regressions for *H*_2_’ (Table S2). Based on all the hypothesised links among composite variables (that is, full model; Fig. S1), we examined all alternative models using AIC and overall model fit statistics ^78^. We chose the final model to report as that with the lowest AIC value from models with a non-significant χ^2^ test (*P* > 0.05), which tests whether the model structure differs from the observed data, high comparative fit index (CFI > 0.95) and low standardised root mean squared residual (SRMR < 0.05) (Table S3). We implemented the SEMs using R package lavaan V.0.5.23 ^79^.

### Predictions of DOM-microbe associations in Taihu Lake

Using the parameter estimates obtained from SEM fitted to the bipartite networks in subtropical China, we estimated spatiotemporal variation of DOM-microbe associations in Taihu Lake based on the direct and indirect effects of climate change and eutrophication via the proximal drivers. We first formulated five linear equations to predict the values of contemporary nutrients (P_nut_), energy supply (P_energy_), biodiversity (P_biodiv_), chemodiversity (P_chemodiv_) and molecular traits (P_trait_) based on climate and eutrophication drivers:

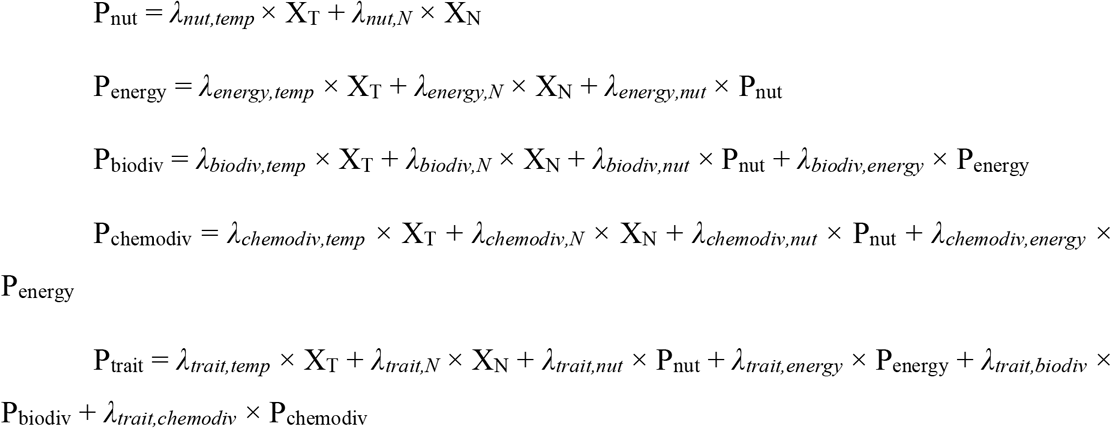

where X_T_ and X_N_ were water temperature and total nitrogen, respectively, for the 32 sites across the whole Taihu Lake (Fig. S27a). The abbreviations of path coefficients (*λ*) are detailed in Table S4.

Similarly, we calculated the specialization of DOM-microbe associations (Y_H2_) using a linear equation: Y_H2_ = *λ*_*H2,temp*_ × X_T_ + *λ*_*H2,N*_ × X_N_ + *λ*_*H2,nut*_ × P_nut_ + *λ*_*H2,energy*_ × P_energy_ + *λ*_*H2,biodiv*_ × P_biodiv_ + *λ*_*H2,chemodiv*_ × P_chemodiv_ + *λ*_*H2,trait*_ × P_trait_. We used the predicted values for contemporary nutrients, energy supply, biodiversity, chemodiversity and molecular traits in the overall prediction model to account for the indirect effects of water temperature and total nitrogen on specialization. The models were calculated with a yearly time step based on the annual means of water temperature and total nitrogen for each site during 2007-2018. The temporal changes in specialization were calculated using 2007 as a baseline to which all predictions were compared.

The above predictions aimed to apply our EDTiA framework to estimate changes in DOM-microbe associations under temperature change and eutrophication in Taihu Lake, and potential uncertainties in the estimated associations should however be noted as follows. First, local environmental variation (e.g., N/P ratio changes) and different microbial species pools between our field microcosms and natural lake sediments would likely influence the accuracy of predictions. Second, spatial and temporal heterogeneity of sediments would influence local environments and the composition of both DOM and microbes and thus the projection of estimates across Taihu Lake. Third, the transferability and extrapolation of SEM models to Taihu Lake would be one of the difficulties in prediction practices. We thus selected the SEM models in China rather than Norway for more similar climatic conditions to the target lake. The annual mean water temperatures in Taihu Lake were covered by the temperature variations across the elevations between 2,286 and 3,822 m a.s.l. in Laojun Mountain, and the annual mean total nitrogen fell into the gradient of nutrient concentrations between 0 and 36 mg N L^-1^. Finally, lake management such as mechanical removal of algae would affect energy supply and consequently prediction accuracy.

## Notes

### Competing Interest Statement

The authors have declared no competing interest.

