## Supplemental materials for "Quantifying microbial associations of dissolved organic matter under global change"

**The file includes:**

Supplementary Tables (S1–S4)

Supplementary Figures (S1–S28)

### Supplementary Tables

**Table S1**. Variables used to explain DOM features or the specialization of DOM-microbe associations. These variables were considered based on the hypothetical casual relationships in Fig. S1: climate change, human impacts and contemporary nutrient variables as distal drivers, and energy supply, biodiversity, chemodiversity and DOM traits as proximal drivers. NMDS: non-metric multidimensional scaling.

| Drivers | Group | Subgroup | Variable | Description |
| --- | --- | --- | --- | --- |
| Distal drivers | Environment | Climate change | Temp | Water temperature[#] |
| | | Human impacts | ADD.$NO_3$ | Nutrient enrichment |
| | | Contemporary nutrient | TN.sedi | Sediment total nitrogen (TN) |
| | | Contemporary nutrient | TP.sedi | Sediment total phosphorus (TP) |
| | | Contemporary nutrient | $NO_x$.sedi | Sediment $NO_x^-$ |
| | | Contemporary nutrient | $NO_2$.sedi | Sediment $NO_2^-$ |
| | | Contemporary nutrient | $NH_4$.sedi | Sediment $NH_4^+$ |
| | | Contemporary nutrient | $PO_4$.sedi | Sediment $PO_4^{3-}$ |
| | | Contemporary nutrient | $NO_3$ | Water $NO_3^-$ |
| | | Contemporary nutrient | $NO_2$ | Water $NO_2^-$ |
| | | Contemporary nutrient | $NH_4$ | Water $NH_4^+$ |
| | | Contemporary nutrient | $PO_4$ | Water $PO_4^{3-}$ |
| Proximal drivers | Energy supply | Energy supply | TC.sedi | Sediment total organic carbon (TOC) |
| | | Energy supply | DOC.sedi | Sediment dissolved organic carbon (DOC) |
| | | Energy supply | pH | Water pH |
| | | Energy supply | Chla.sedi | Sediment Chlorophyll *a* (Chl *a*) |
| | Diversity | Biodiversity | bac.rich | Species richness of bacteria |
| | | Biodiversity | bac.nmds1 | NMDS axis 1 of bacterial composition |
| | | Biodiversity | bac.nmds2 | NMDS axis 2 of bacterial composition |
| | | Chemodiversity | chemo.rich | Chemical richness of molecules |
| | | Chemodiversity | mol.nmds1 | NMDS axis 1 of molecular composition |
| | | Chemodiversity | mol.nmds2 | NMDS axis 2 of molecular composition |
| | Molecular traits | Molecular weight | Mass | The mass to charge ratio (m/z) |
| | | Molecular weight | C | The number of carbon |
| | | Molecular weight | $kdefect_{CH2}$ | Kendrick Defect |
| | | Stoichiometry | O/C | O/C ratio |
| | | Stoichiometry | H/C | H/C ratio |
| | | Stoichiometry | N/C | N/C ratio |
| | | Stoichiometry | P/C | P/C ratio |
| | | Stoichiometry | N/P | N/P ratio |

| | | | |
|---|---|---|---|
| | Stoichiometry | S/C | S/C ratio |
| | Chemical structure | $AI_{Mod}$ | The modified aromaticity index |
| | Chemical structure | DBE | Double bond equivalence |
| | Chemical structure | $DBE_O$ | Double bond equivalence minus oxygen |
| | Chemical structure | $DBE_{AI}$ | Double bond equivalence minus aromaticity index |
| | Oxidation state | GFE | Gibbs free energy |
| | Oxidation state | NOSC | Nominal oxidation state of carbon |
| | Carbon use efficiency | $Y_{met}$ | Carbon use efficiency |

[#] Water temperature was used to represent climatic variables due to its strong associations with elevation in both mountain ranges (Wang *et al*. 2016).

**Table S2**. Formulae to calculate composite variables for structure equation models of the specialization $H_2$’ of DOM-microbe associations. We constructed four bipartite networks, that is, the negative and positive networks in China or Norway. The obtained composite variables were used in Fig. 5. The abbreviations of included variables are listed in Table S1.

| Response | Region | Network type | Composite | Formula |
|---|---|---|---|---|
| $H_2$' | China | Negative | Contemporary nutrient | -0.606 × $NO_3$ + -0.222 × $NO_2$ + 0.221 × $PO_4$ + -0.294 × TN.sedi + 0.184 × $PO_4$.sedi + 0.296 × $NH_4$.sedi + 0.219 × $NO_2$.sedi |
| | | | Energy supply | -0.589 × pH + 0.493 × DOC.sedi |
| | | | Biodiversity | -0.278 × bac.nmds1 + 0.160 × bac.nmds2 |
| | | | Chemodiversity | -0.451 × chemo.rich + 0.481 × mol.nmds1 |
| | | | Molecular traits | 66.076 × Mass + -64.710 × C + -1.402 × $AI_{Mod}$ + -15.166 × DBE + -20.154 × $DBE_O$ + 61.402 × $DBE_{AI}$ + -3.279 × GFE + -23.908 × $kdefect_{CH2}$ + -1.378 × O/C + 2.627 × H/C + 0.652 × N/P |
| $H_2$' | China | Positive | Contemporary nutrient | 0.339 × $NO_2$ + -0.442 × $PO_4$ + 0.443 × $PO_4$.sedi + -0.173 × NO2.sedi |
| | | | Energy supply | 0.525 × pH + 0.253 × Chla.sedi + -0.102 × TC.sedi + -0.262 × DOC.sedi |
| | | | Biodiversity | -0.293 × bac.rich + 0.718 × bac.nmds1 |
| | | | Chemodiversity | -0.110 × chemo.rich + -0.591 × mol.nmds1 + 0.416 × mol.nmds2 |
| | | | Molecular traits | -50.009 × Mass + 48.984 × C + 1.728 × $AI_{Mod}$ + 3.075 × $DBE_O$ + -26.770 × $DBE_{AI}$ + 3.904 × GFE + 24.822 × $kdefect_{CH2}$ + 1.204 × O/C + -5.311 × H/C |
| $H_2$' | Norway | Negative | Contemporary nutrient | -0.381 × $NO_3$ + 0.365 × $PO_4$.sedi |
| | | | Energy supply | 0.210 × DOC.sedi |
| | | | Chemodiversity | -0.359 × chemo.rich + -0.186 × mol.nmds1 + -0.120 × mol.nmds2 |
| | | | Molecular traits | -1.316 × Mass + 1.062 × $AI_{Mod}$ + 3.571 × DBE + 2.524 × $DBE_O$ + -4.638 × $DBE_{AI}$ + 2.193 × H/C + -0.387 × N/C + -0.372 × N/P |
| $H_2$' | Norway | Positive | Contemporary nutrient | -0.338 × $NO_3$ + 0.280 × $NO_2$ + 0.338 × $PO_4$ + 0.350 × $PO_4$.sedi + -0.105 × $NO_2$.sedi |
| | | | Energy supply | -0.327 × pH + 0.231 × TC.sedi + -0.350 × DOC.sedi |
| | | | Biodiversity | -0.510 × bac.nmds1 |
| | | | Chemodiversity | -0.169 × mol.nmds1 + -0.480 × mol.nmds2 |
| | | | Molecular traits | 1.872 × Mass + -1.825 × DBE + 2.282 × $DBE_{AI}$ + -2.674 × GFE + -1.431 × $kdefect_{CH2}$ + 1.288 × H/C + -0.877 × N/C + 1.653 × P/C |

**Table S3**. Summary of the model fit statistics evaluated for standardized structural equation model (SEM). We explored the potential links between predictor variables and the specialization $H_2$' of the negative and positive bipartite networks in China or Norway, and the best-fitting models are shown in Fig. 5. We constructed the full SEM models based on the hypothetical casual relationships (Fig. S1), and further performed sequential models by dropping non-significant paths from the full models. $\chi^2$: Chi-square. *P*: p-value of chi-square test. df: Degrees of freedom. CFI: Comparative fit index. SRMR: Standardized root mean squared residual. AICc: Second-order Akaike information criterion. ΔAICc: Delta AICc.

| SEM models | Omitted paths | df | $\chi^2$ | *P* | CFI | SRMR | AICc | ΔAICc |
|---|---|---|---|---|---|---|---|---|
| **China; Negative** | | | | | | | | |
| 1[a] | | 1 | 0.563 | 0.453 | 1 | 0.004 | 1818.8 | 10.75 |
| 2 | Human impacts -> $H_2$' | 2 | 1.267 | 0.531 | 1 | 0.005 | 1816.6 | 8.53 |
| 3 | Human impacts -> $H_2$'<br>Human impacts -> Chemodiversity | 4 | 2.924 | 0.571 | 1 | 0.017 | 1812.5 | 4.46 |
| 4 | Human impacts -> $H_2$'<br>Human impacts -> Chemodiversity<br>Human impacts -> Biodiversity | 5 | 4.335 | 0.502 | 1 | 0.017 | 1811.2 | 3.08 |
| 5 | Human impacts -> $H_2$'<br>Human impacts -> Chemodiversity<br>Human impacts -> Biodiversity<br>Human impacts -> Energy | 6 | 6.201 | 0.401 | 1 | 0.017 | 1810.3 | 2.19 |
| 6 | Human impacts -> $H_2$'<br>Human impacts -> Chemodiversity<br>Human impacts -> Biodiversity<br>Human impacts -> Energy<br>Nutrient -> DOM traits | 7 | 8.631 | 0.280 | 1 | 0.018 | 1810.0 | 1.92 |
| 7 | Human impacts -> $H_2$'<br>Human impacts -> Chemodiversity<br>Human impacts -> Biodiversity<br>Human impacts -> Energy<br>Nutrient -> DOM traits<br>Biodiversity -> DOM traits | 8 | 10.579 | 0.227 | 1 | 0.020 | 1809.3 | 1.20 |
| 8[b] | Human impacts -> $H_2$'<br>Human impacts -> Chemodiversity<br>Human impacts -> Biodiversity<br>Human impacts -> Energy<br>Nutrient -> DOM traits<br>Biodiversity -> DOM traits<br>Climate change -> DOM traits | 7 | 6.714 | 0.459 | 1 | 0.018 | 1808.1 | 0 |
| **China; Positive** | | | | | | | | |
| 1[a] | | 1 | 0.760 | 0.383 | 1 | 0.004 | 1966.3 | 15.04 |

| | | | | | | | | |
|---|---|---|---|---|---|---|---|---|
| 2 | Nutrient -> Biodiversity | 2 | 0.768 | 0.681 | 1 | 0.004 | 1963.4 | 12.11 |
| 3 | Nutrient -> Biodiversity<br>Energy -> $H_2$' | 3 | 0.869 | 0.833 | 1 | 0.004 | 1960.6 | 9.33 |
| 4 | Nutrient -> Biodiversity<br>Energy -> $H_2$'<br>Human impacts -> $H_2$' | 4 | 1.086 | 0.896 | 1 | 0.005 | 1958.0 | 6.71 |
| 5 | Nutrient -> Biodiversity<br>Energy -> $H_2$'<br>Human impacts -> $H_2$'<br>Nutrient -> DOM traits | 5 | 1.477 | 0.916 | 1 | 0.006 | 1955.6 | 4.31 |
| 6 | Nutrient -> Biodiversity<br>Energy -> $H_2$'<br>Human impacts -> $H_2$'<br>Nutrient -> DOM traits<br>Human impacts -> Biodiversity | 6 | 2.526 | 0.866 | 1 | 0.009 | 1953.9 | 2.61 |
| 7 | Nutrient -> Biodiversity<br>Energy -> $H_2$'<br>Human impacts -> $H_2$'<br>Nutrient -> DOM traits<br>Human impacts -> Biodiversity<br>Climate change -> $H_2$' | 7 | 3.888 | 0.793 | 1 | 0.010 | 1952.6 | 1.26 |
| 8[b] | Nutrient -> Biodiversity<br>Energy -> $H_2$'<br>Human impacts -> $H_2$'<br>Nutrient -> DOM traits<br>Human impacts -> Biodiversity<br>Climate change -> $H_2$'<br>Climate change -> Chemodiversity | 8 | 5.291 | 0.726 | 1 | 0.013 | 1951.3 | 0 |
| 9 | Nutrient -> Biodiversity<br>Energy -> $H_2$'<br>Human impacts -> $H_2$'<br>Nutrient -> DOM traits<br>Human impacts -> Biodiversity<br>Climate change -> $H_2$'<br>Climate change -> Chemodiversity<br>Human impacts -> DOM traits | 9 | 8.220 | 0.512 | 1 | 0.019 | 1951.6 | 0.30 |
| 10 | Nutrient -> Biodiversity<br>Energy -> $H_2$'<br>Human impacts -> $H_2$'<br>Nutrient -> DOM traits<br>Human impacts -> Biodiversity<br>Climate change -> $H_2$'<br>Climate change -> Chemodiversity<br>Human impacts -> DOM traits<br>Nutrient -> $H_2$' | 10 | 11.134 | 0.347 | 1 | 0.023 | 1951.9 | 0.63 |
| **Norway; Negative** | | | | | | | | |
| 1[a] | | 0 | 0 | 0 | 1 | 0 | 1225.2 | 11.51 |
| 2 | Energy -> Chemodiversity | 1 | 0.120 | 0.729 | 1 | 0.004 | 1222.6 | 8.96 |

| | | | | | | | | |
|---|---|---|---|---|---|---|---|---|
| 3 | Energy -> Chemodiversity<br>Climate change -> Energy | 2 | 0.263 | 0.877 | 1 | 0.006 | 1220.1 | 6.48 |
| 4 | Energy -> Chemodiversity<br>Climate change -> Energy<br>Energy -> $H_2$' | 3 | 0.656 | 0.884 | 1 | 0.007 | 1218.0 | 4.29 |
| 5 | Energy -> Chemodiversity<br>Climate change -> Energy<br>Energy -> $H_2$'<br>Nutrient -> DOM traits | 4 | 1.097 | 0.895 | 1 | 0.011 | 1215.8 | 2.18 |
| 6 | Energy -> Chemodiversity<br>Climate change -> Energy<br>Energy -> $H_2$'<br>Nutrient -> DOM traits<br>Climate change -> Nutrient | 5 | 2.097 | 0.836 | 1 | 0.022 | 1214.3 | 0.67 |
| 7[b] | Energy -> Chemodiversity<br>Climate change -> Energy<br>Energy -> $H_2$'<br>Nutrient -> DOM traits<br>Climate change -> Nutrient<br>Climate change -> $H_2$' | 6 | 3.893 | 0.691 | 1 | 0.024 | 1213.7 | 0 |
| 8 | Energy -> Chemodiversity<br>Climate change -> Energy<br>Energy -> $H_2$'<br>Nutrient -> DOM traits<br>Climate change -> Nutrient<br>Climate change -> $H_2$'<br>Chemodiversity -> DOM traits | 7 | 6.735 | 0.457 | 1 | 0.038 | 1214.1 | 0.41 |
| **Norway; Positive** | | | | | | | | |
| 1[a] | | 0 | 0 | 0 | 1 | 0 | 1751.2 | 19.79 |
| 2 | Human impacts -> Chemodiversity | 1 | 0.097 | 0.756 | 1 | 0.002 | 1748.3 | 16.91 |
| 3 | Human impacts -> Chemodiversity<br>Biodiversity -> DOM traits | 2 | 0.225 | 0.894 | 1 | 0.002 | 1745.5 | 14.11 |
| 4 | Human impacts -> Chemodiversity<br>Biodiversity -> DOM traits<br>Energy -> DOM traits | 3 | 0.391 | 0.942 | 1 | 0.003 | 1742.8 | 11.39 |
| 5 | Human impacts -> Chemodiversity<br>Biodiversity -> DOM traits<br>Energy -> DOM traits<br>Energy -> $H_2$' | 4 | 0.802 | 0.938 | 1 | 0.006 | 1740.4 | 8.97 |
| 6 | Human impacts -> Chemodiversity<br>Biodiversity -> DOM traits<br>Energy -> DOM traits<br>Energy -> $H_2$'<br>Climate change -> $H_2$' | 5 | 1.286 | 0.936 | 1 | 0.008 | 1738.0 | 6.66 |

| | | | | | | | | |
|---|---|---|---|---|---|---|---|---|
| 7 | Human impacts -> Chemodiversity<br>Biodiversity -> DOM traits<br>Energy -> DOM traits<br>Energy -> $H_2$'<br>Climate change -> $H_2$'<br>Human impacts -> Biodiversity | 6 | 1.892 | 0.929 | 1 | 0.009 | 1735.9 | 4.51 |
| 8 | Human impacts -> Chemodiversity<br>Biodiversity -> DOM traits<br>Energy -> DOM traits<br>Energy -> $H_2$'<br>Climate change -> $H_2$'<br>Human impacts -> Biodiversity<br>Nutrient -> Energy | 7 | 2.513 | 0.926 | 1 | 0.011 | 1733.8 | 2.43 |
| 9 | Human impacts -> Chemodiversity<br>Biodiversity -> DOM traits<br>Energy -> DOM traits<br>Energy -> $H_2$'<br>Climate change -> $H_2$'<br>Human impacts -> Biodiversity<br>Nutrient -> Energy<br>Chemodiversity -> $H_2$' | 8 | 3.949 | 0.862 | 1 | 0.012 | 1732.6 | 1.2 |
| 10[b] | Human impacts -> Chemodiversity<br>Biodiversity -> DOM traits<br>Energy -> DOM traits<br>Energy -> $H_2$'<br>Climate change -> $H_2$'<br>Human impacts -> Biodiversity<br>Nutrient -> Energy<br>Chemodiversity -> $H_2$'<br>Nutrient -> $H_2$' | 9 | 5.377 | 0.800 | 1 | 0.013 | 1731.4 | 0 |
| 11 | Human impacts -> Chemodiversity<br>Biodiversity -> DOM traits<br>Energy -> DOM traits<br>Energy -> $H_2$'<br>Climate change -> $H_2$'<br>Human impacts -> Biodiversity<br>Nutrient -> Energy<br>Chemodiversity -> $H_2$'<br>Nutrient -> $H_2$'<br>Human impacts -> DOM traits | 10 | 8.458 | 0.584 | 1 | 0.017 | 1731.9 | 0.5 |

[a] Full SEM models; [b] Best-fitting models shown in red.

**Table S4**. The hypothesized causal relationships and path coefficients in the structural equation model (Fig. S1).

| Relationship | Path coefficients |
|---|---|
| Climate change -> Nutrient | $\lambda_{nut,temp}$ |
| Human impacts -> Nutrient | $\lambda_{nut,N}$ |
| Climate change -> Energy | $\lambda_{energy,temp}$ |
| Human impacts -> Energy | $\lambda_{energy,N}$ |
| Nutrient -> Energy | $\lambda_{energy,nut}$ |
| Climate change -> Biodiversity | $\lambda_{biodiv,temp}$ |
| Human impacts -> Biodiversity | $\lambda_{biodiv,N}$ |
| Nutrient -> Biodiversity | $\lambda_{biodiv,nut}$ |
| Energy -> Biodiversity | $\lambda_{biodiv,energy}$ |
| Climate change -> Chemodiversity | $\lambda_{chemodiv,temp}$ |
| Human impacts -> Chemodiversity | $\lambda_{chemodiv,N}$ |
| Nutrient -> Chemodiversity | $\lambda_{chemodiv,nut}$ |
| Energy -> Chemodiversity | $\lambda_{chemodiv,energy}$ |
| Climate change -> DOM traits | $\lambda_{trait,temp}$ |
| Human impacts -> DOM traits | $\lambda_{trait,N}$ |
| Nutrient -> DOM traits | $\lambda_{trait,nut}$ |
| Energy -> DOM traits | $\lambda_{trait,energy}$ |
| Biodiversity -> DOM traits | $\lambda_{trait,biodiv}$ |
| Chemodiversity -> DOM traits | $\lambda_{trait,chemodiv}$ |
| Climate change -> $H_2$' | $\lambda_{H2,temp}$ |
| Human impacts -> $H_2$' | $\lambda_{H2,N}$ |
| Nutrient -> $H_2$' | $\lambda_{H2,nut}$ |
| Energy -> $H_2$' | $\lambda_{H2,energy}$ |
| Biodiversity -> $H_2$' | $\lambda_{H2,biodiv}$ |
| Chemodiversity -> $H_2$' | $\lambda_{H2,chemodiv}$ |
| DOM traits -> $H_2$' | $\lambda_{H2,trait}$ |

### Supplementary Figures

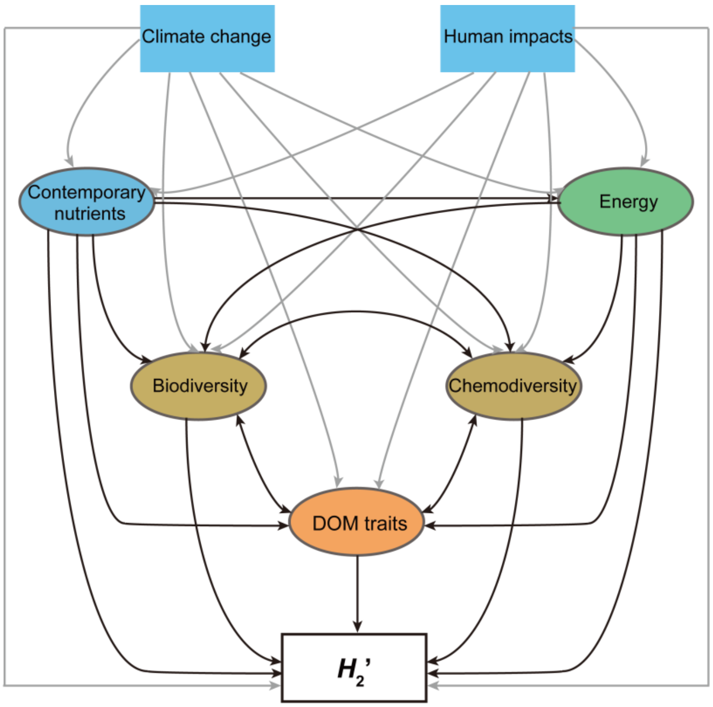


**Figure S1**. Conceptual model showing the hypothesized causal relationships among distal and proximal drivers for the specialization $H_2$' of DOM-microbe associations. The distal drivers include climate change, human impacts and contemporary nutrients, and the proximal drivers are energy supply, biodiversity, chemodiversity and DOM traits. Grey or black arrows indicate the hypothesized relationships among the exogenous or endogenous composite variables and the DOM-microbe associations, respectively.

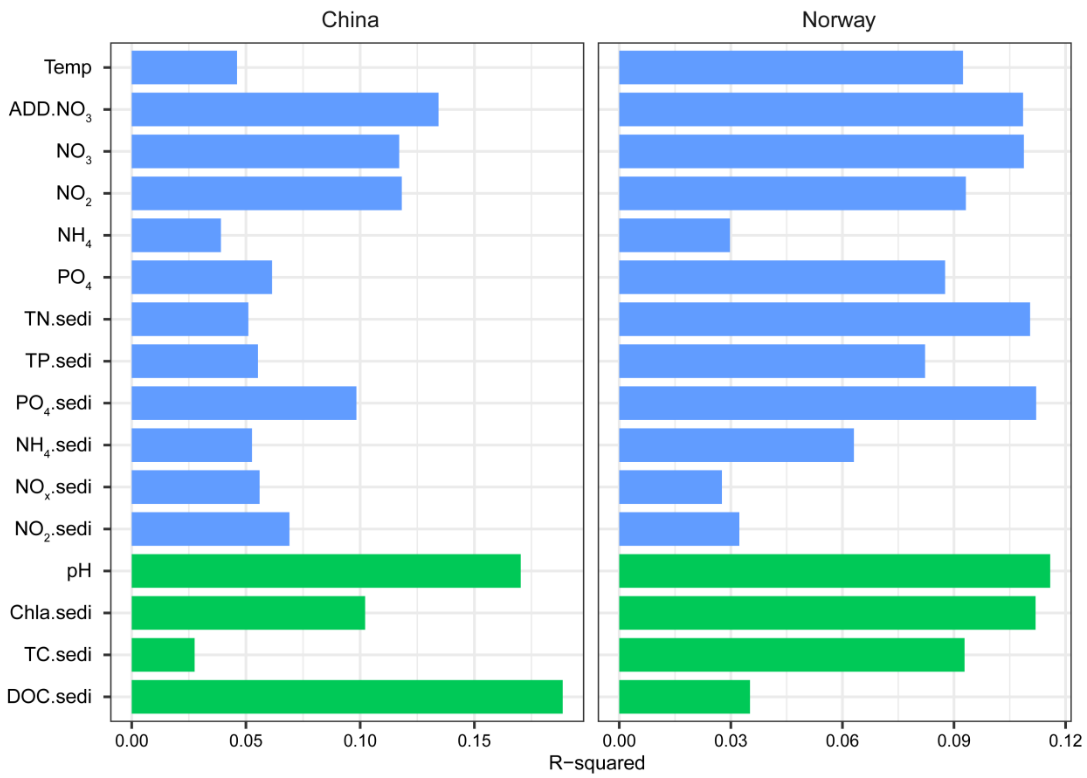


**Figure S2**. Bar plots showing R-squared to identify the effects of environmental (blue) and energy supply (green) variables on DOM composition in China and Norway. R-squared was determined by permutational multivariate analysis of variance (PERMANOVA) with 999 permutations and was statistically significant ($P \leq 0.001$) for explanatory variables. The abbreviations of explanatory variables are detailed in Table S1.

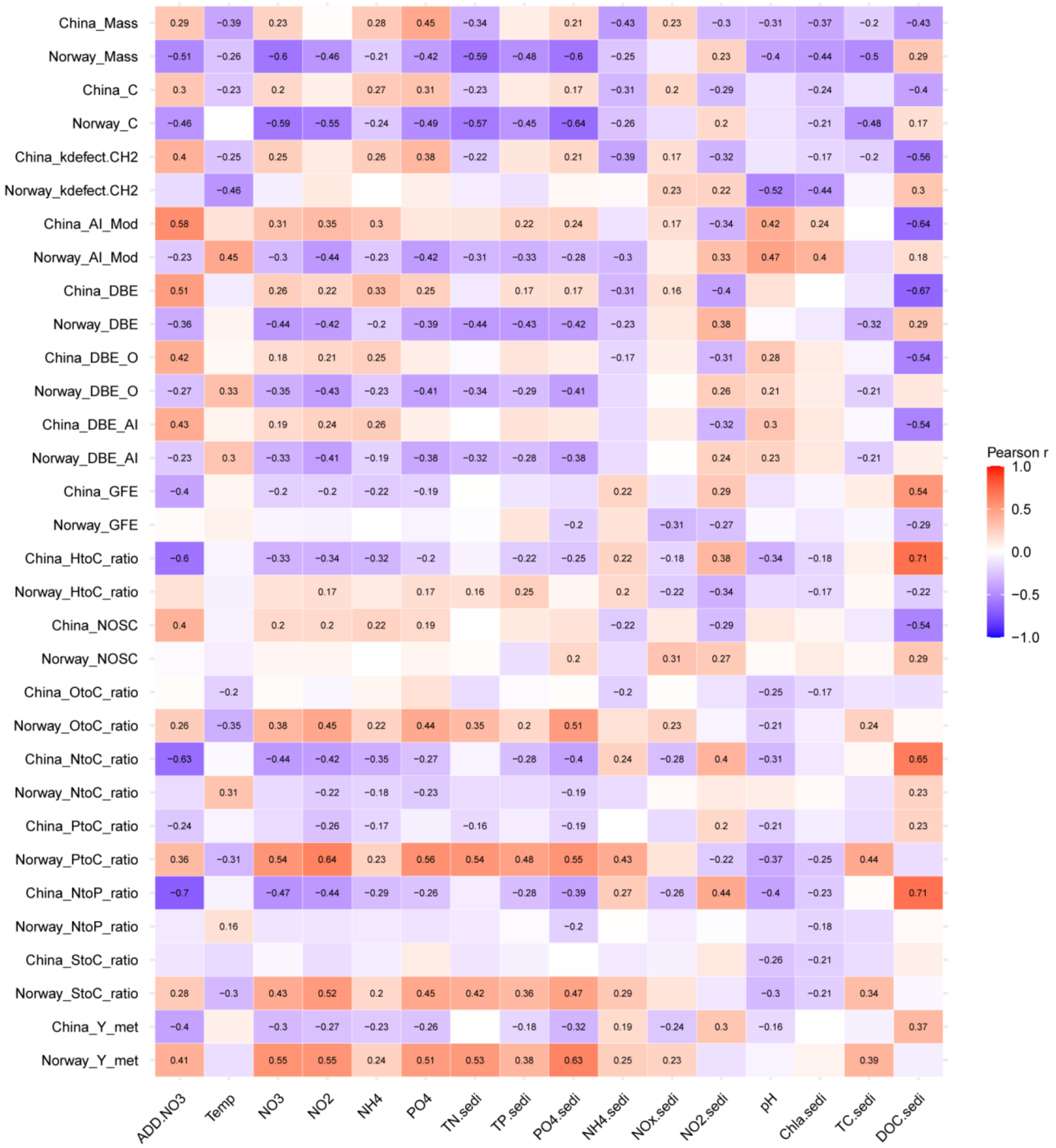


**Figure S3**. The heatmap shows the positive (red) and negative (blue) Pearson coefficients between DOM traits and explanatory variables in China and Norway. We included explanatory variables relevant to environment and energy supply (Table S1). Numbers in the heatmap indicate the significant coefficients ($P \leq 0.05$). For better visualization, weighted mean of each DOM trait in China or Norway was shortened by Country_Traits. For instance, China_Mass stands for weighted mean of molecular mass in China.

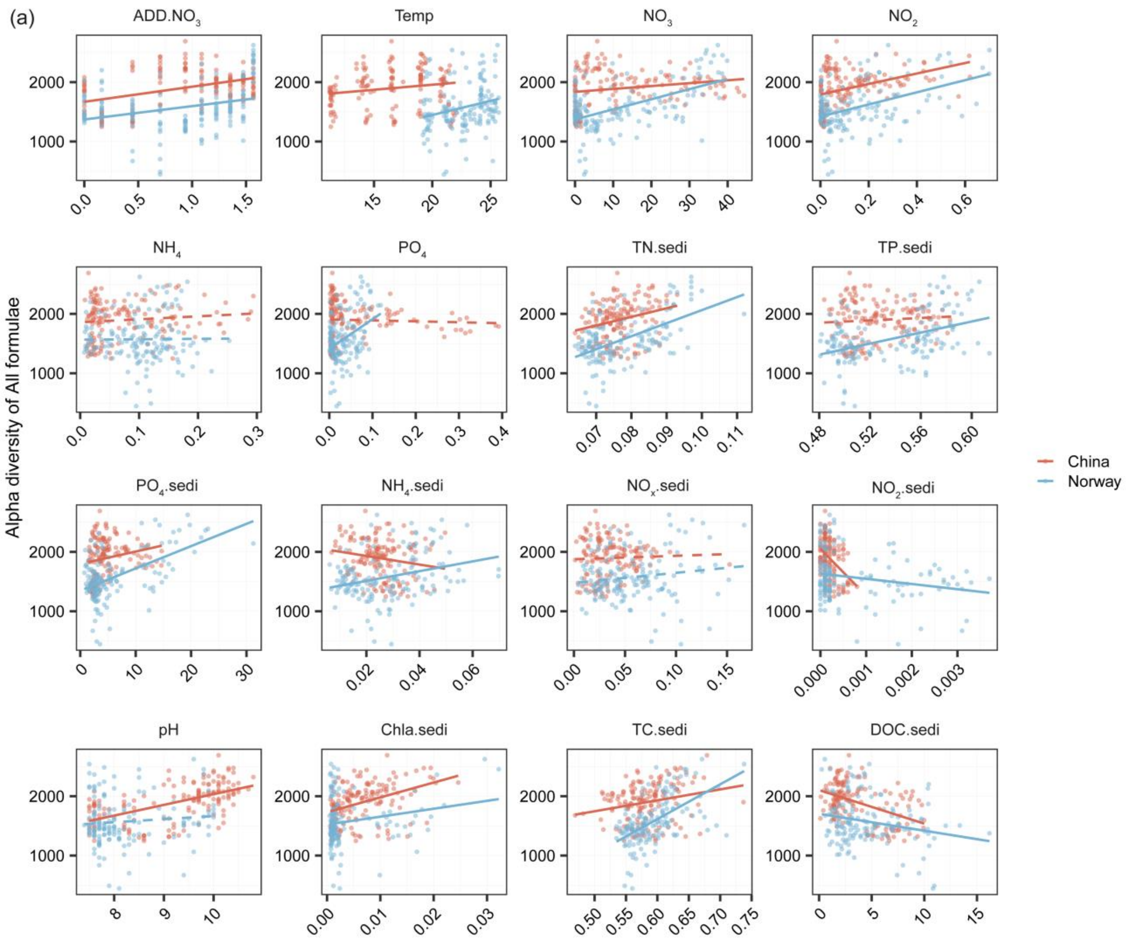


**Figure S4**. The relationships between DOM features and explanatory variables in China (red lines) and Norway (blue lines). We considered alpha diversity (i.e., molecular richness) of all molecules (a) and also molecular traits such as weighted means of mass (b) and H/C ratio (c) in China and Norway, which were plotted against the variables relevant to environment and energy supply (Table S1). The relationships are indicated by solid ($P \leq 0.05$) and dotted ($P > 0.05$) lines estimated using linear models.

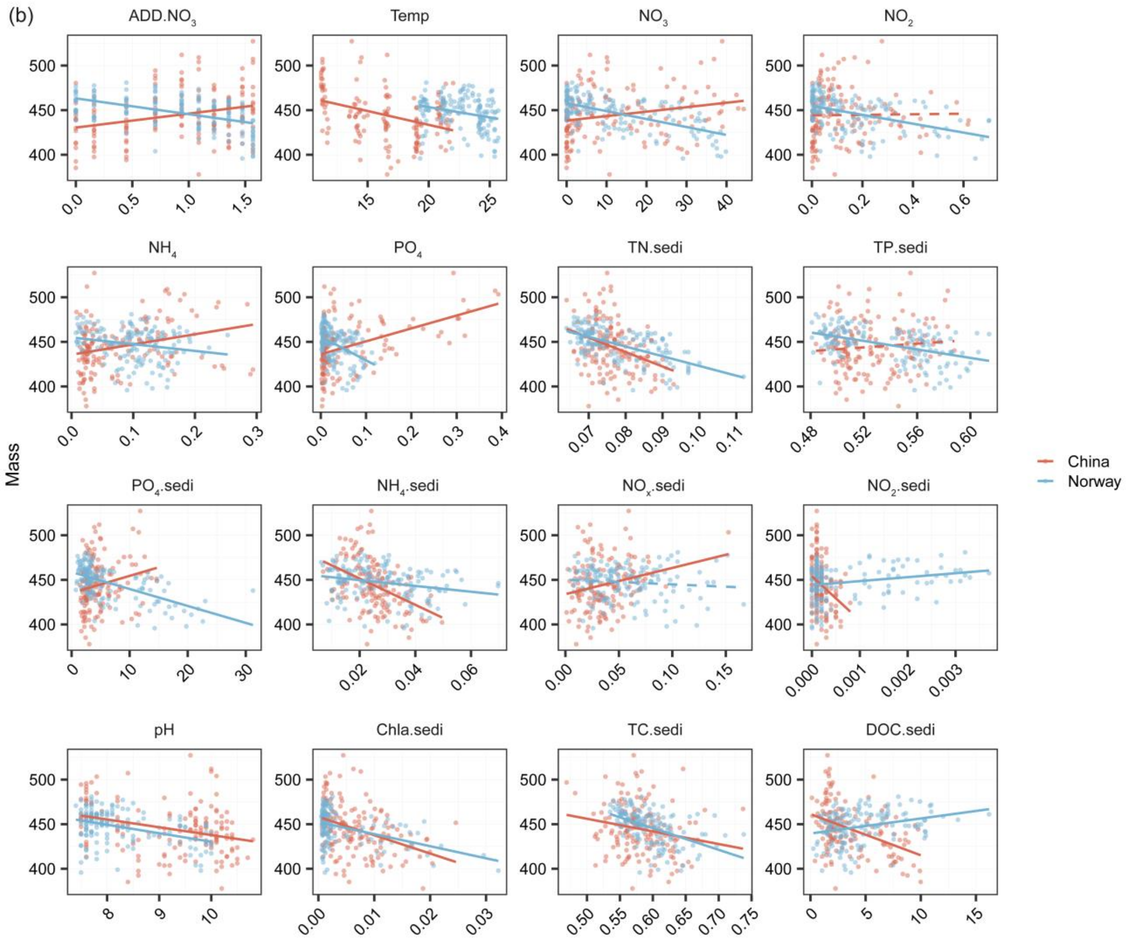


**Figure S4**. Continued. Weighted mean of mass (b).

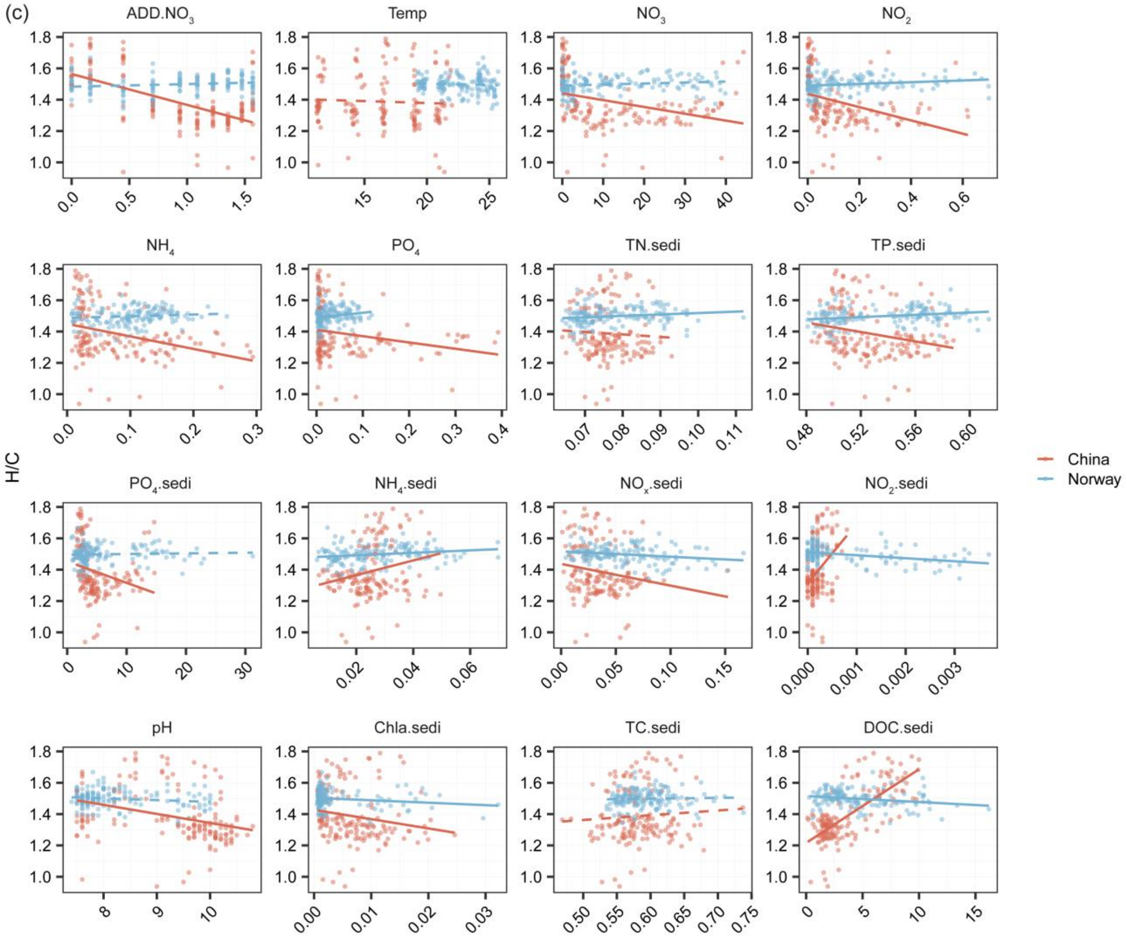


**Figure S4**. Continued. Weighted mean of H/C ratio (c).

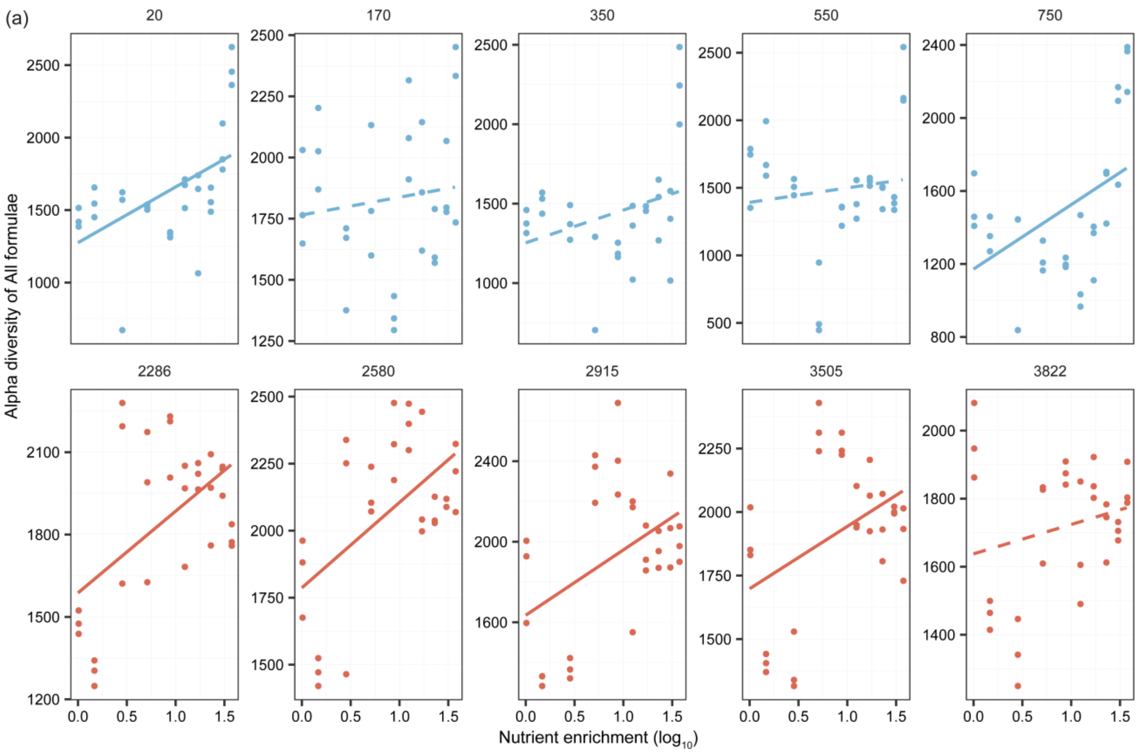


**Figure S5**. The relationships between nutrient enrichment and DOM alpha diversity or molecular traits in China (red lines) and Norway (blue lines) at different elevations (20 to 3,822 m a.s.l.). We considered richness for all formulae (a) and also molecular traits such as weighted means of mass (b), H/C ratio (c), O/C ratio (d) and $AI_{Mod}$ (e) in China (red lines and dots) and Norway (blue lines and dots). We plotted the richness or traits against the nutrient gradient of nitrate, and their relationships are indicated by solid ($P \leq 0.05$) and dotted ($P > 0.05$) lines using linear models.

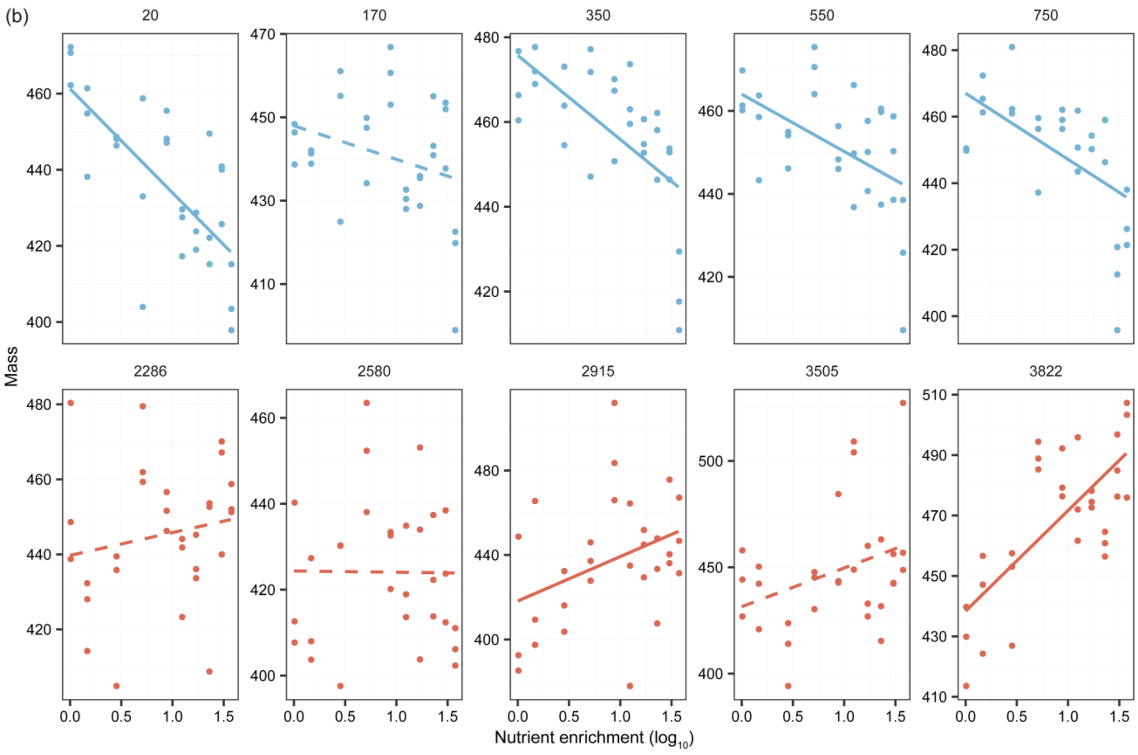


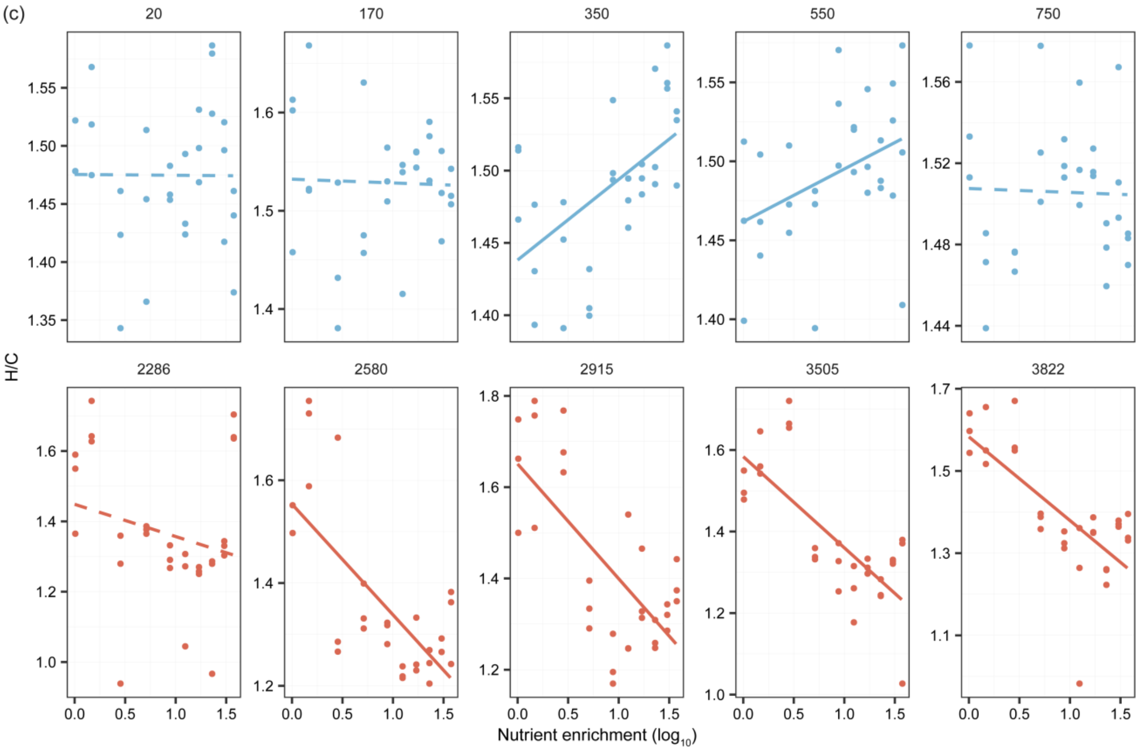


**Figure S5**. Continued. Weighted means of mass (b), and H/C ratio (c).

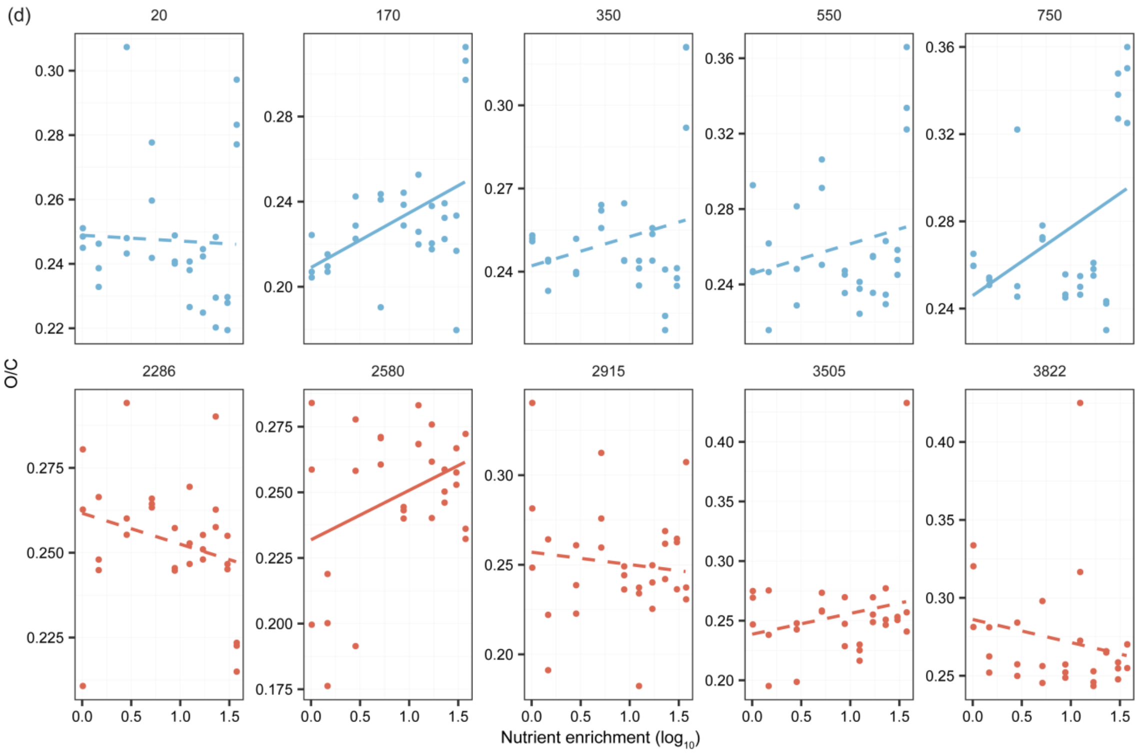


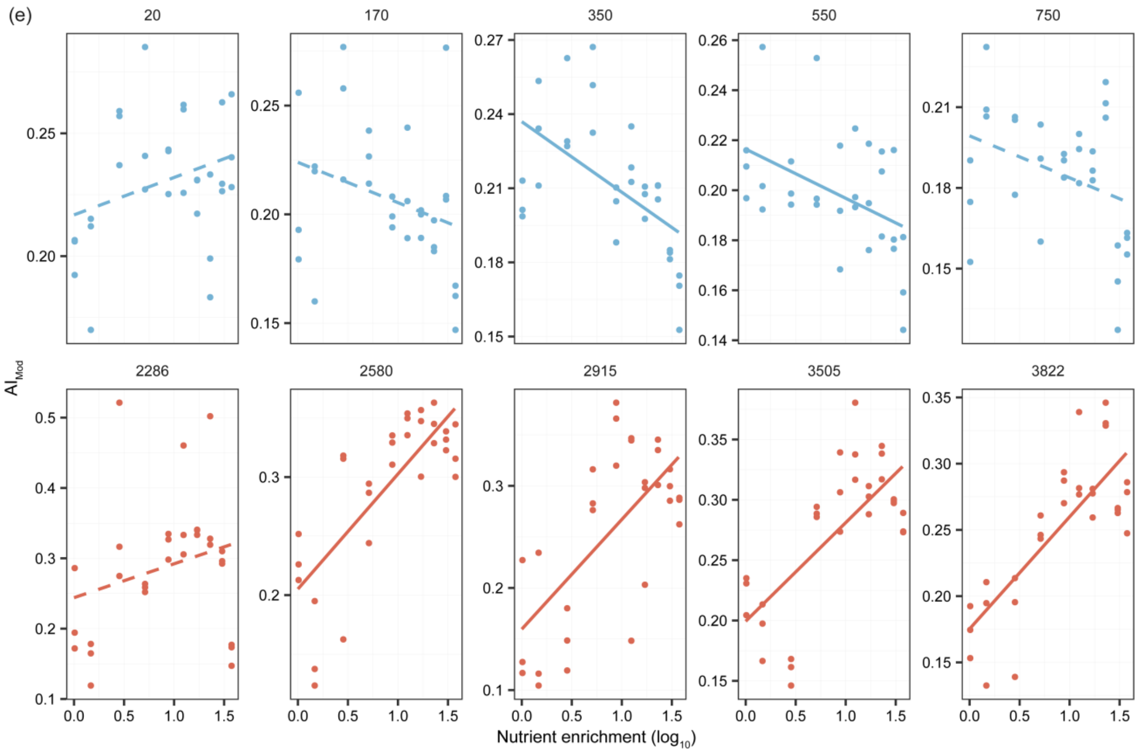


**Figure S5**. Continued. Weighted means of O/C ratio (d) and $AI_{Mod}$ (e).

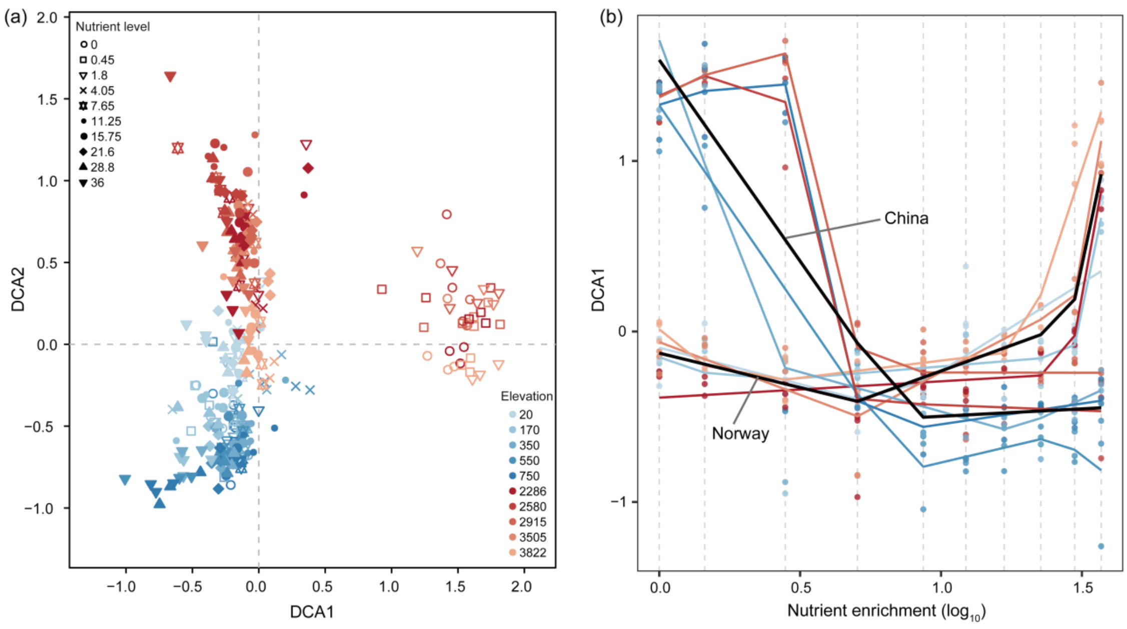


**Figure S6**. Variations in DOM compositions along the nutrient gradient of nitrate. (a) Detrended correspondence analyses (DCA) of DOM compositions. (b) Nutrient breakpoint estimation of the first axis of the DCA of DOM composition for each elevation using piecewise regression analysis (Muggeo, 2008). Black lines indicate regions, and the colored dots or lines indicate the elevations of the two regions, which are consistent with the figure legend of Fig. S6a. The vertical gray lines indicate the ten nutrient levels. We found that there were significant breakpoints ($P \leq 0.05$) of the first axis of the DCA mostly occurring between 1.80 and 4.05 mg N $L^{-1}$ along the nutrient gradient especially in China.

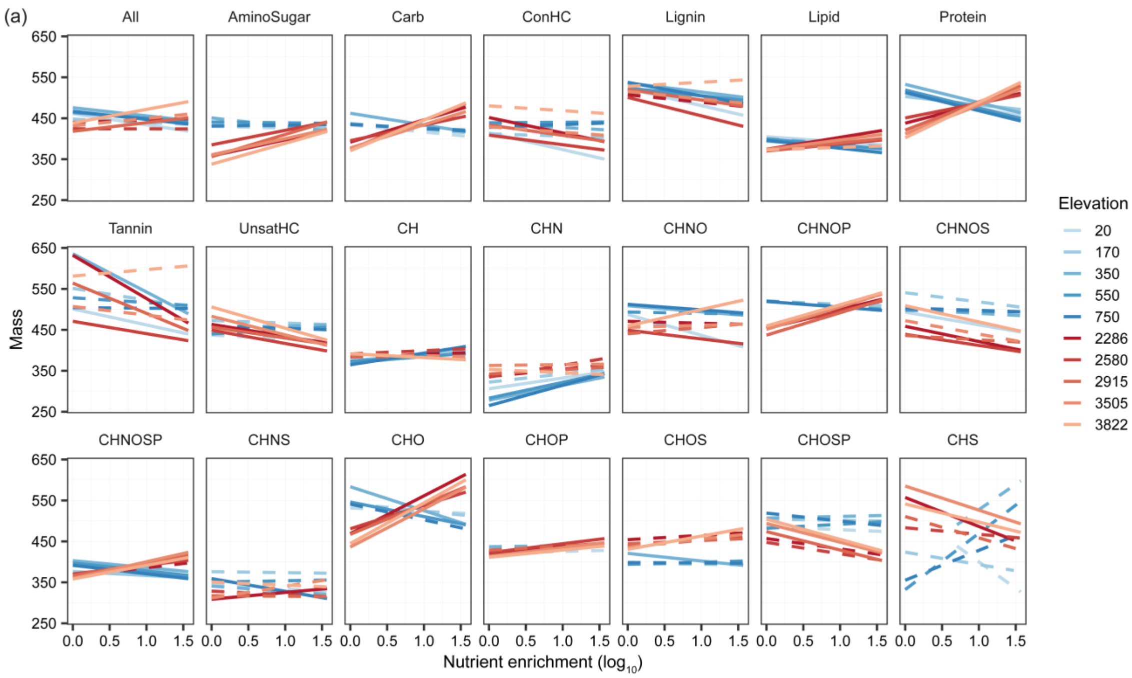


**Figure S7**. Effects of nutrient enrichment on DOM traits for all formulae and subsets of formulae within compound classes or elemental combinations across different elevations in China (red lines) and Norway (blue lines). We considered molecular traits such as weighted means of mass (a), H/C ratio (b) and $AI_{Mod}$ (c), which were plotted against the nutrient gradient of nitrate, and their relationships are indicated by the solid ($P \leq 0.05$) or dotted ($P > 0.05$) lines using linear models. The details of abbreviations of DOM traits are available in Table S1. We found that nutrient enrichment increased the weighted means of molecular mass more strongly at higher elevations in China (with maximal 495 Da at 3,822 m a.s.l.), but decreased more strongly at lower elevations in Norway (with minimal 405 Da at 20 m a.s.l.). This finding implies that nutrient enrichment leads to an increase in the molecular mass especially at colder temperatures in subtropical regions, but a decline at the warmer temperatures in subarctic regions.

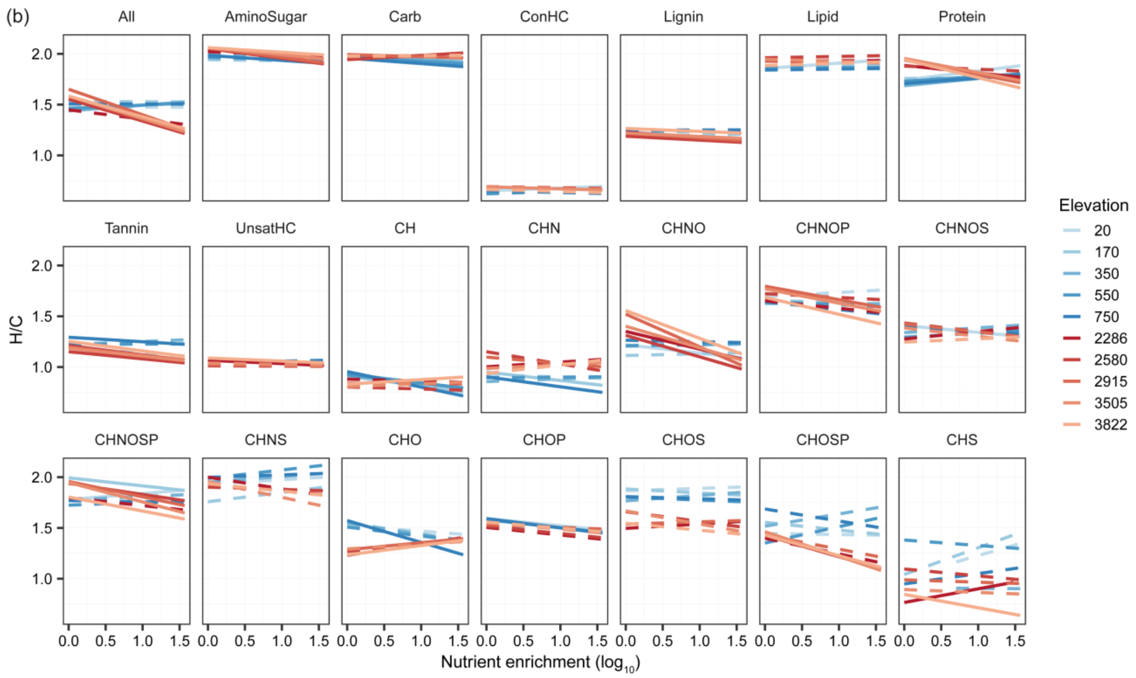


**Figure S7**. Continued. Weighted means of H/C ratio (b).

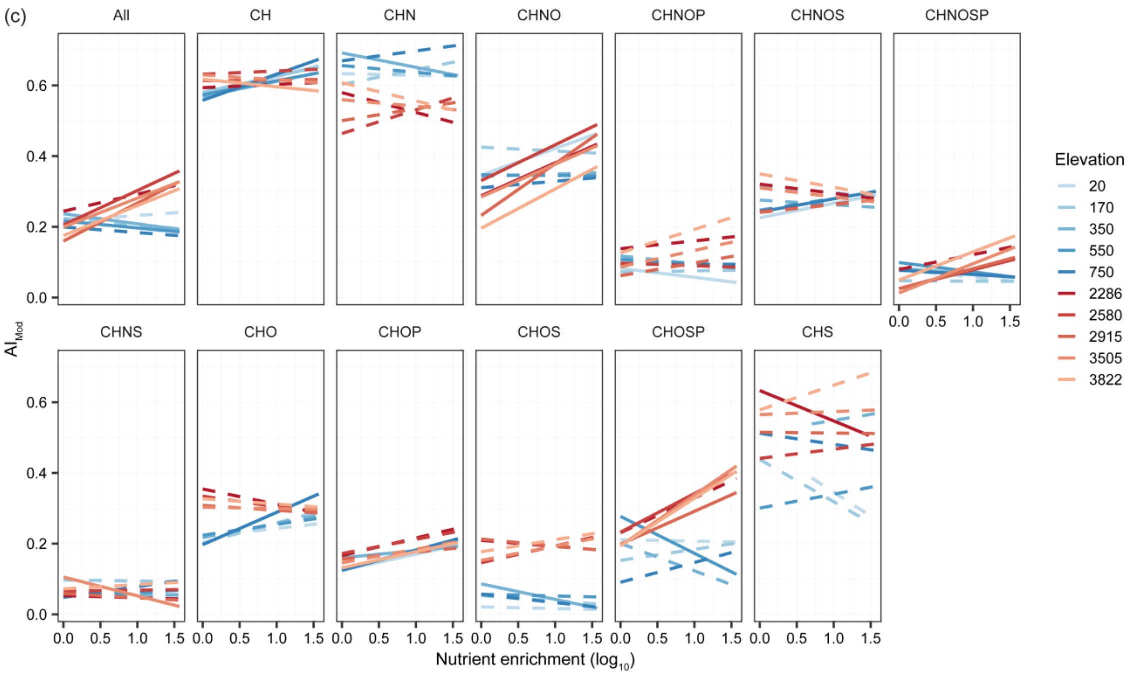


**Figure S7**. Continued. Weighted means of $AI_{Mod}$ (c).

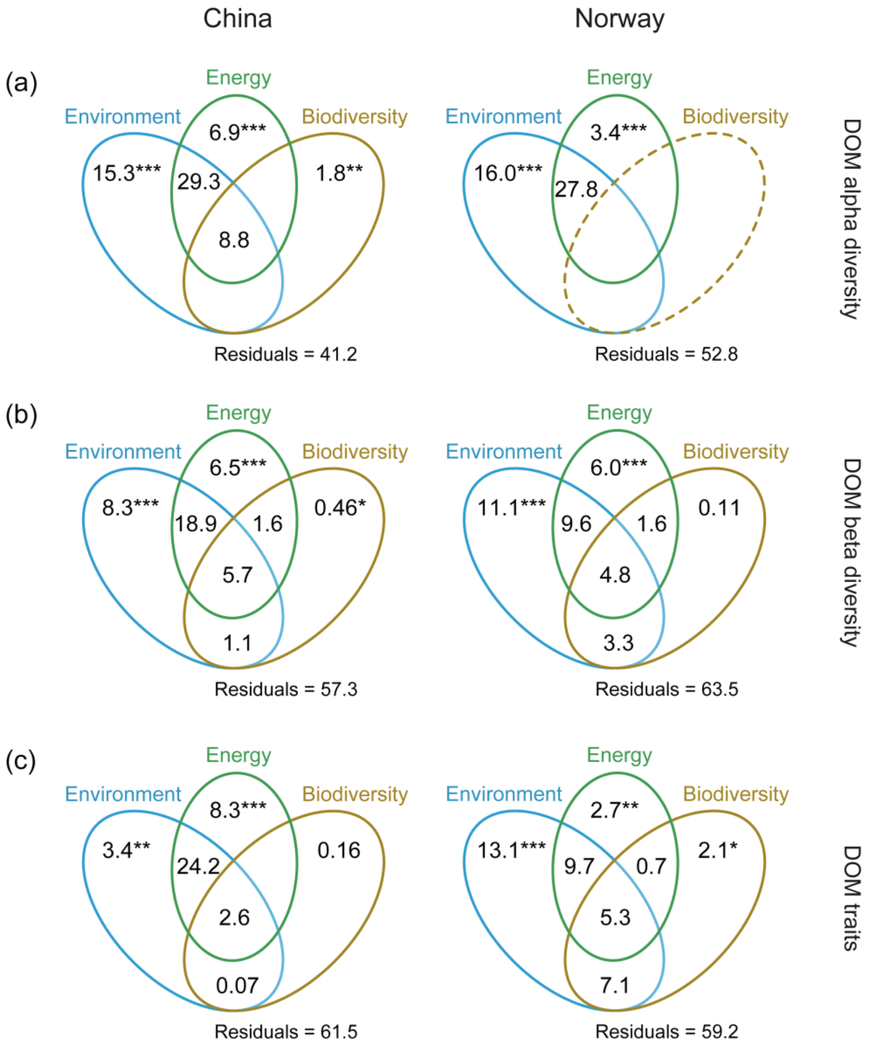


**Figure S8**. The roles of microbes in explaining the alpha diversity (upper panel), beta diversity (middle panel) and molecular traits (lower panel) of DOM using variation partitioning analysis. The numbers indicate the variance explained (%) by environments, energy supply and bacterial biodiversity which were described in detail in Table S1. Asterisks represent statistically significant effects at $^{***}$, $P \leq 0.001$; $^{**}$, $P \leq 0.01$; $^{*}$, $P \leq 0.05$.

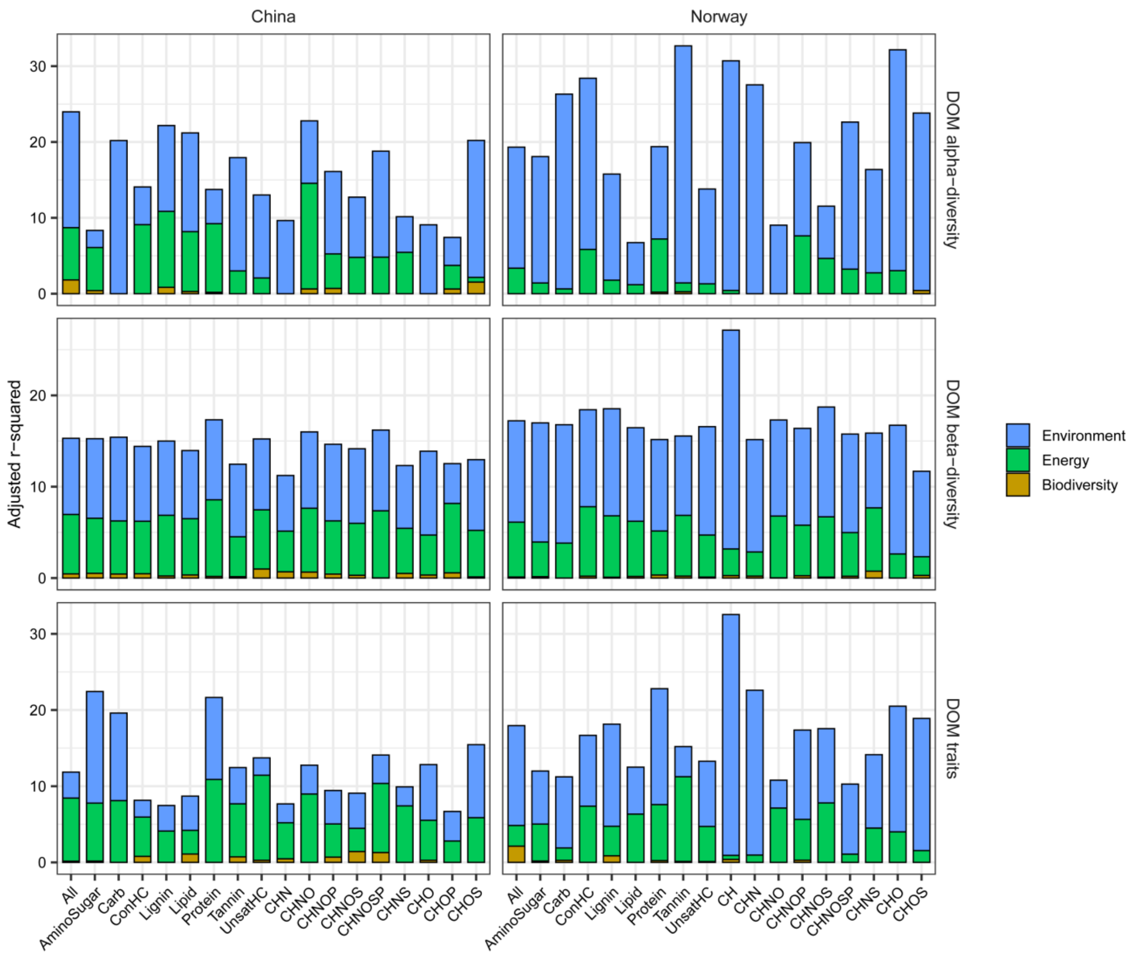


**Figure S9**. Variations (%) in DOM features purely explained by environment, energy supply and bacterial biodiversity determined by variation partitioning analysis. The DOM features were examined from three aspects: alpha diversity (upper panel), beta diversity (middle panel) and molecular traits (lower panel) for all formulae and subsets of formulae within compound classes or elemental combinations in China and Norway. Explanatory variables are described in detail in Table S1. We found that bacterial biodiversity including alpha and beta diversity had statistically significant ($P \leq 0.05$) pure effects on DOM features across all molecules (All) and the compound classes or elemental combinations, although environment and energy supply variables still had the dominant effects. For example, bacterial diversity purely explained 0-1.8%, 0-1.0% and 0-2.1% of variations in DOM alpha diversity, beta diversity and molecular traits, respectively, across different combinations of molecular composition.

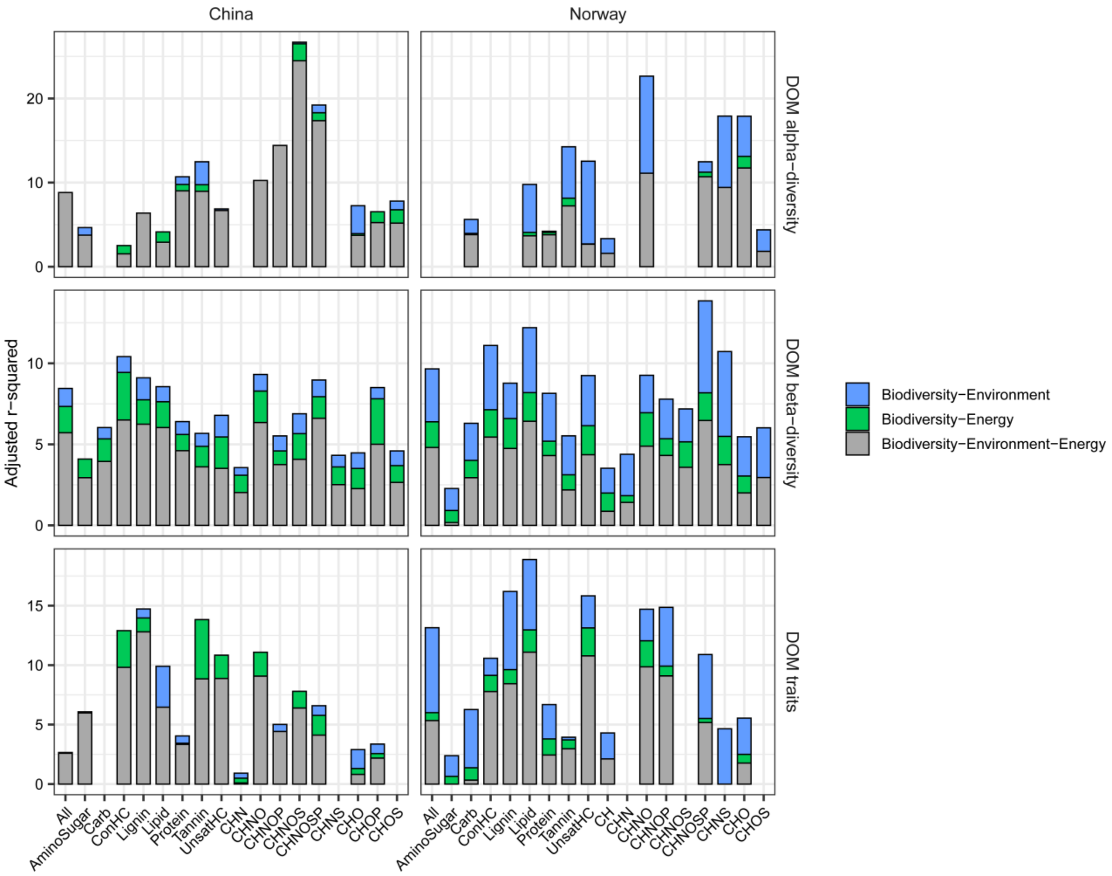


**Figure S10**. Variations (%) in DOM features explained by bacterial biodiversity jointly with other explanatory variables on DOM features determined by variation partitioning analysis. The DOM features were examined from three aspects: alpha diversity (upper panel), beta diversity (middle panel) and molecular traits (lower panel) for all formulae and subsets of formulae within compound classes or elemental combinations in China and Norway. Other explanatory variables were related to environments and/or energy supply, and are described in detail in Table S1. We found that bacterial diversity showed large shared effects with environments and energy supply on DOM features, accounting for 2.5-26.7%, 2.2-13.9% and 1.0-18.9% of the variations in DOM alpha diversity, beta diversity and molecular traits, respectively, across different combinations of molecular composition.

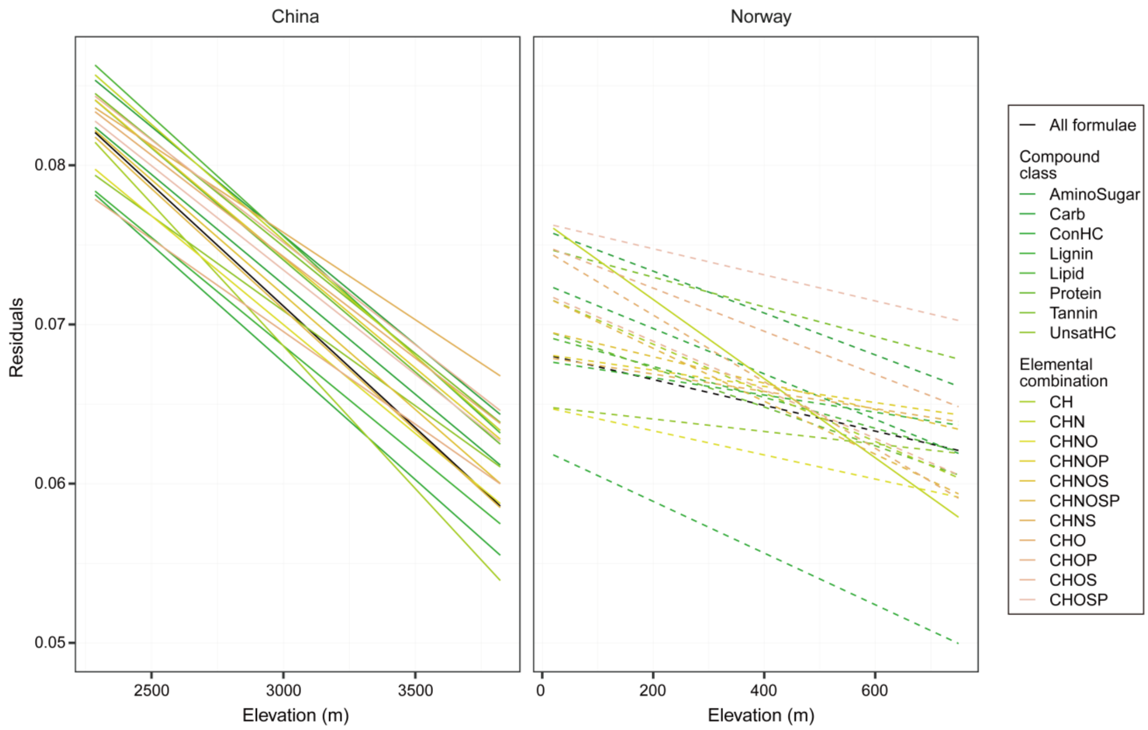


**Figure S11**. The effects of elevation on DOM-microbe associations indicated by Procrustes residuals, that is, the difference in composition between DOM and bacteria for each microcosm. These effects are indicated by solid ($P \leq 0.05$) or dotted ($P > 0.05$) lines estimated using linear models. The colours of the lines indicate the DOM composition for all formulae and categories of compound classes or elemental combinations.

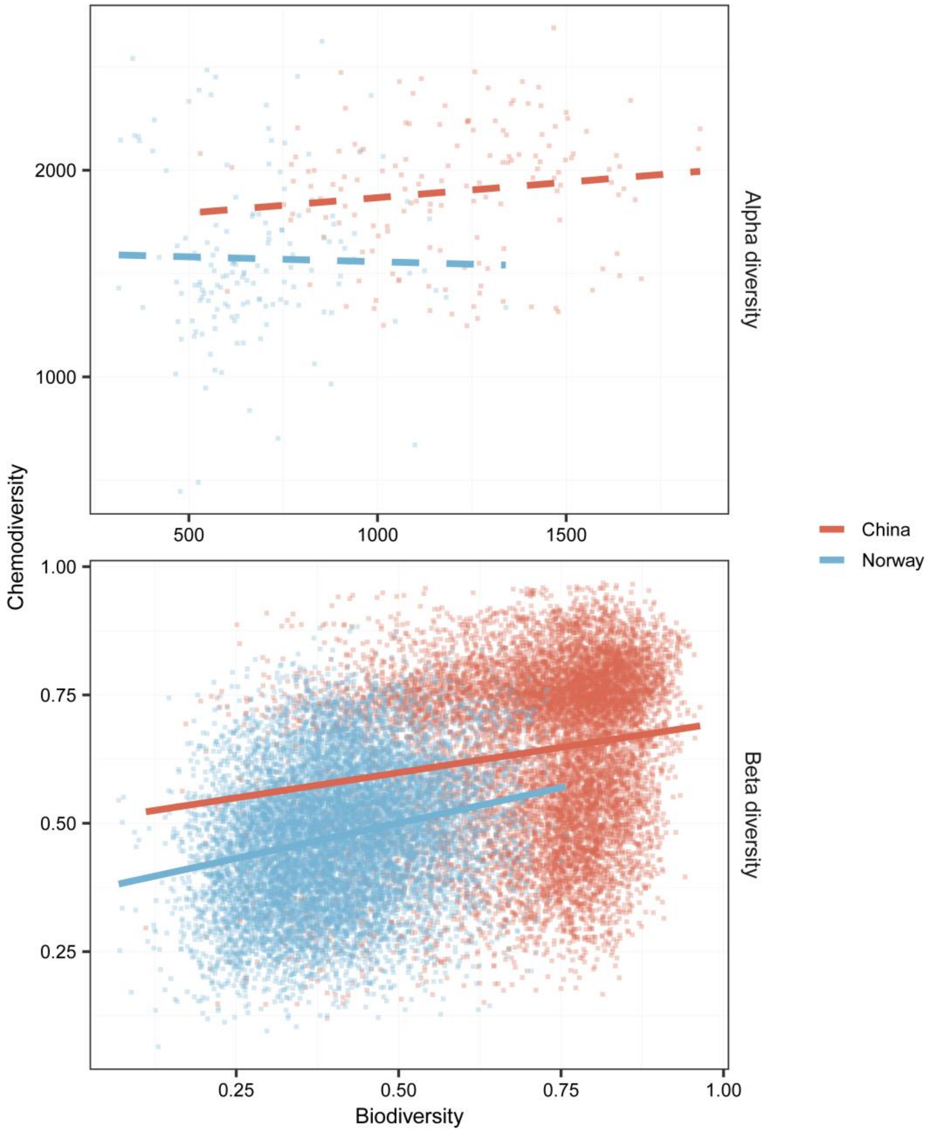


**Figure S12**. The relationships between chemodiversity and bacterial diversity in China (red lines) and Norway (blue lines). Upper panel: Alpha diversity (richness) of molecular formulae and bacterial OTUs. Lower panel: Beta diversity determined by the Bray–Curtis similarity index for mixtures of molecular formulae and communities of OTUs. The relationships are indicated by solid ($P \leq 0.05$) and dotted ($P > 0.05$) lines using linear models, and the significance was determined by ANOVA or Mantel test.

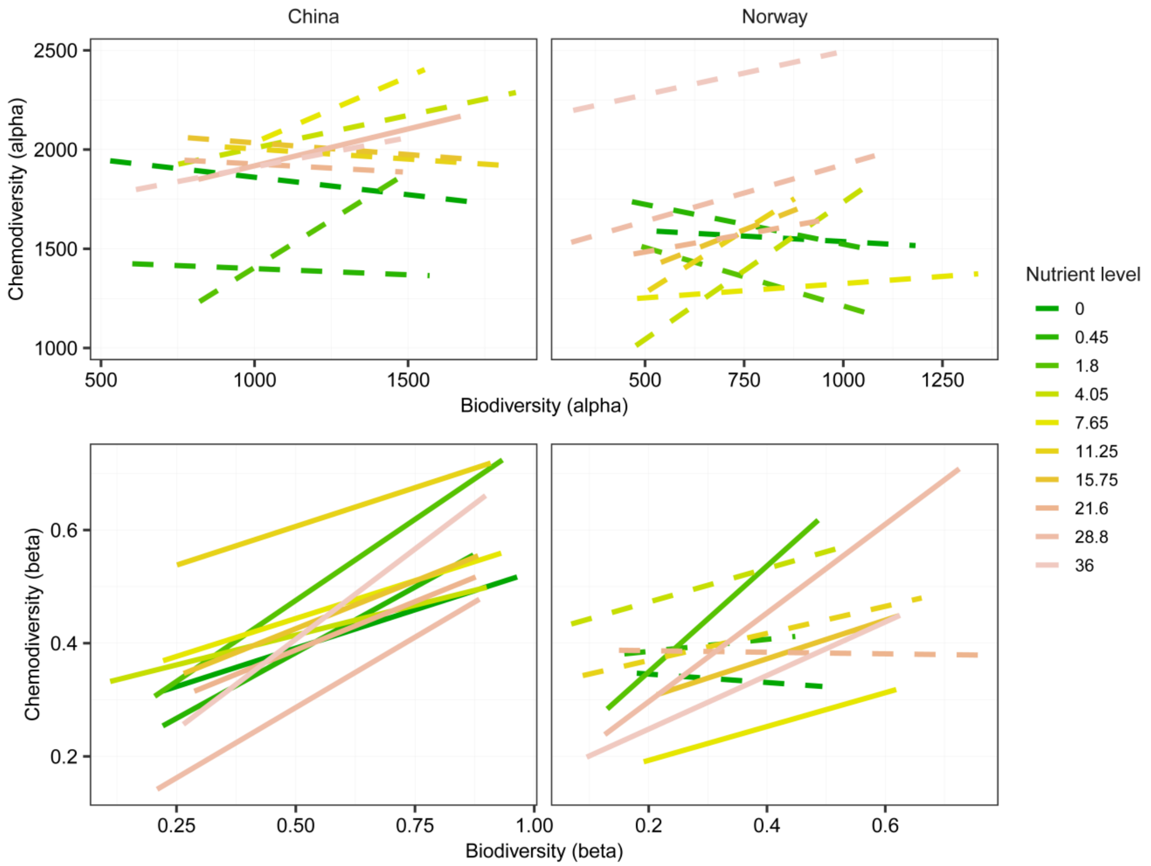


**Figure S13**. The relationships between chemodiversity and bacterial diversity in China and Norway along the nutrient gradient of nitrate. Upper panel: Alpha diversity (richness) of molecular formulae and bacterial OTUs. Lower panel: Beta diversity determined by the Bray–Curtis similarity index for mixtures of molecular formulae and communities of OTUs. The relationships are indicated by solid ($P \leq 0.05$) and dotted ($P > 0.05$) lines using linear models, and the significance was determined by ANOVA or Mantel test.

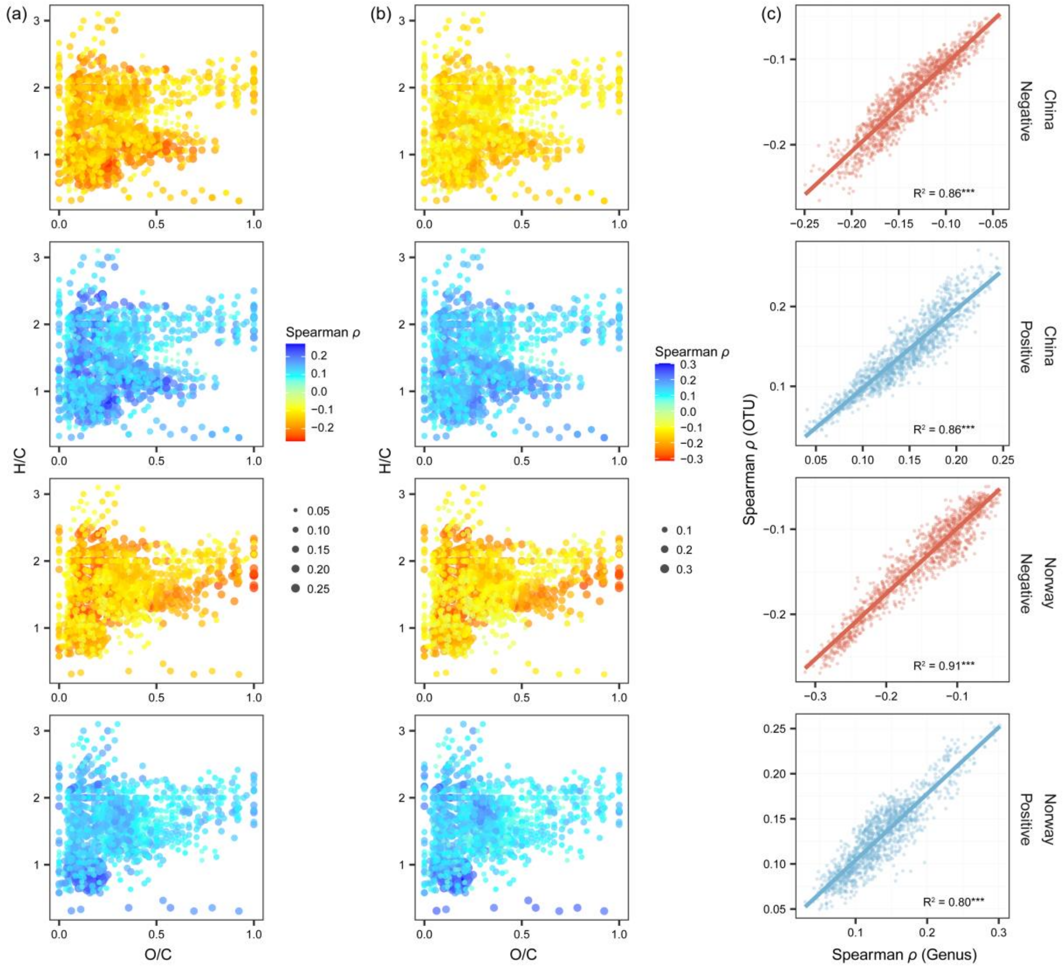


**Figure S14**. Molecular formulae correlating with bacterial OTUs (a) or genera (b) in China and Norway using Spearman's rank correlation ($\rho$). Each molecule is colored by the mean $\rho$ value of negative or positive correlations across all bacterial OTUs, and the absolute value of mean $\rho$ is indicated by the dot size. (c) The relationships in Spearman $\rho$ between bacterial OTU and genus levels for both negative and positive correlations in China and Norway. Solid lines indicate significant linear fits ($P \leq 0.05$).

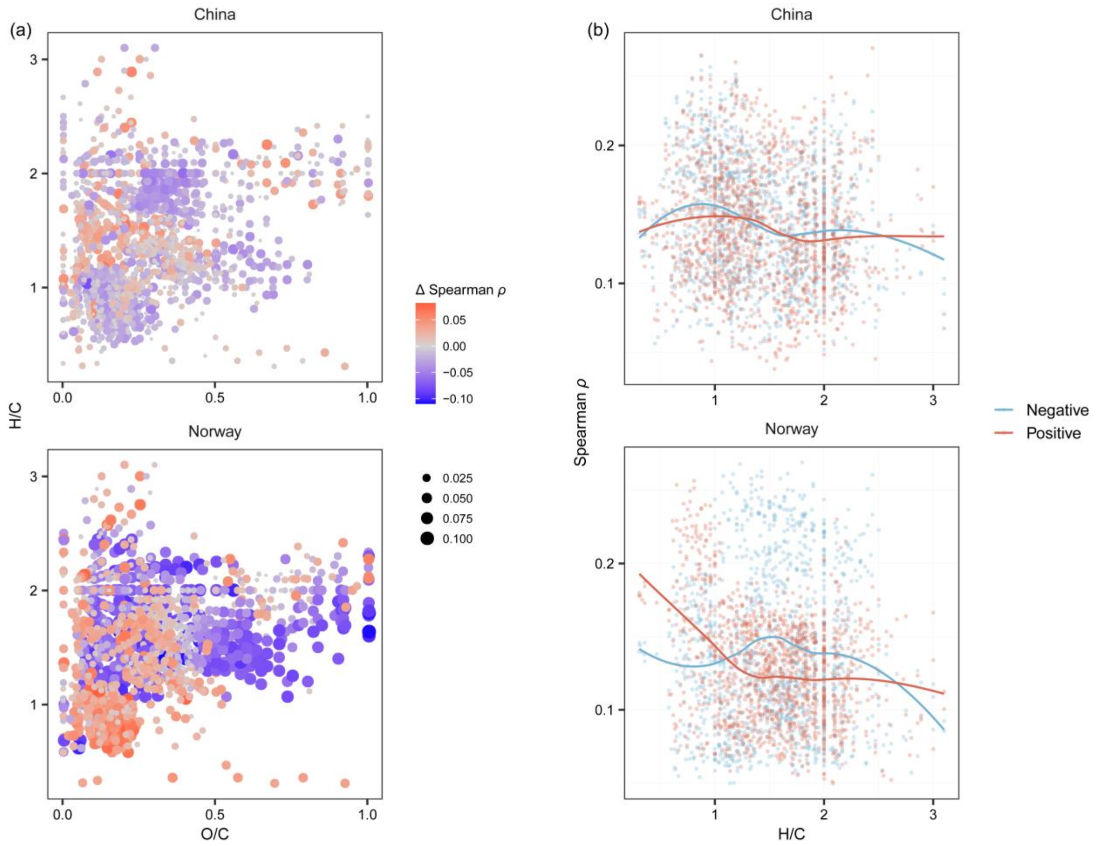


**Figure S15**. Correlations between DOM and bacteria regarding molecular traits. (a) Molecular formulae correlating with bacterial OTUs in China and Norway using Spearman's rank correlation ($\rho$). Each molecule is colored by difference in the absolute Spearman $\rho$ ($\Delta\rho$). The difference for each molecule was calculated by subtracting the mean absolute $\rho$ value of the negative correlations across all bacterial OTUs from that of the negative correlations, and the absolute value of $\rho$ difference is indicated by the dot size. (b) The patterns of absolute $\rho$ values of positive and negative correlations along the gradient of H/C ratio visualized with loess regression models.

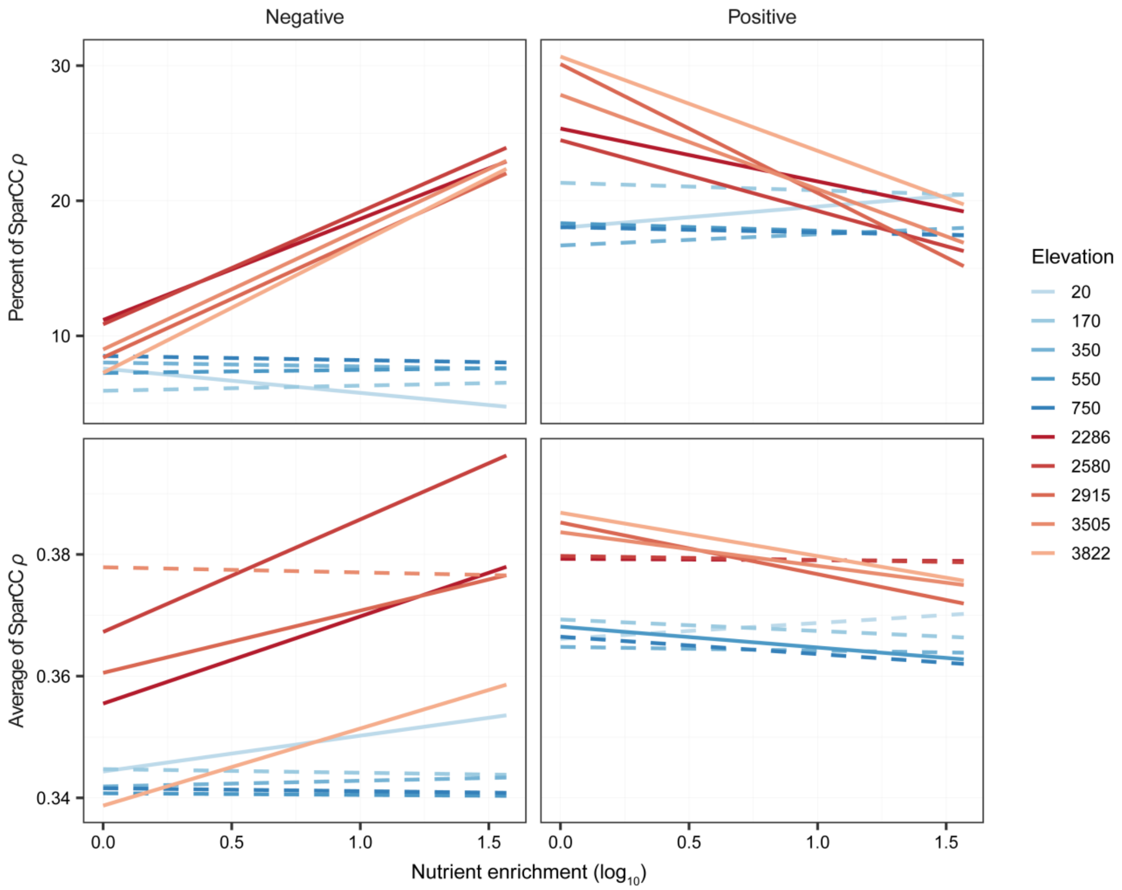


**Figure S16**. The effects of nutrient enrichment on weighted means of indices of DOM-microbe bipartite networks. The network indices include the percent and average of strong correlations (|SparCC $\rho$| ≥ 0.3) of negative or positive networks. We plotted these indices against the nutrient gradient of nitrate for both negative (left panel) and positive (right panel) networks for each elevation in China (red lines) or Norway (blue lines), and their relationships are indicated by solid ($P \leq 0.05$) or dotted ($P > 0.05$) lines using linear models.

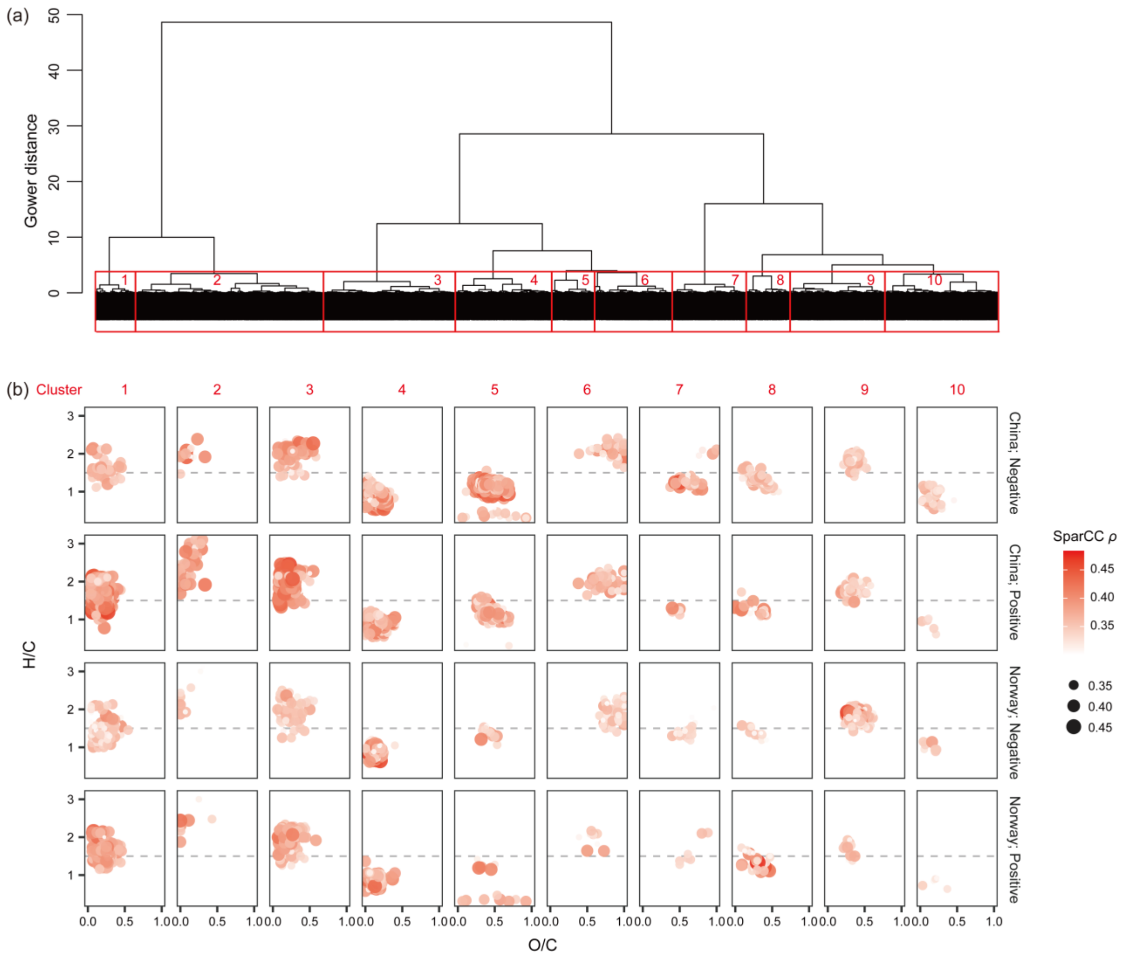


**Figure S17**. Correlations between DOM and bacteria regarding molecular traits. (a) Cluster analysis identified ten molecular sub-mixtures based on 16 molecular traits (Table S1). (b) Location of the ten clusters in Van Krevelen space with colour- and size-coded correlations between molecule-specific intensities and the relative abundance of bacterial genera using SparCC (Sparse Correlations for Compositional data). For each molecule, we showed the mean absolute SparCC $\rho$ values of negative or positive correlations across all bacterial OTUs. We considered only strong correlations ($|\text{SparCC } \rho| \geq 0.3$).

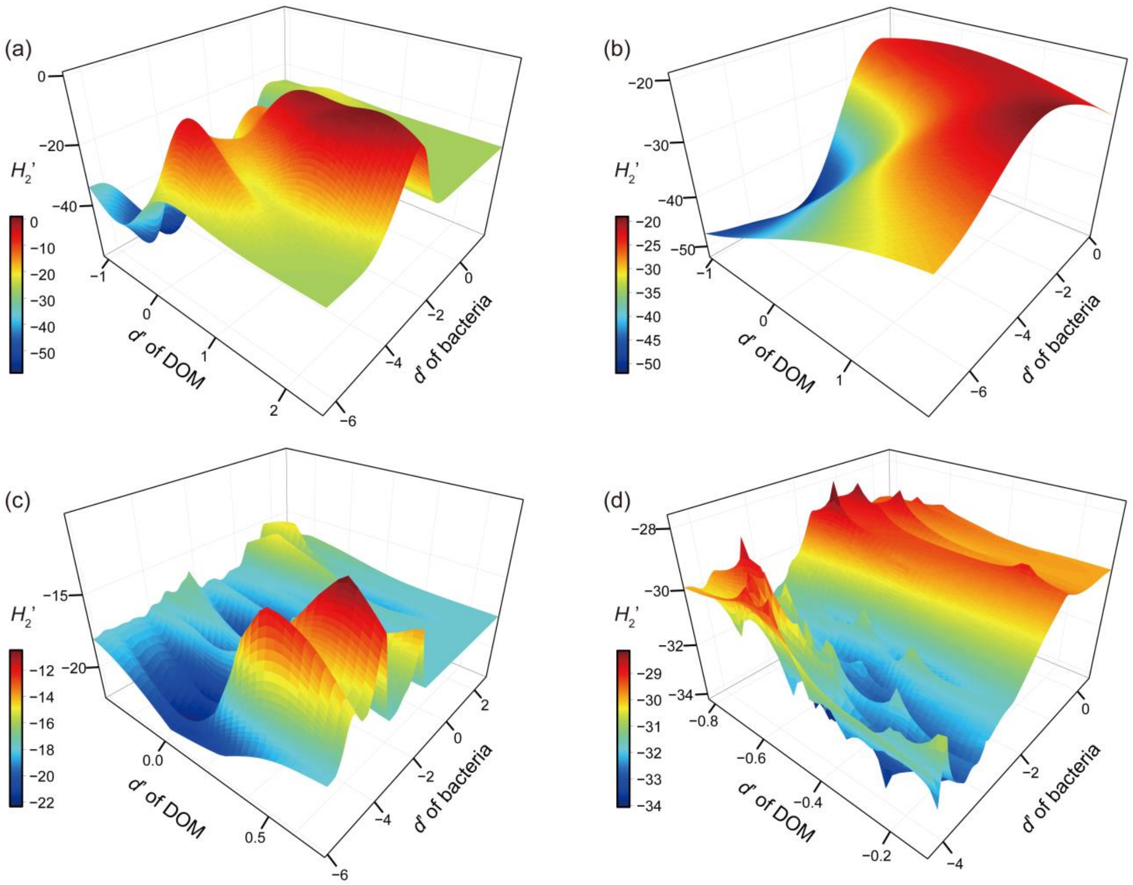


**Figure S18**. The relationships among the three specialization indices of DOM-microbe bipartite networks: network-level specialization $H_2$', the weighted means of specialization *d*' for DOM molecules and bacterial genera. These relationships were visualized for negative (a, c) and positive (b, d) networks in China (a, b) and Norway (c, d). The colours indicate the values of $H_2$'.

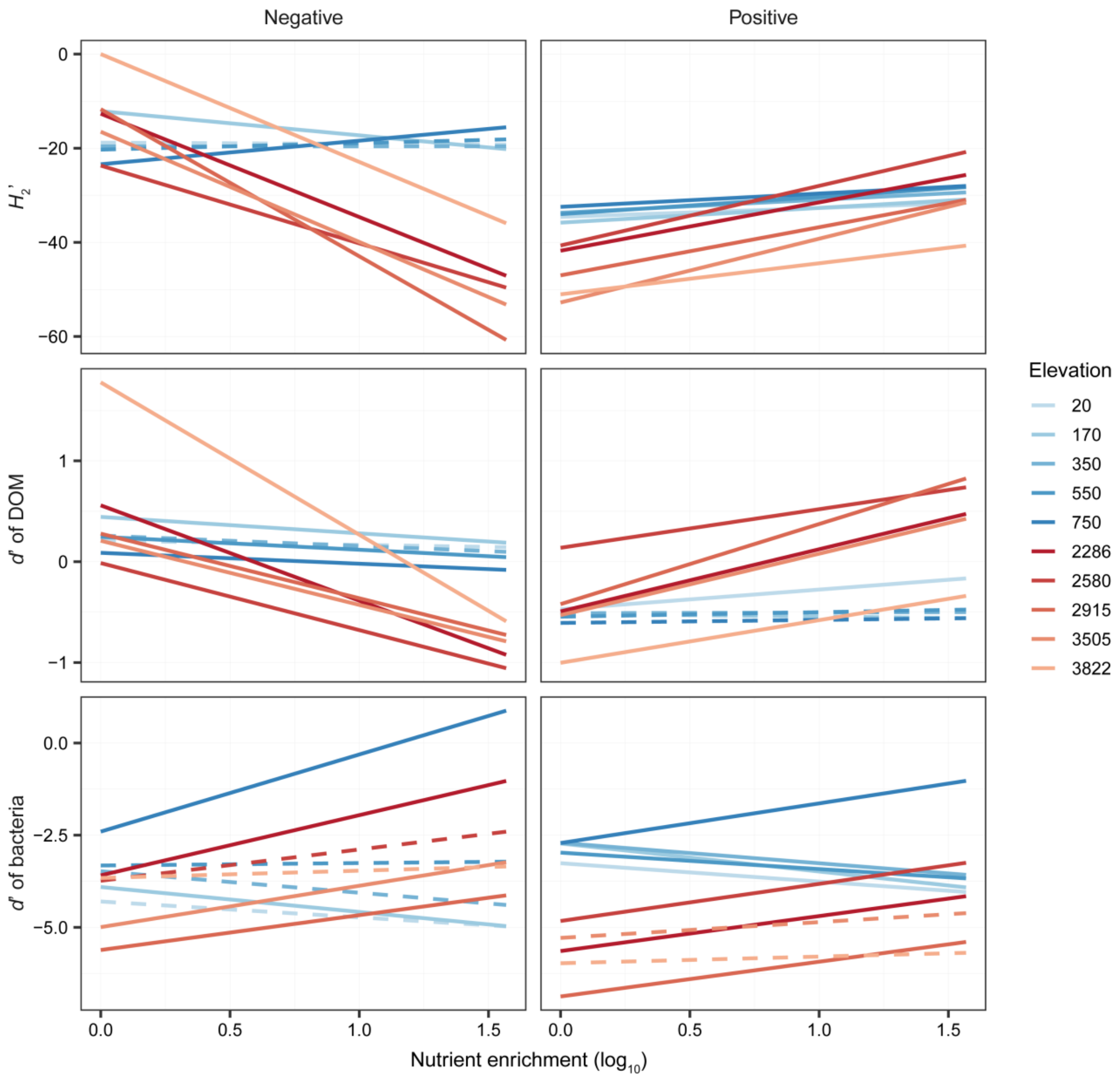


**Figure S19**. The effects of nutrient enrichment on specialization indices of DOM-microbe bipartite networks. The specialization indices include network-level specialization $H_2$', and the weighted means of specialization *d*' for DOM molecules and bacterial genera. We plotted these indices against the nutrient gradient of nitrate for both negative (left panel) and positive (right panel) networks for each elevation in China (red lines) or Norway (blue lines), and their relationships are indicated by solid ($P \leq 0.05$) or dotted ($P > 0.05$) lines using linear models.

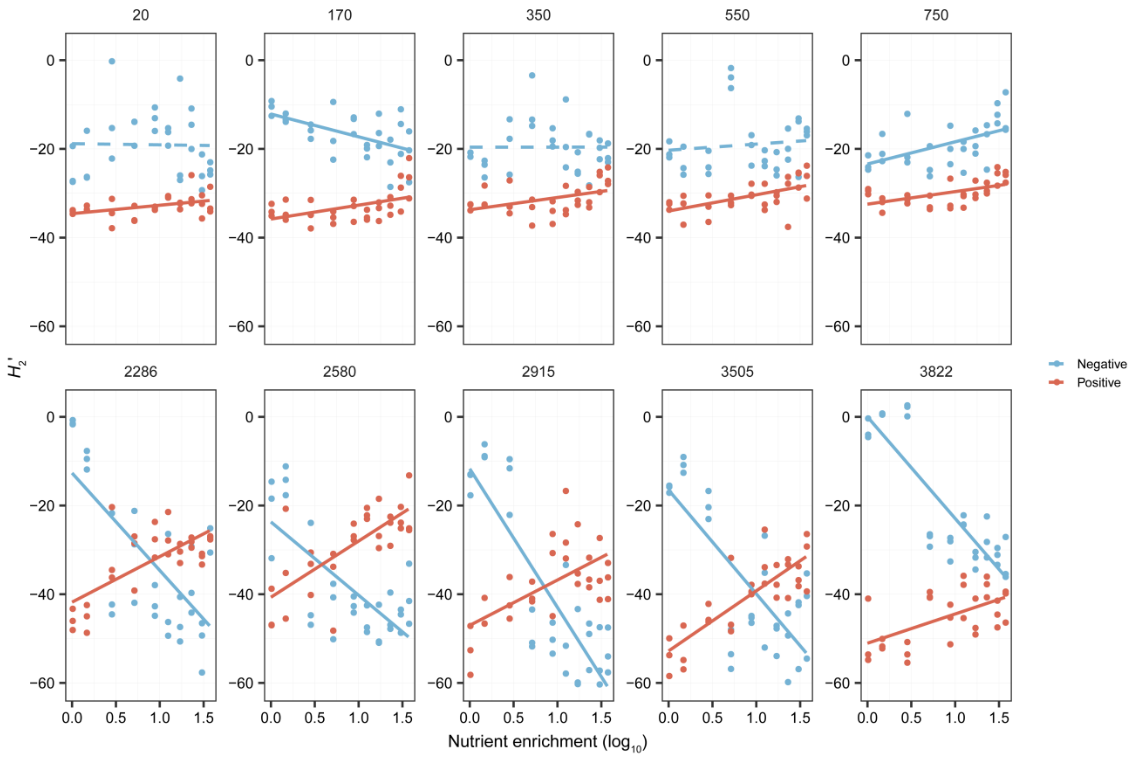


**Figure S20**. The effect of nutrient enrichment on the specialization $H_2$' of DOM-microbe bipartite networks. We plotted the $H_2$' against the nutrient gradient of nitrate for both negative (blue lines) and positive (red lines) networks for each elevation in China or Norway, and their relationships are indicated by solid ($P \leq 0.05$) or dotted ($P > 0.05$) lines using linear models.

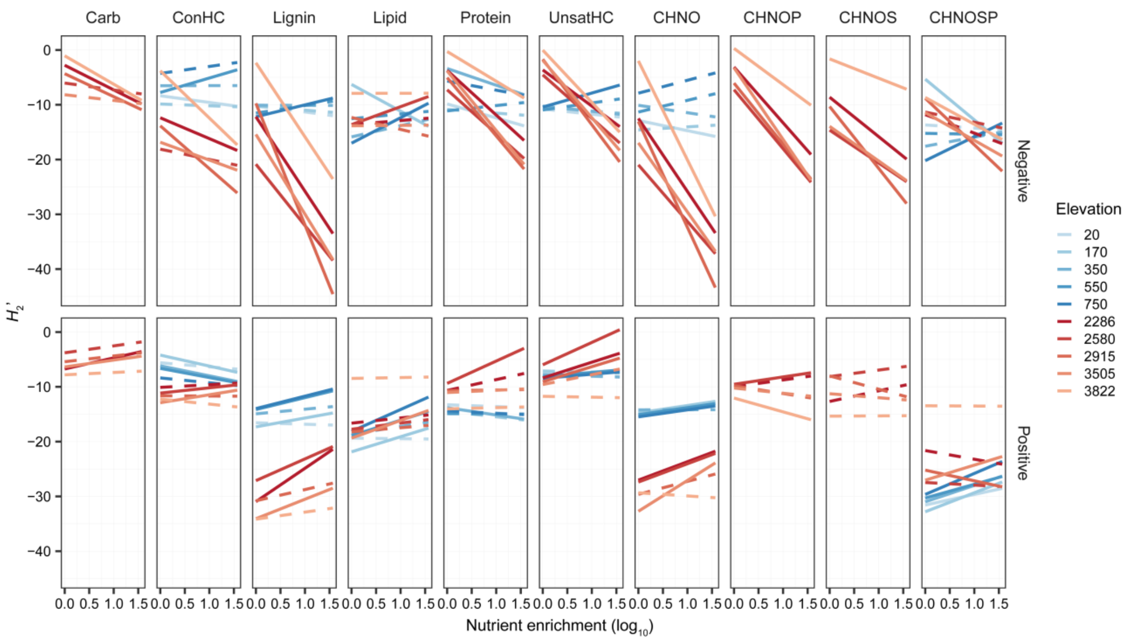


**Figure S21**. The effects of nutrient enrichment on the specialization $H_2$' of DOM-microbe associations for all formulae and subsets of formulae within the category of compound classes or elemental combinations. We plotted the $H_2$' against the nutrient gradient of nitrate for both negative (upper panel) and positive (lower panel) networks for each elevation in China (red lines) or Norway (blue lines), and their relationships are indicated by solid ($P \leq 0.05$) or dotted ($P > 0.05$) lines using linear models.

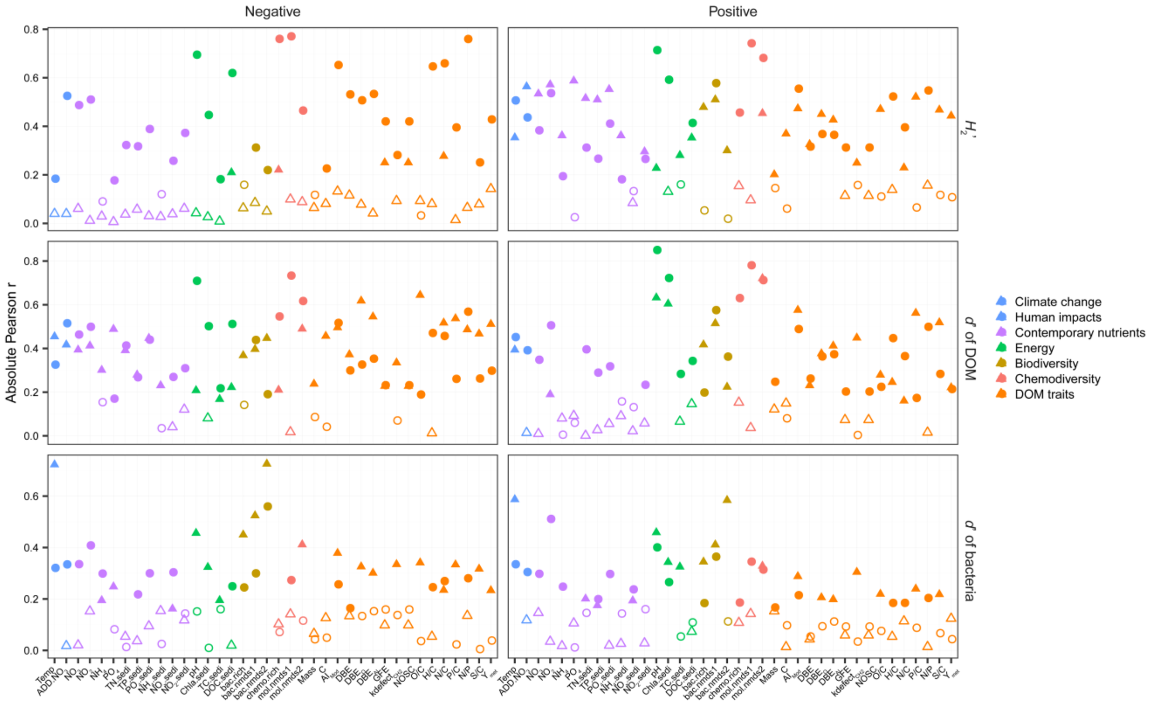


**Figure S22**. The relative influence of explanatory variables on the specialization $H_2$' of negative and positive DOM-microbe bipartite networks using Pearson correlation analysis. The specialization indices include the network-level specialization $H_2$' (upper panel), and the weighted means of specialization *d*' for DOM molecules (middle panel) and bacterial genera (lower panel). Each circle and triangle are the absolute values of Pearson r for individual explanatory variable in China and Norway, respectively. Solid and open circles or triangles indicate the significant ($P \leq 0.05$) and non-significant ($P > 0.05$) Pearson r, respectively. The details of abbreviations of explanatory variables are available in Table S1.

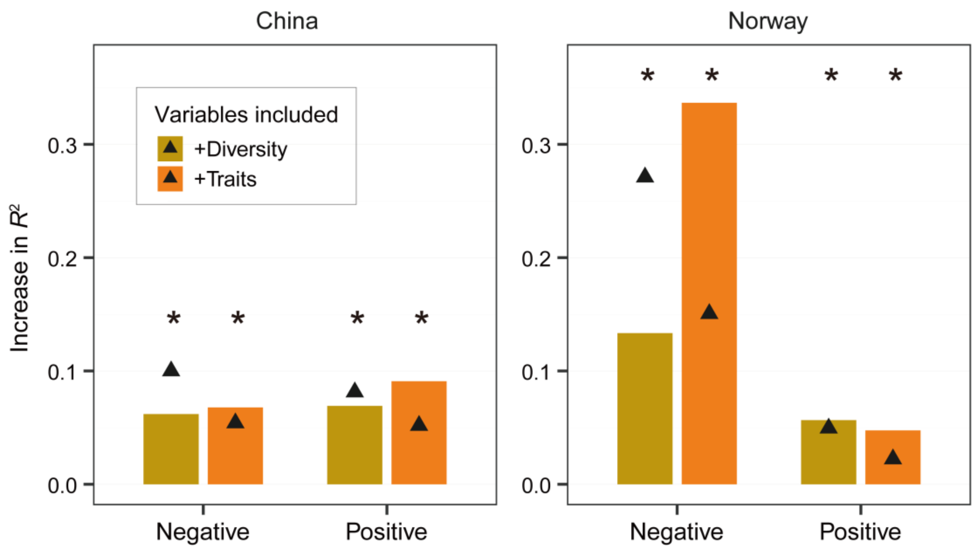


**Figure S23**. Relative importance of diversity (i.e., bacterial diversity and chemodiversity, brown bars) and molecular traits (orange bars) in explaining specialization $H_2$’ of negative and positive bipartite networks. We calculated improvements in explained variances ($R^2$) relative to the linear models without diversity and molecular traits (i.e., using only variables associated with environments and energy supply) in China (left panel) and Norway (right panel). Asterisks denote statistically significant improvements at $P \leq 0.05$. Triangles show the net effects estimated by random forests.

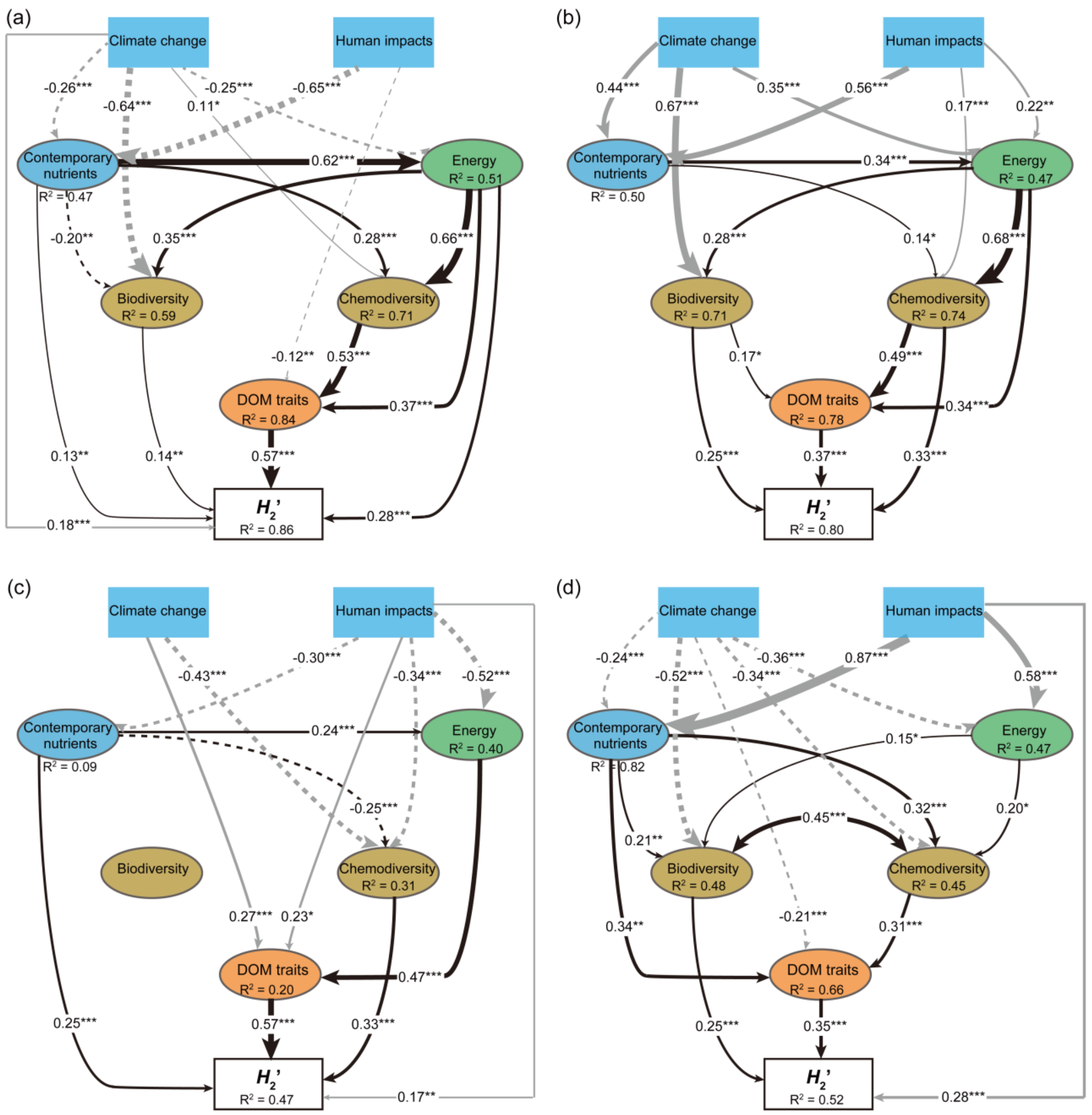


**Figure S24**. Structural equation models to explain specialization of DOM-microbe networks. Best-fitting models illustrate the effects of predictor variables on the $H_2$’ of negative (a, c) and positive (b, d) bipartite networks in China (a-b) or Norway (c-d). Predictor variables were grouped by climate change, human impacts, contemporary nutrients, energy supply, biodiversity, chemodiversity and DOM traits, and described in detail in Table S2. $R^2$ denotes the proportion of variance explained for the endogenous variables. Dotted and solid arrows indicate statistically significantly negative and positive ($^{***}$, $P \leq 0.001$; $^{**}$, $P \leq 0.01$; $^{*}$, $P \leq 0.05$) relationships, respectively. Grey or black arrows indicate the hypothesized relationships among the exogenous or endogenous variables and $H_2$', respectively. Arrow widths and accompanying numbers are the relative effects (that is, standardized path coefficients) of modeled relationships. Composite and observed variables are indicated in ovals and rectangles, respectively. Details of model fit are summarized in Table S3.

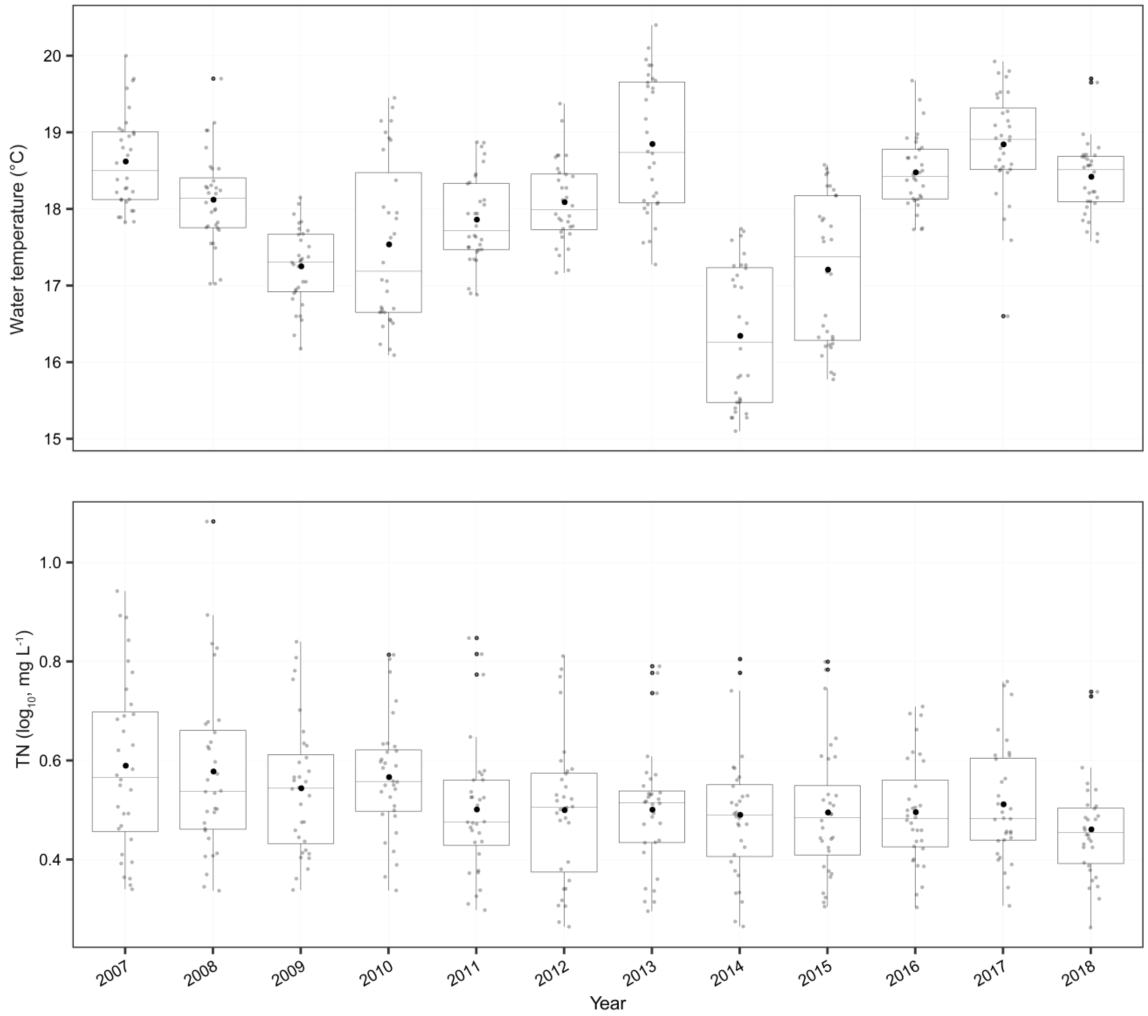


**Figure S25**. Water temperature and total nitrogen (TN) during 2007-2018 in Taihu Lake. The grey dots indicate water temperature and TN for individual sampling sites (Fig. S27a) and black dots are the mean values for each year.

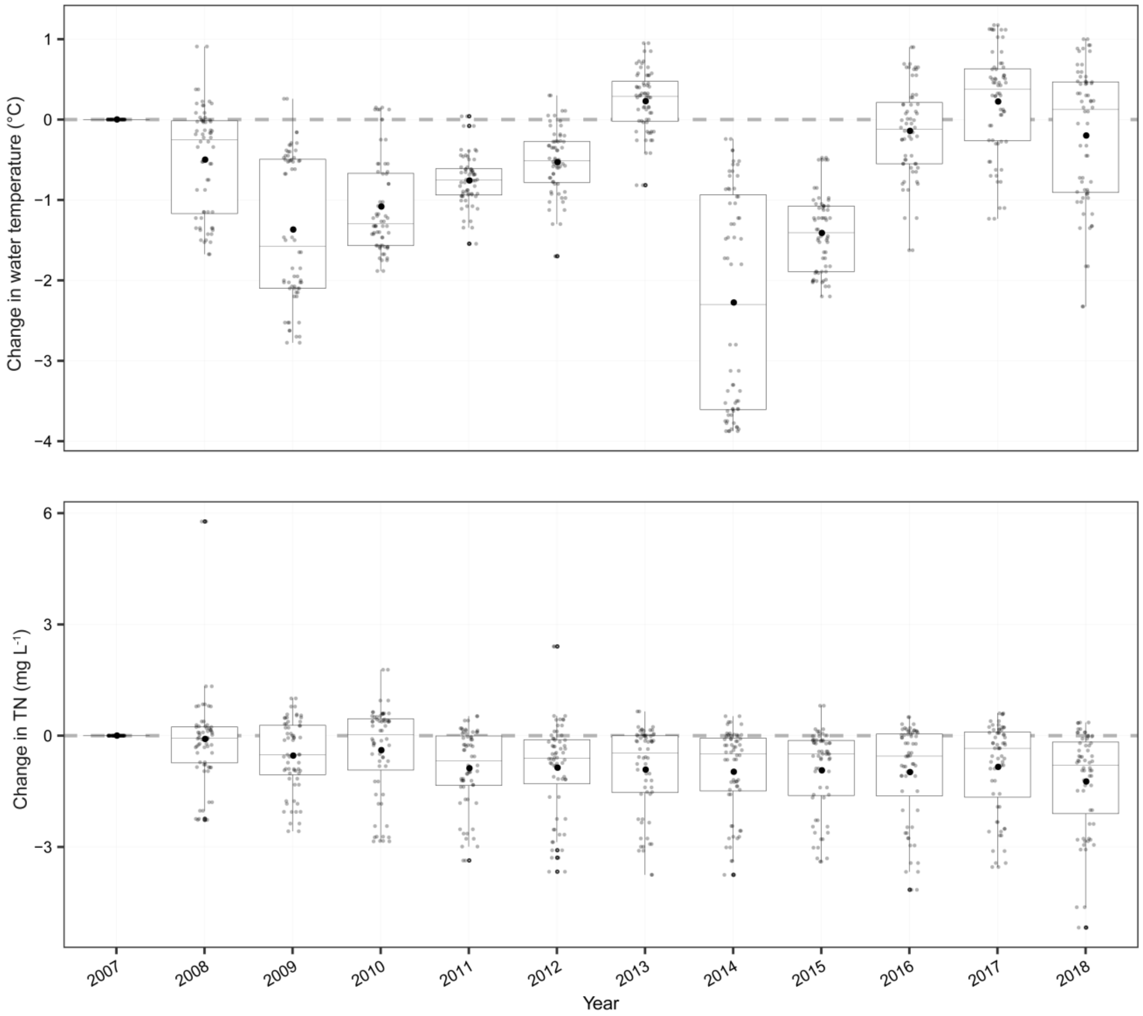


**Figure S26**. Changes in water temperature and total nitrogen (TN) from 2007 to 2018 in Taihu Lake. The temporal changes were calculated using 2007 as a baseline to which all predictions were compared. The grey dots indicate changes for individual sampling sites (Fig. S27a) and black dots are the mean values for each year.

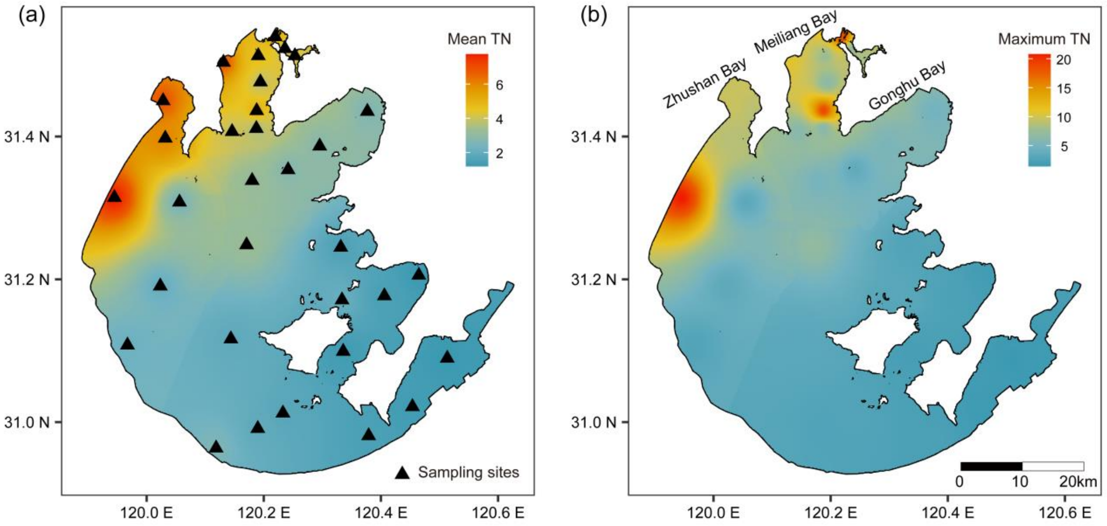


**Figure S27**. The distribution of mean (a) and maximum (b) total nitrogen (TN) concentrations (mg $L^{-1}$) in 2007 across the Taihu Lake. Triangles indicate 32 sampling sites.

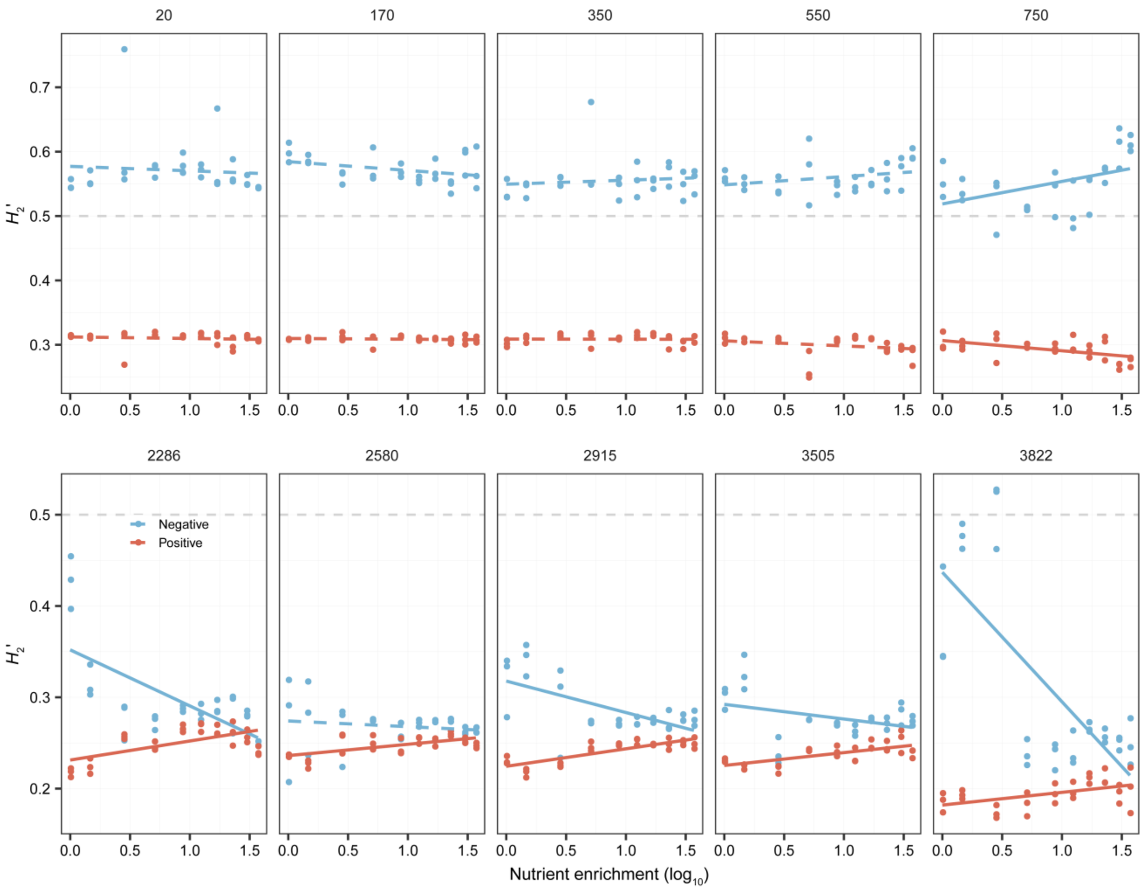


**Figure S28**. The effects of nutrient enrichment on the observed values of specialization $H_2$' of DOM-microbe associations. We plotted the $H_2$' against the nutrient gradient of nitrate for both negative (blue lines) and positive (red lines) networks for each elevation in China or Norway, and their relationships are indicated by solid ($P \leq 0.05$) or dotted ($P > 0.05$) lines using linear models. The horizontal dashed lines indicate more specialized associations with a $H_2$' value above 0.5.
